# Unveiling the role of SRY in male-biased cancers: Insights into the molecular basis of sex disparities in high-grade glioma and melanoma

**DOI:** 10.1101/2023.07.14.548747

**Authors:** Gabriela D A Guardia, Rafael Loch Batista, Luiz O. Penalva, Pedro A F Galante

## Abstract

Sex disparities have been observed in many tumor types affecting non-reproductive organs. Typically, the incidence and mortality rates of such cancers are higher in men. Although differences in lifestyle and environmental exposures are known contributors, knowledge of the molecular mechanisms driving sexual dimorphism in tumor development and therapy response remains limited. To address this question, we comprehensively studied the sex-determining region Y (SRY) gene, a male-specific gene that is critical in development. First, we screened 2,448 samples from 11 cancer types to identify those with a higher incidence in men and increased expression of SRY. In cases of high-grade glioma and melanoma, men with tumors exhibiting high SRY expression had a worse prognosis. Our results suggest that SRY target genes show altered expression when SRY is overexpressed. These gene sets are linked to cell growth, epithelial-mesenchymal transition, inflammation, and repression of tumor suppressor pathways. In summary, we present the first comprehensive investigation of SRY expression and its association with clinical outcomes in men with high-grade glioma and melanoma. Our results shed light on the molecular basis for sex disparities and lay the foundation for investigation of various target genes and novel cancer treatments in men with high-grade glioma and melanoma.

## INTRODUCTION

Sexual dimorphism refers not only to the differences between males and females in physiologic and behavioral characteristics, but also to disease susceptibility and treatment outcomes (Clocchiatti et al., 2016). Many cancers exhibit sexual dimorphism in terms of incidence, prognosis, and treatment response (Siegel et al., 2023). For organs with non-reproductive functions, cancer incidence and mortality are usually higher in men (Cook et al., 2009; Dorak and Karpuzoglu, 2012). While lung cancer, bladder cancer, liver hepatocarcinoma, esophageal cancer, and high-grade gliomas are more common in men, gallbladder, anal, and thyroid cancers are more common in women (Cook et al., 2009; Dorak and Karpuzoglu, 2012). Overall, the underlying reasons for sexual dimorphism in cancer are poorly understood, in particular the genetic and hormonal contributions involved (Dorak and Karpuzoglu, 2012). More knowledge is required for developing tailored prevention and treatment strategies that consider the unique biology and health needs of each gender.

Among the male-specific genetic factors potentially related to cancers that are more common in men, the male-specific region in chromosome Y has attracted interest. This region is 54 MB in size and contains roughly 80 protein-coding genes (Skaletsky et al., 2003). The sex-determining region Y (SRY) gene is a transcription factor that stands out among these genes. SRY is the crucial gene that initiates male sex determination in mammals, including humans (Hughes and Page, 2016; Kashimada and Koopman, 2010). Elucidating the role of SRY has greatly advanced our understanding of the molecular basis of sexual differentiation. SRY codes for a protein that regulates the expression of other genes (e.g., SOX9 and SF1) involved in the differentiation of the bipotential gonad into a male gonad, a primordial step in male sexual development (Mäkelä et al., 2019). In humans, mutations or abnormalities in SRY expression can lead to disorders of sexual development (Lucas-Herald and Bashamboo, 2014). For example, individuals with XY chromosomes but a mutation in the SRY gene may develop as phenotypic females, while individuals with XX chromosomes but the SRY gene translocated to the X chromosome may develop as phenotypic males (Ahmed et al., 2022). Moreover, growing evidence suggests that the SRY may be involved in the development of certain types of cancer, particularly those that are more common in men (Liu et al., 2017; Xue et al., 2015),(Correa et al., 2014; Murakami et al., 2015). SRY expression levels have already been correlated with cancer progression and prognosis in patients with hepatocellular carcinoma (Liu et al., 2017; Xue et al., 2015) and in a few other tumors, such as prostate cancer (Correa et al., 2014; Murakami et al., 2015).

Skin cutaneous melanoma (SKCM) is a type of cancer that arises from the pigment-producing skin cells known as melanocytes. SKCM is one of the top 10 most common cancers (Saginala et al., 2021), with exposure to ultraviolet radiation being its primary cause (Lopes et al., 2021). Additionally, SKCM is the most aggressive and deadly form of skin cancer, and is responsible for most skin cancer deaths (Hartman and Lin, 2019). Early detection of melanoma is associated with a favorable prognosis (Petrie et al., 2019; Rabbie et al., 2019). However, if the melanoma has metastasized, the five-year survival rate drops to around 27%. Melanoma treatment options include surgery, radiation therapy, chemotherapy, and newer targeted and immunotherapy treatments (Babacan and Eroglu, 2020; Luke et al., 2017). Melanoma incidence rates differ between genders, with a higher incidence (up to 50% more in some countries) observed in men (Saginala et al., 2021).

High-grade gliomas (HGG), which include glioblastoma, anaplastic astrocytoma, and anaplastic oligodendroglioma, originate from the supportive neuroglial cells of the central nervous system (Wen and Kesari, 2008). HGG can cause cognitive impairment, personality changes, and seizure disorders, and treatment requires complex multidisciplinary care. Despite intensive efforts to develop new therapies for HGG, treatment options remain limited, and prognosis is still poor (Stupp et al., 2014). Moreover, HGG are more prevalent in men than in women, with a male-to-female ratio of approximately 3:2 (Ostrom et al., 2022). The reasons for this difference in incidence are not fully understood, but related, in potential, to hormonal or genetic factors (Yang et al., 2019).

Here, we investigated the role of the SRY gene in tumors that are more common in men (in non-reproductive organs). We used a comprehensive strategy to assess SRY expression in bulk and single-cell datasets along with clinical and survival information, followed by a large-scale functional analysis in 11 cancer types. Our results identified two highly aggressive tumor types with higher incidence in men, high-grade glioma and melanoma, in which increased SRY expression was associated with a worse prognosis. Moreover, our results linked high SRY expression to specific oncogenic pathways.

## RESULTS

### SRY is differentially expressed in aggressive tumors that are more common in men

We employed an integrative approach to assess SRY expression levels and its potential regulatory impact on tumors with higher incidence in men. We used gene expression (bulk and single-cell) as well as clinical data to investigate SRY expression patterns and its association with patients’ prognosis and oncogenic pathways in 2,448 tumor samples from 11 cancer types (Figure 1A) (see Methods for details). We started analyzing the gender distribution of cancer patients from The Cancer Genome Atlas (TCGA) and identified several tumor types that were at least 20% more frequent in men (Figure 1B and Supplementary Table S1). As expected, based on data from The United States and Europe, a higher number of cases was observed among men in TCGA for esophageal carcinoma, lung squamous cell carcinoma, bladder urothelial carcinoma, kidney carcinoma, liver hepatocellular carcinoma, skin cutaneous melanoma, and high-grade (III-IV) glioma (Dyba et al., 2021; Siegel et al., 2023).

**Figure 1.**
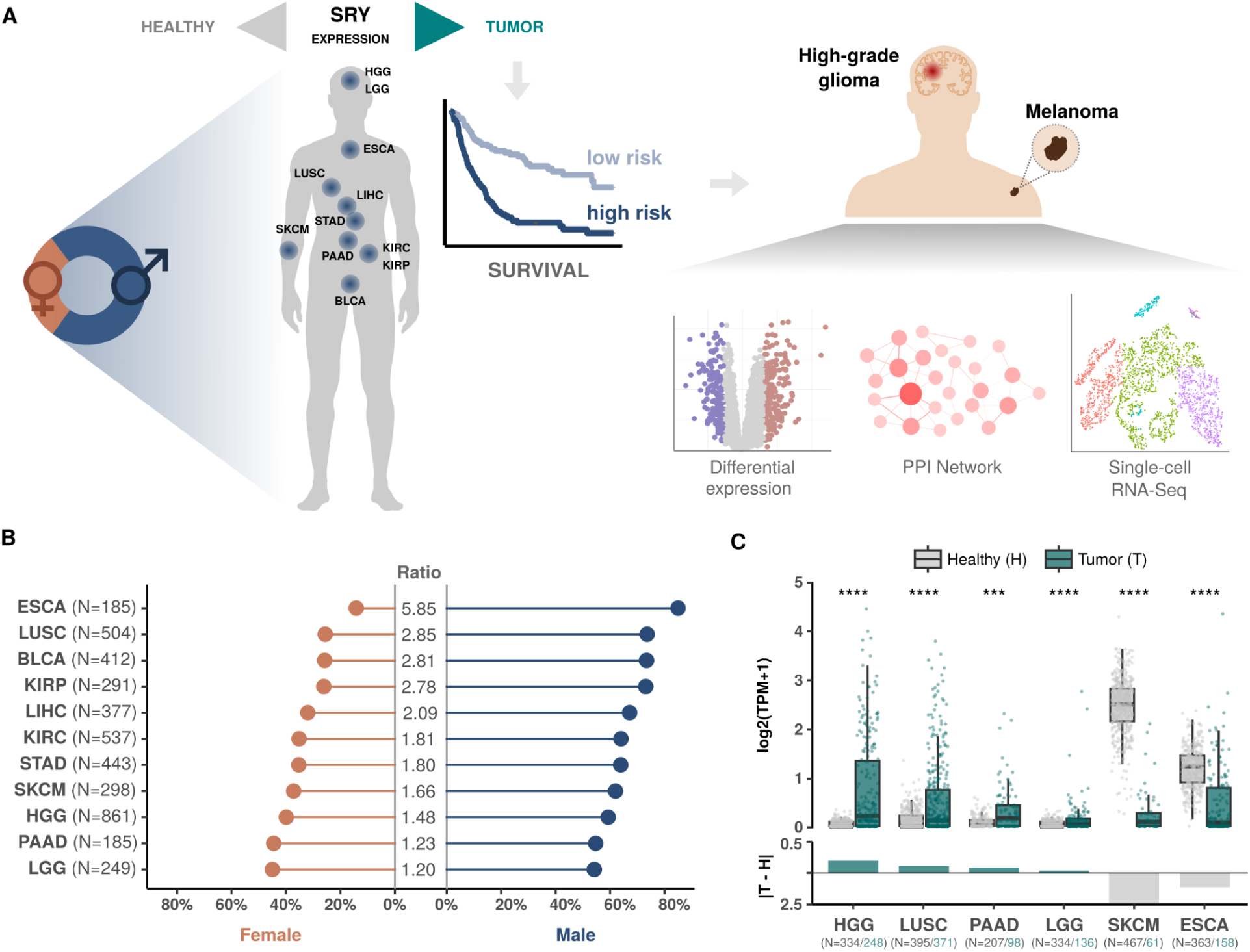
Identifying SRY differential expression in cancer types that are more prevalent among men. **A)** Schematic view of the approach used to investigate SRY gene expression in male cancer patients. **B)** Tumors that are more common in men (based on the number of samples) in the TCGA cohort. **C)** Comparison of SRY expression levels (TPM) in tumors from male patients from TCGA and healthy tissues from GTEx (Wilcoxon tests; **** p-value ≤ 0.0001, *** p-value ≤ 0.001). Abbreviations: ESCA, esophageal carcinoma; LUSC, lung squamous cell carcinoma; BLCA, bladder urothelial carcinoma; KIRP, kidney renal papillary cell carcinoma; LIHC, liver hepatocellular carcinoma; KIRC, kidney renal clear cell carcinoma; STAD, stomach adenocarcinoma; SKCM, skin cutaneous melanoma; HGG, high-grade glioma (grades III-IV); PAAD, pancreatic adenocarcinoma; and LGG, low-grade glioma (grade II).

To assess the contribution of SRY expression levels in these types of tumors, we first examined differences between tumors and normal tissues. Increased SRY expression was observed in high-grade and low-grade gliomas, lung squamous cell carcinoma, and pancreatic adenocarcinoma, while skin cutaneous melanoma and esophageal cancer exhibited SRY underexpression (p-value < 0.001; Wilcoxon test) (Figure 1C).

### SRY is preferentially expressed in malignant glioma cells, and its high expression is associated with worse prognosis in men with HGG

We investigated the prognostic value of SRY in the six cancer types with higher frequency in men (Figure 1C). We used log-rank tests to compare progression-free survival rates of 1,068 patients stratified in two groups based on their SRY expression levels: high (transcripts per million [TPM] ≥ 0.1) or low (TPM < 0.1). Considering all tumors, only HGG and SKCM showed significant associations (log rank test p-values < 0.05) between SRY expression and patients’ prognoses (Supplementary Figure S1 and Figure 2A). In both types of cancer, high SRY expression was associated with a worse prognosis.

**Figure 2.**
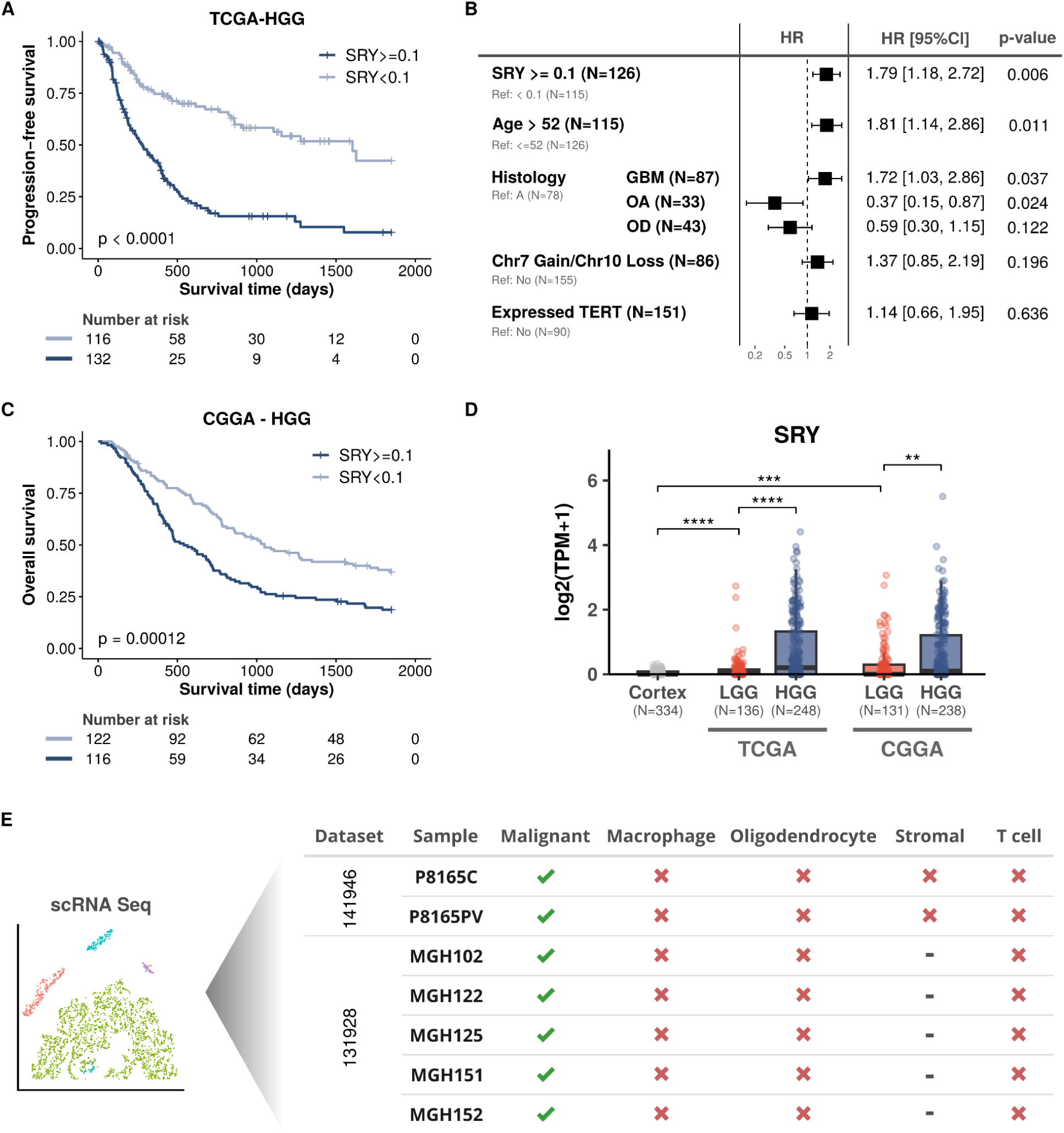
SRY is preferentially expressed in malignant glioma cells and its overexpression is associated with a worse prognosis in men with HGG. **A)** Progression-free survival in the TCGA cohort of men with HGG and high (TPM ≥ 0.1) or low (TPM < 0.1) SRY expression. **B)** Multivariate Cox regression analysis considering several variables among men with HGG: SRY expression, age (median: 52 years), gain of chromosome 7 and loss of chromosome 10, TERT expression status and four glioma histologies: GBM, glioblastoma; A, astrocytoma; OA, oligoastrocytoma; and OD, oligodendroglioma. **C)** Overall survival for CGGA cohort of men with HGG and high (TPM ≥ 0.1) or low (TPM < 0.1) SRY expression. **D)** Expression levels of SRY in healthy (frontal) cortex samples from male individuals (GTEx cohort) and in tumor samples from men with LGG or HGG from the TCGA and CGGA cohorts (Wilcoxon tests; **** p-value ≤ 0.0001, *** p-value ≤ 0.001, ** p-value ≤ 0.01); **E)** Results of single-cell RNA sequencing from seven HGG samples from two studies (acessions GSE141946 and GSE131928) showing SRY expression in cancer cells (malignant; in green), but not in other cells identified.

Deeper investigation of SRY in HGG patients from the TCGA (Supplementary Table S2) determined that 50% of patients with high SRY expression had recurrence in up to 300 days, while patients with low SRY expression had later recurrence (over 1,500 days) (Figure 2A). Next, we performed multivariate Cox regression to estimate the associations between SRY expression and progression-free survival in HGG together with relevant clinical information. High SRY expression emerged as a pivotal factor in disease recurrence (hazard ratio of 1.79 [95% CI, 1.18 to 2.72]; p-value = 0.006), Figure 2B).

To confirm the prognostic value of SRY in an independent set of gliomas, we used gene expression and clinical information from 238 men with HGG in the Chinese Glioma Genome Atlas (CGGA) (Supplementary Table S3). In agreement with our findings in the TCGA cohort, high expression of SRY (TPM > 0.1) was associated with a worse prognosis (Figure 2C). While 50% of HGG patients with a high SRY expression died within 500 days, patients with low SRY expression had survival times of ∼1,000 days (Figure 2C). To further investigate the relationship between SRY expression and glioma aggressiveness, we assessed SRY expression in low-grade gliomas (LGG), which are less aggressive and have a better prognosis than HGG. SRY expression was significantly lower in LGG (and even lower in normal cortex) compared to HGG (p-value < 0.001; Wilcoxon rank test) (Figure 2D). This result is consistent with our previous findings showing that gliomas with higher SRY expression are more aggressive (Figure 2A-C).

Finally, we used single-cell gene expression data to assess SRY expression in cells from HGG samples (Supplementary Figures S2-S3). SRY was exclusively expressed in malignant tumor cells and not in other cell types such as macrophages, oligodendrocytes, stromal cells, or T-cells in the tumor microenvironment (Figure 2E). Taken together, our results indicate that SRY is predominantly expressed in malignant HGG cells and that its overexpression is significantly associated with a worse prognosis for patients with HGG.

### SRY target genes are associated with essential cancer aggressiveness pathways

Previous reports have shown that SRY regulates genes associated with cancer development and progression (Liu et al., 2017; Murakami et al., 2014). To evaluate SRY’s downstream impact on tumor progression, we first selected its target genes identified in chromatin immunoprecipitation (ChIP-Chip) experiments (Li et al., 2014) (Supplementary Table S4). Differential expression profiling between tumors with high (TPM ≥ 0.1) or low (TMP < 0.1) SRY expression (see Methods) identified 310 SRY target genes (Figure 3A; Supplementary Table S5). Next, pathway enrichment analysis of this set of SRY targets identified three enriched pathways: *epithelial mesenchymal transition* (associated with metastatic expansion, generation of tumor cells with stem cell properties, and resistance to cancer treatment); *proteoglycans in cancer* (associated with cancer cell adhesion and migration); and *NF Kappa B signaling* (associated with proliferation, cell survival, growth, and angiogenesis), among others (Figure 3B).

**Figure 3.**
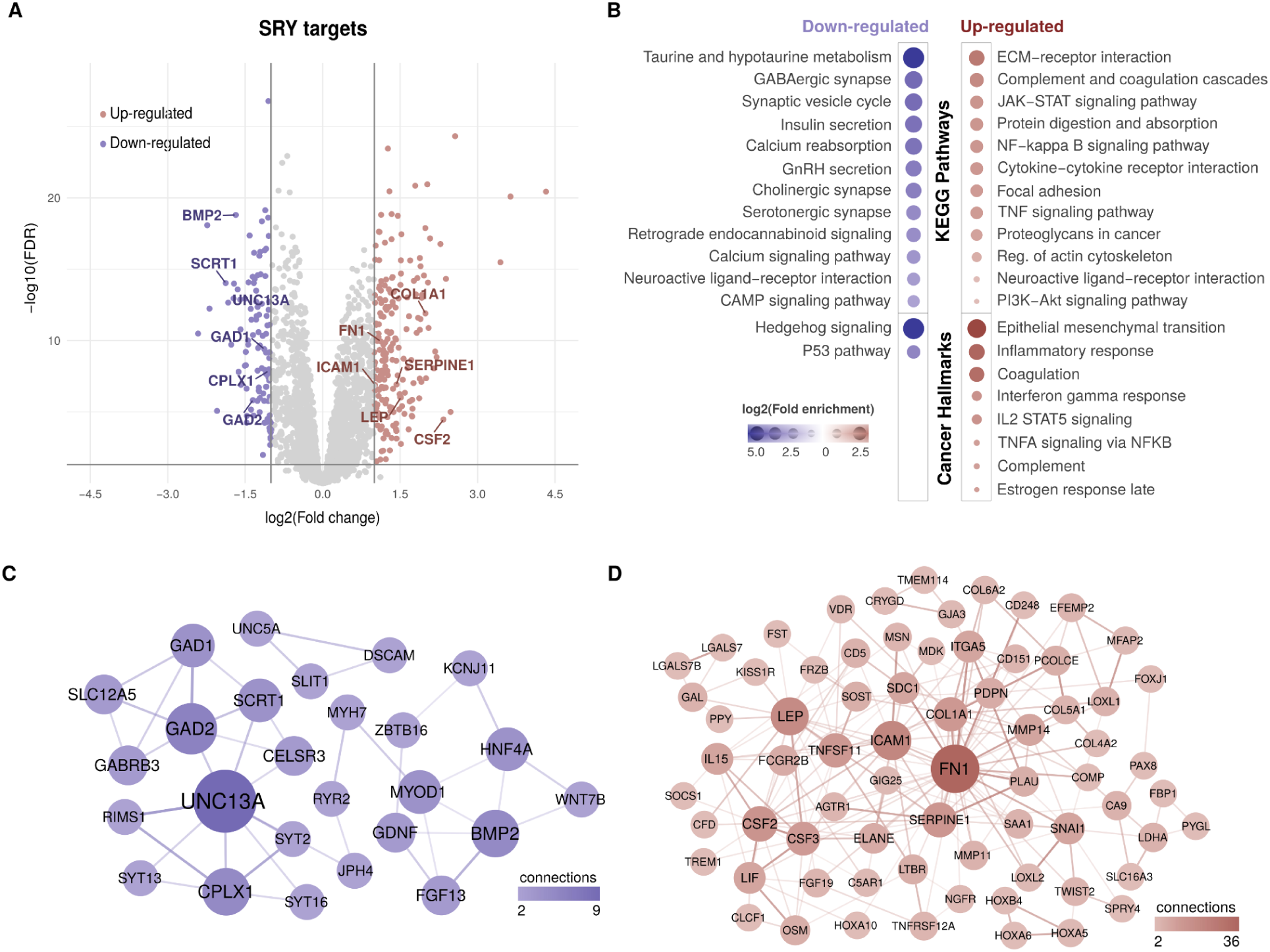
HGG samples with high expression of SRY show target genes associated with activation of cancer aggressiveness pathways and the suppression of tumor suppressor features. **A)** Differentially expressed SRY target genes between HGG samples showing higher (≥ 0.1 TPM) versus lower SRY expression (< 0.1 TPM). **B)** Functional enrichment analysis (KEGG pathways and MSigDB Hallmarks) of SRY target genes up- and down-regulated in HGG samples. **C)** Protein-protein interaction networks of SRY target genes that were down-regulated in samples with higher SRY expression. **D)** Protein-protein interaction networks of SRY target genes that were upregulated in samples with higher SRY expression.

This pathway analysis also identified the *Estrogen response late* pathway as enriched. While estrogen is typically associated with female sex hormones, it is also produced in men, albeit at lower levels. Additionally, abnormal activation of the estrogen response late pathway has been associated with an increased risk of developing cancers in men (Nelles et al., 2011). On the other hand, the *P53 tumor suppressor pathway* was associated with downregulated target genes for SRY (Figure 3B).

To provide a comprehensive view of target genes differentially expressed in tumors with high SRY expression, we performed protein-protein interaction (PPI) analyses using these genes. The PPI network of downregulated SRY target genes preferentially contained proteins related to synthesis, release, and regulation of neurotransmitters (Figure 3C), essential functions of the nervous system commonly altered in HGG (Guardia et al., 2020). Furthermore, the majority of these genes (69.2%) have already been associated with cancer, particularly demonstrating reduced expression (61%), and with validated or putative tumor suppression functions (Supplementary Table S6). These findings are thus in agreement with our observation of their downregulation in HGG samples.

The PPI network of overexpressed SRY target genes revealed proteins linked to extracellular matrix remodeling, cell adhesion, migration, angiogenesis, tumor invasion, and metastasis, such as FN1 (Bouvier et al., 2022), ICAM1 (Bui et al., 2020), COL1A1 (Verginadis et al., 2022), PLAU (Kisling et al., 2021), and SERPINE1 (Simone et al., 2014) (Figure 3D). These functions are commonly associated with tumors that have a poor prognosis, in agreement with our previous results (Figure 2A-C) and our hypothesis that SRY expression is associated with tumor aggressiveness. Collectively, these results provide insights into the molecular mechanisms driving cancer aggressiveness, including activation of cell migration, tissue invasion, angiogenesis, and suppression of immune cell infiltration, which could explain why men with HGG and high SRY expression have a worse prognosis (with more advanced tumor progression and lower survival rates) than those with low SRY expression.

### High expression of SRY is associated with a worse prognosis in melanoma patients

High SRY expression (TPM ≥ 0.1) was also associated with a worse prognosis (log rank test p-value < 0.05) in samples from men with skin cutaneous melanoma (SKCM) in the TCGA (Supplementary Table S7). While half of those with high SRY expression had disease recurrence within 500 days, recurrence in half of the patients with lower SRY expression persisted up to 1,000 days (Figure 4A). To more accurately assess other variables potentially associated with tumor progression, we performed multivariate Cox regression analysis considering SRY expression together with other clinical information from these patients (age and tumor stage). Only tumors with high expression of SRY (TPM ≥ 0.1) were significantly associated with length of survival (p-value < 0.03; hazard ratio of 2.77 [95% CI, 1.9 to 7.02]) (Figure 4B). Our data suggest that SRY was a major factor contributing to tumor recurrence in this cohort of men with melanoma.

**Figure 4.**
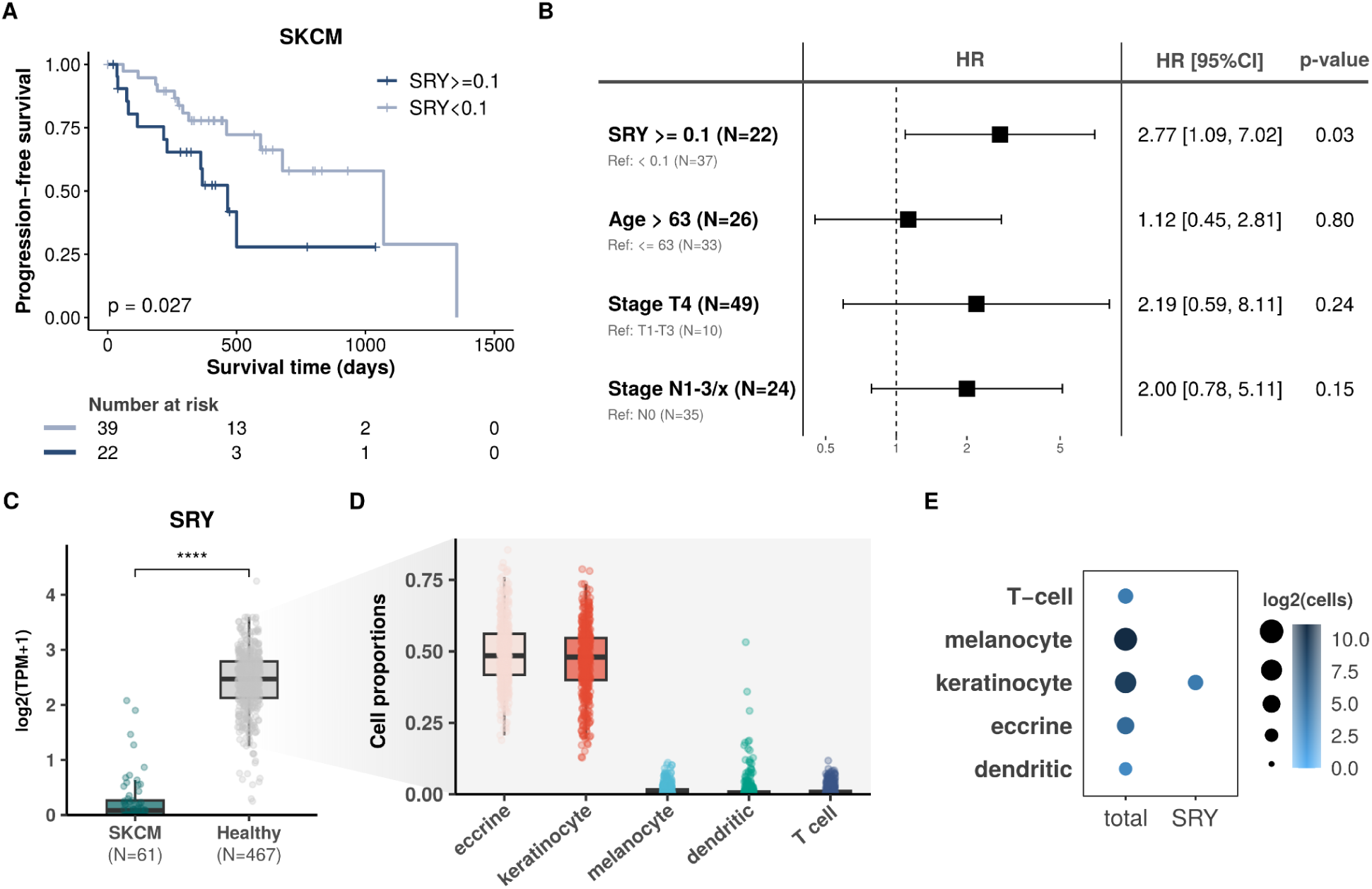
SRY is associated with worse prognosis in men with melanoma and over-expressed in melanocytes only in tumors, but not in healthy skin. **A)** Progression-free survival of men with melanoma expressing SRY at high (TPM ≥ 0.1) or low (TPM < 0.1) levels in the TCGA cohort. **B)** Multivariate Cox regression of melanoma patients considering SRY expression together with clinical information about age (median: 63 years), and AJCC T (T1 - T4) and N (N0 - N3) stages. **C)** Expression levels of SRY in melanoma tumors from men in the TCGA cohort are underexpressed versus healthy sun-exposed skin samples (GTEx cohort) (Wilcoxon test; **** p-value ≤ 0.0001). **D)** Proportions of distinct cell types estimated from bulk RNA-seq data of 464 healthy skin samples (p-value < 0.05). **E)** In healthy skin samples from men in a scRNA-seq cohort (accession: GSE151091), SRY is expressed only in keratinocytes.

We then profiled SRY expression in tumors from men with melanoma versus healthy skin samples. SRY was underexpressed in these tumors (p-value < 0.0001; Wilcoxon rank test) compared to healthy sun-exposed skin samples from the Genotype-Tissue Expression Project (GTEx) cohort (Figure 4C). Since human skin is composed of three layers made up of different cells (e.g., keratinocytes, melanocytes, Langerhans cells, and eccrine cells), we hypothesized that SRY may have distinct expression in these cell types. To test this hypothesis, we performed deconvolution of bulk RNA-seq expression data (healthy sun-exposed skin samples from GTEx) using CIBERSORTx (Newman et al., 2019), a tool that combines support vector regression with prior knowledge on scRNA-Seq expression profiles to estimate cell fractions. In agreement with our hypothesis, healthy skin keratinocytes and eccrine cells have robust SRY expression, while melanocytes have limited SRY expression (Figure 4D). Next, we measured SRY expression in an original single cell dataset (Belote et al., 2021). Figure 4E shows that only keratinocytes express SRY in healthy skin. Altogether, these results indicate that SRY overexpression was associated with a poor prognosis for men with melanoma, similar to its role in HGG. Furthermore, the finding that SRY is also expressed in healthy skin, but not in melanocytes, suggests that its dysregulation in melanocytes may contribute to the development of melanomas.

### High expression of SRY activates genes associated with melanoma progression and aggressiveness

Similar to the analyses of HGG tumors (Figure 3), we explored the differential expression of SRY target genes (Li et al., 2014) (Supplementary Table S8). Four proto-oncogenes (epidermal growth factor receptor [EGFR], FLT, BCL11B and PAX5) were identified among overexpressed genes (Figure 5A). These genes have been implicated in various cellular processes related to cancer, including cell proliferation, differentiation, and apoptosis. For example, EGFR is often upregulated in tumors, including melanoma, and its high expression contributes to tumor development and progression by promoting cell growth and survival, inhibiting apoptosis, and enhancing tumor invasion and metastasis (Pastwińska et al., 2022; Sigismund et al., 2018); FLT3 has been explored as a potential therapeutic target and showed promising results in melanoma (Cueto and Sancho, 2021; Esche et al., 1998).

**Figure 5.**
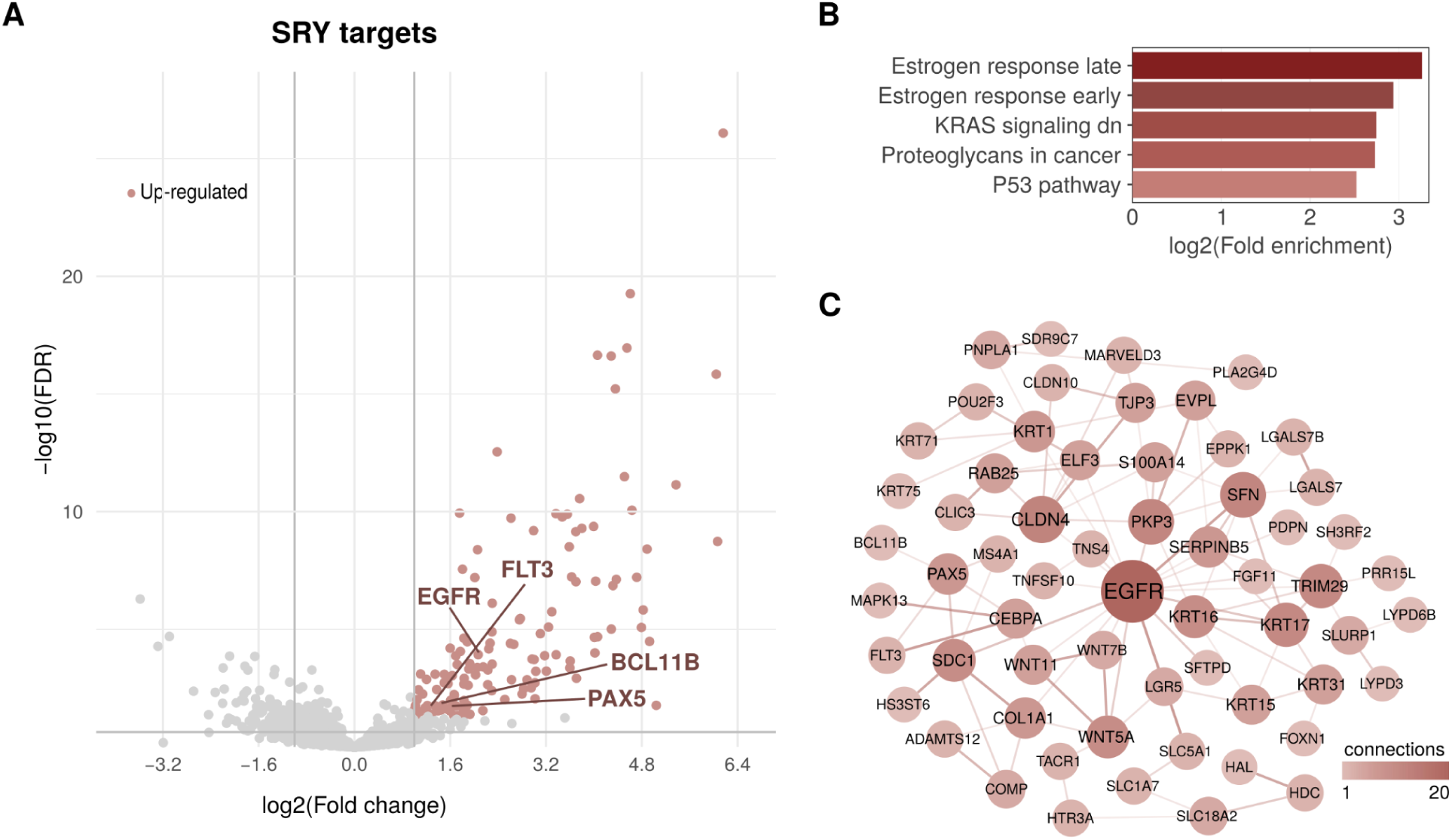
Several SRY targets are overexpressed in men with melanoma who had a worse prognosis. **A)** Overexpressed SRY target genes in SKCM samples with high expression of SRY (≥ 0.1 TPM). **B)** Functional enrichment analysis (KEGG pathways and MSigDB Hallmarks) of SRY target genes overexpressed in SKCM samples with high expression of SRY. **C)** Protein-protein interaction network of overexpressed SRY targets in SKCM patients with high expression of SRY.

Pathway enrichment analysis of target genes overexpressed in SKCM samples with higher SRY expression identified associations with key cancer pathways (Figure 4B). These included the KRAS signaling pathway, which is associated with cell proliferation, differentiation, and survival (Kim et al., 2021); proteoglycans in cancer, which is associated with changes in the composition and organization of the extracellular matrix, thereby affecting the tumor microenvironment and tumor progression (Barkovskaya et al., 2020); and, similar to HGG, the estrogen response (early and late) pathways. Moreover, two cellular pathways were shared between HGG and SKCM overexpressed SRY target genes: the estrogen response (late) and proteoglycans in cancer, suggesting their importance in tumors with high SRY expression.

In PPI analysis of target genes overexpressed in SKCM tumors with high SRY expression, EGFR emerged as the most highly connected protein (Figure 5C). Activation of the EGFR pathway leads to increased cell growth and tumor formation in many cancers, including melanoma. Other proteins related to melanoma progression and aggressiveness, such as WNT7B (Chen et al., 2023) , WNT5A (Bueno et al., 2022), WNT11 (Menck et al., 2021) (Wnt pathway) and other individual proteins (e.g., SFN (Shiba-Ishii, 2021), COL1A1 (Verginadis et al., 2022), CLDN4 (Hao et al., 2022), PKP3 (Ruan et al., 2021), and SDC1 (Maeda et al., 2006)), were highly connected in the PPI analysis. Overall, these results indicate that SRY target genes in samples with high SRY expression are implicated in cellular pathways associated with melanoma progression and aggressive tumors, supporting high SRY expression as a contributor to poor prognosis among men with melanoma (Figure 5).

## DISCUSSION

Although sexual dimorphism has been observed in many cancer types (Colopi et al., 2023; Costa et al., 2020; Suteau et al., 2021), our understanding of these sex disparities at the molecular level is still very limited. Here, we integrated and analyzed a large amount of gene expression data (including single-cell expression) along with clinical and functional datasets to perform a comprehensive analysis of SRY gene expression in men with high-grade glioma or melanoma. Overall, high SRY expression was associated with a worse prognosis and linked to oncogenic pathways in men with these two cancers.

High-grade glioma is more prevalent in men than women (Siegel et al., 2023), but the reasons for this difference are not fully understood and likely involve hormonal, genetic, and environmental factors (Clocchiatti et al., 2016). In our investigation, we discovered that high (TPM ≥ 0.1) expression of SRY, a male-specific gene, was a poor prognostic factor among men with high-grade glioma. Moreover, SRY’s target genes which were overexpressed in high-grade glioma patients with a worse prognosis, are involved in cellular pathways associated with tumorigenic processes such as extracellular matrix remodeling, cell adhesion, migration, angiogenesis, tumor invasion, and metastasis. Conversely, those with higher SRY expression also had underexpression of genes that, in normal conditions, facilitate infiltration and activity of immune cells within the tumor microenvironment. Thus, altogether these results indicate that SRY overexpression may activate key genes related to tumor aggressiveness and immune system modulation in high-grade glioma patients, and may be driving the worse prognosis in these patients.

Skin cutaneous melanoma (SKCM) is an increasingly common cancer and has a higher incidence rate in men than in women, with a lifetime risk of 1 in 26 for men and 1 in 41 for women (Hartman and Lin, 2019). Again, it is reasonable to suppose that this biased incidence is due to behavioral, genetic and hormonal factors. Our results show that men with melanoma and high SRY expression have a worse prognosis. As in high-grade glioma, tumors with a higher SRY expression showed overexpression of SRY target genes associated with cell proliferation, cell differentiation, and changes in the composition and organization of the extracellular matrix – all well-described cellular functions associated with melanomas aggressiveness. We found higher SRY expression in healthy skin than in melanoma, but this expression is limited to non-melanocyte cells, mostly keratinocytes. These findings suggest that SRY overexpression in melanoma (mainly melanocytes) may be a dysregulation that contributes to activation of genes involved in tumorigenesis, tumor progression and aggressiveness.

Among all cellular pathways enriched in men with high-grade glioma or melanoma and overexpression of SRY, only two were shared, the estrogen response (late) and the proteoglycans in cancer pathways. The estrogen response cellular signaling pathway is involved in the regulation of various physiologic processes, including cell proliferation and differentiation. While estrogens regulate many aspects of reproductive development and function in women, they also play a role in men. In men, estrogen 17β-estradiol (E2) is a relevant hormone for hypothalamic-pituitary-testicular axis regulation, reproductive function, growth hormone-insulin-like growth factor-1 axis regulation, bone growth, skeletal health, glucose metabolism, brain development, and behavior (Mahboobifard et al., 2022; Russell and Grossmann, 2019). Moreover, in men, estrogens are synthesized by aromatase (an enzyme that converts androgens to estrogens), which is expressed in several somatic tissues, especially in the brain (Cisternas et al., 2018). In the male brain, aromatase is involved in several brain functions, including synaptic plasticity, neurogenesis, neuroprotection, and regulation of sexual behavior (Azcoitia et al., 2021; Cisternas et al., 2018). On the other hand, high levels of estrogen in men are linked to an increased risk of some cancers, including prostate cancer (Nelles et al., 2011). The fact that the SRY DNA-binding domain specifically recognizes proximal upstream elements in promoters of sex-specific genes encoding P450 aromatase suggests a connection between SRY and aromatase expression (and thus estrogen production) (Haqq et al., 1993; Larney et al., 2014).

Additionally, the effects of SRY expression on the increased expression of genes related to epithelial-mesenchymal transition (EMT) may also be associated with estrogens (Hernández-Vega et al., 2020). Estrogen-promoted signaling is related to induction of EMT in estrogen-responsive tissues (Diaz-Ruano et al., 2023; Hernández-Vega et al., 2020). In ovarian and prostate cancer, estradiol treatment induces estrogen receptor (ER)-α-dependent EMT, which is inhibited by receptor silencing (Di Zazzo et al., 2019). Nevertheless, loss of ER-α expression in breast and endometrial cancer promotes morphologic changes, motility, improved invasion, and increased expression of EMT markers (Bouris et al., 2015; Liu et al., 2020). In human glioblastoma-derived cell lines, 17β-estradiol induced EMT via changes in cell morphology, expression of EMT markers, and cell migration and invasion assays (Hernández-Vega et al., 2020).

While no relationships between SRY and proteoglycans in tumors or tumorigenesis have been reported, evidence from developmental diseases may suggest a potential connection. One such disease, campomelic dysplasia, is a congenital skeletal abnormality in humans which manifests with notable skeletal deformities along with distinct facial features, including macrocephaly, micrognathia, cleft palate, flat nasal bridge, and sex reversal (Kayhan et al., 2019). The genetic basis of this condition involves mutations in the SRY-related gene SOX9 (Csukasi et al., 2019). Specifically, SOX9 expression initiates during mesenchymal condensation preceding cartilage formation and remains within the perichondrium and chondrocytes. Thus, this transcriptional activator likely plays a significant role in the establishment and maintenance of the chondrocytic phenotype by regulating cartilage-specific genes, including collagen types II and XI, aggrecan, and proteoglycans (Haseeb et al., 2021). Consequently, SRY indirectly influences the expression of proteoglycans through its interaction with SOX9. Therefore, in conditions with high SRY expression, an upregulation of proteoglycans would be expected, reinforcing our findings in both glioblastoma and melanoma.

In light of our findings linking high SRY expression and estrogen-response genes in high-grade glioma and melanoma, we raise the question whether endocrine therapy, which involves drugs that block the estrogen receptor or reduce estrogen levels, could benefit these patients. While we have found limited evidence to support the use of endocrine therapy to treat high-grade glioma or melanoma, based on our data, clinical trials may be warranted, particularly in cases where other standard therapies have proven ineffective.

## CONCLUSION

Taken together, our results provide insight into the biological mechanisms driving two cancers that are more common in men – high-grade glioma and melanoma – and suggest novel targets for putative therapeutic intervention. Specifically, we demonstrate that SRY overexpression predicts poor prognosis in high-grade glioma and melanoma, and we showed that this high SRY expression may activate target genes and cellular pathways involved in tumorigenesis, tumor progression, and immune system repression. Further studies are needed to explore the potential of these findings in developing new treatments for men with high-grade glioma or melanoma.

## MATERIALS AND METHODS

### RNA sequencing data

To determine cancers in The Cancer Genome Atlas - TCGA cohort that are more common in men (https://portal.gdc.cancer.gov), we obtained clinical data with gender information for 8,065 patients with 19 distinct types of solid primary tumors. Sex-specific tumor types and tumor types represented by fewer than 150 patients were not considered for analysis. We defined cancers more common in men as those with male to female ratios ≥ 1.2.

For the 11 tumor types identified as more common in men, we downloaded all available processed RNA-Seq data (raw gene counts and transcript per million [TPM] expression levels) from men: bladder urothelial carcinoma (BLCA, N=300), esophageal carcinoma (ESCA, N=158), high-grade glioma (HGG, N=248), kidney renal clear cell carcinoma (KIRC, N=345), kidney renal papillary cell carcinoma (KIRP, N=214), low-grade glioma (LGG, N=136), liver hepatocellular carcinoma (LIHC, N=250), lung squamous cell carcinoma (LUSC, N=371), pancreatic adenocarcinoma (PAAD, N=98), skin cutaneous melanoma (SKCM, N=61), and stomach adenocarcinoma (STAD, N=267). We evaluated the median TPM-normalized expression levels of SRY and considered for further analyses only tumors with a median TPM expression ≥ 0. These were: BLCA, ESCA, LGG, LUSC, HGG, PAAD and SKCM.

To compare SRY expression levels between these selected tumor samples and corresponding healthy tissue samples from men, we also obtained processed RNA-Seq data of 1,780 samples from The Genotype-Tissue Expression (GTEx Release V8) data portal (https://gtexportal.org): bladder (N=14), esophagus (N=363), brain (frontal) cortex (N=334), lung (N=395), pancreas (N=207), and skin (N=467).

To evaluate our results in gliomas in an independent cohort, we also obtained processed RNA-Seq data from men in the Chinese Glioma Genome Atlas - CGGA (http://www.cgga.org.cn), comprising 238 HGG and 131 LGG samples of primary tumors.

### Survival analyses

To evaluate whether SRY expression impacts the prognosis of cancer patients, we obtained progression-free survival data from TCGA and curated by Liu *et al*. (Liu et al., 2018), and retrieved additional clinical information using the R package *TCGAbiolinks* (version 2.24.3). Overall survival and clinical data for the CGGA cohort were obtained directly from the CGGA data portal.

For each tumor type, we stratified patients into two groups based on SRY expression: patients with high (TPM ≥ 0.1) or low (TPM < 0.1) SRY expression. Survival analyses between these groups were then performed using log-rank tests; results were considered significant at a p-value < 0.05. Kaplan-Meier survival curves were created using the R packages *survival* (version 3.5.3) and *survminer* (version 0.4.9). For tumors with significant survival differences (p-value < 0.05), we next performed multivariate Cox regression analyses to evaluate the prognostic value of SRY in context with other relevant clinical variables. Only variables with < 5% of missing values were included. Forest plots were built using the R package *forestmodel* (version 0.6.2).

### Single-cell RNA sequencing data

To check SRY expression in single-cell RNA sequencing (scRNA-Seq) data obtained from patients with primary HGG, we obtained data from two previously published studies: (Neftel et al., 2019) (GEO accession: GSE131928) and (Jacob et al., 2020) (GEO accession: GSE141946). For the first dataset, SRY expression was directly assessed in 3,825 cells from 14 men. For the second dataset, scRNA-Seq data from 3 male individuals was reprocessed using the R package *Seurat* (version 4.3.0) (Hao et al., 2021). Briefly, only cells with (a) at least 400 and at most 5,000 genes detected and (b) less than 25% mitochondrial reads were retained for further analysis (8,892 cells). Standard preprocessing steps, including normalization, scaling, and cell cycle phase scoring, were run without modifications. UMAP dimensionality reduction was performed with default parameters, and the Seurat function *FindAllMarkers* was used to perform differential gene expression with thresholds set to log2(fold-change) > 0.1 and minimum percent expressed > 0.1. The top 10 marker genes based on highest log2(fold-changes) were selected for each cell type.

Moreover, we evaluated SRY expression in 2,981 skin cells of 8 healthy men using processed scRNA-Seq data obtained directly from a previous study (Belote et al., 2021) (GEO accession: GSE151091).

### Gene expression analyses

Differences of SRY expression levels (TPM-normalized) between healthy and tumor samples, and between distinct glioma grades, were assessed using Wilcoxon rank tests (R package *ggpubr* version 0.5.0). Boxplots were generated using the R package *ggplot2* (version 3.4.0).

For men with HGG or SKCM, we performed differential gene expression analyses between individuals with high (TPM ≥ 0.1) or low (TPM < 0.1) SRY expression using DESeq2 (Love et al., 2014). Only genes with absolute log2(fold change) > 1 and a false discovery rate (FDR) adjusted p-value < 0.05 were considered as differentially expressed. Differentially expressed SRY target genes were identified based on previous ChIP-Chip experiments performed by Li and collaborators (Li et al., 2014).

### Functional and protein-protein network analyses

To elucidate the potential biological impact of SRY expression in men with HGG or SKCM, we performed functional enrichment analyses of differentially expressed SRY targets using the ShinyGO tool (v 0.77) (Ge et al., 2019). As input reference datasets, we used KEGG pathways (Release 86.1, https://www.genome.jp/kegg) and cancer hallmarks from the Molecular Signatures Database - MSigDB (Release 6.1, http://www.gsea-msigdb.org/gsea/msigdb/).

Protein-protein networks of differentially expressed SRY target proteins were constructed using the STRING database (v 2023) (Szklarczyk et al., 2023) and Cytoscape (v 3.8.0, https://cytoscape.org). Only experimental and text-mining protein-protein interactions with scores > 0.4 were considered, and only proteins with at least one interaction were maintained.

### Cell fraction imputation of healthy skin samples

To estimate the proportions of melanocytes, keratinocytes, eccrine, dendritic cells, and T-cells in RNA-Seq samples obtained from GTEx (467 healthy skin samples), we used the CIBERSORTx tool (Newman et al., 2019). This tool uses support vector regression combined with prior knowledge on scRNA-Seq expression profiles (reference datasets) to estimate the abundance of distinct cell subpopulations in bulk RNA-Seq samples. As input for the reference dataset, we provided the scRNA-Seq expression profile of marker genes used to cluster the five cell types in healthy skin samples from a previous study (Belote et al., 2021). Imputation of cell fractions was run in CIBERSORTx with batch correction enabled (B-mode), disabled quantile normalization, relative run mode, and 1,000 permutations. Only samples with a p-value < 0.05 were considered for further analyses.

## Data Availability

All used data are publicly available through the GEO (GSE131928, GSE141946), GDC portal (https://portal.gdc.cancer.gov), the Genotype-Tissue Expression (https://gtexportal.org) and the Chinese Glioma Genome Atlas - CGGA (http://www.cgga.org.cn).

## Authors’ Contributions

**G. D. A. Guardia:** conceptualization, investigation, formal analysis, methodology, writing-original draft. **R. L. Batista:** conceptualization, investigation, methodology, writing-original draft. **L. O Penalva:** conceptualization, investigation, methodology, writing-original draft. **P. A. F. Galante:** conceptualization, investigation, methodology, writing-original draft, project administration, funding acquisition.

## Acknowledgments

Financial support: This work was supported by grant #2018/15579-8, São Paulo Research Foundation (FAPESP) to PAFG. Partially supported by funds from Serrapilheira Foundation, CNPq and Hospital Sírio-Libanês to PAFG.

