## Supplemental Figures and Tables for "Unveiling the role of SRY in male-biased cancers: Insights into the molecular basis of sex disparities in high-grade glioma and melanoma"

### Supplementary Figures

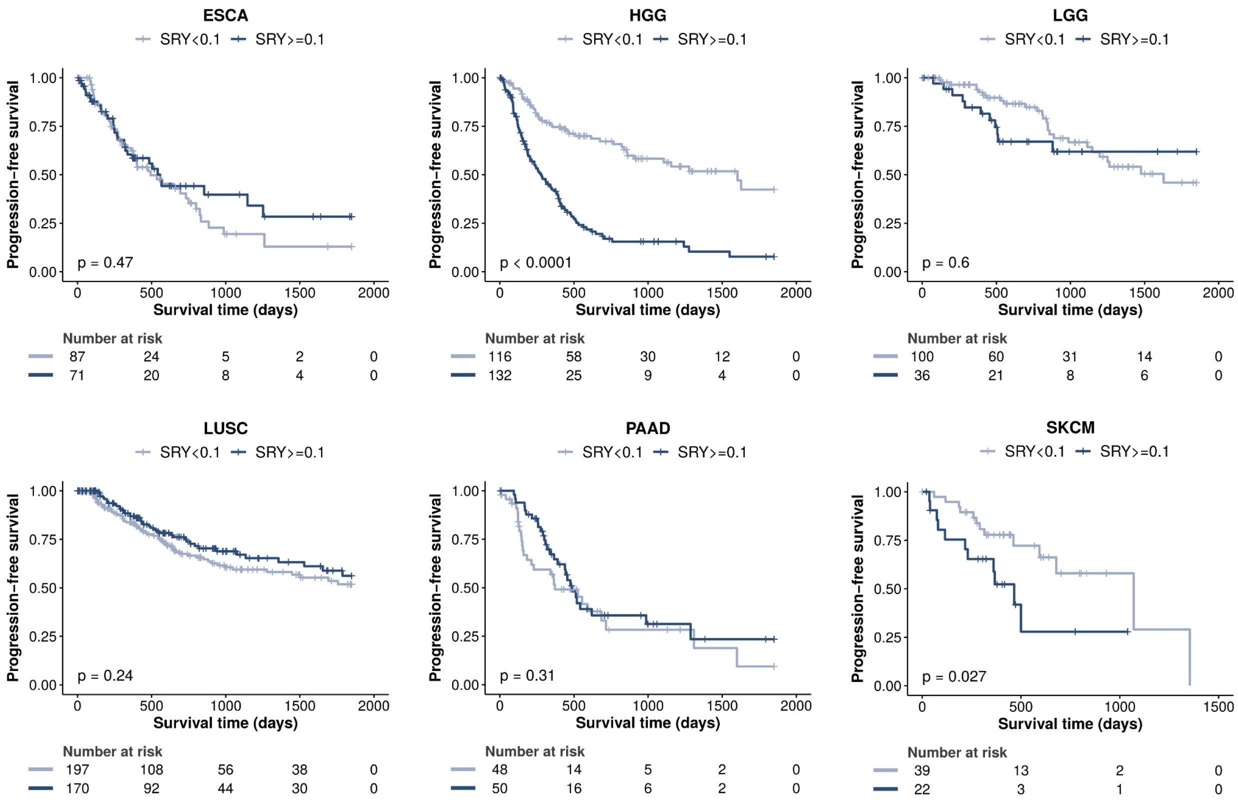

**Figure S1.** Progression-free survival curves of men with ESCA, HGG, LGG, LUSC, PAAD and SKCM tumors stratified based on high/low expression of SRY (TPM >= 0.1 / TPM < 0.1).

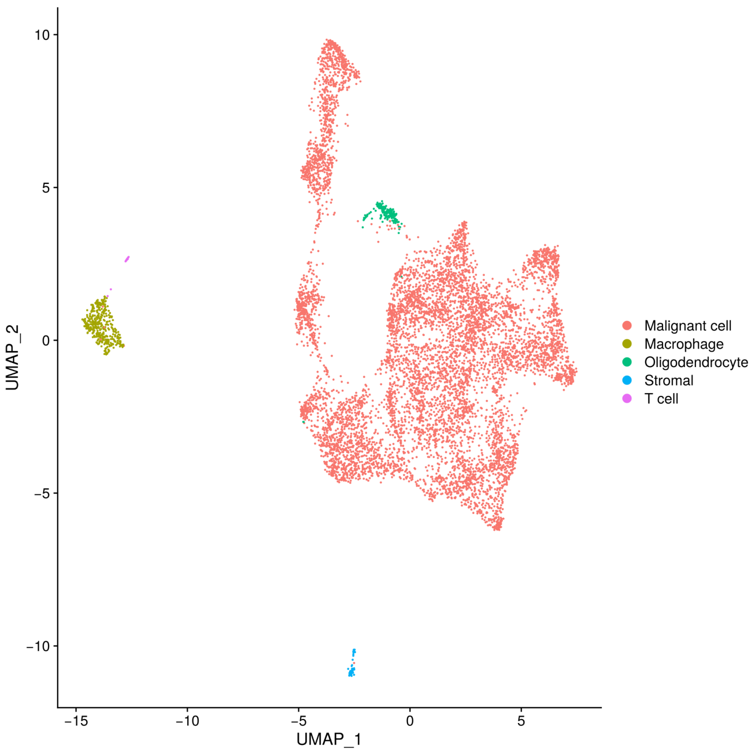

**Figure S2.** UMAP plot of single-cell RNA expression of two men with HGG (UP-8165-C and UP-8165-PV) from GEO study GSE141946. Cell clusters identified include malignant cells, macrophages, oligodendrocytes, and stromal and T cells.

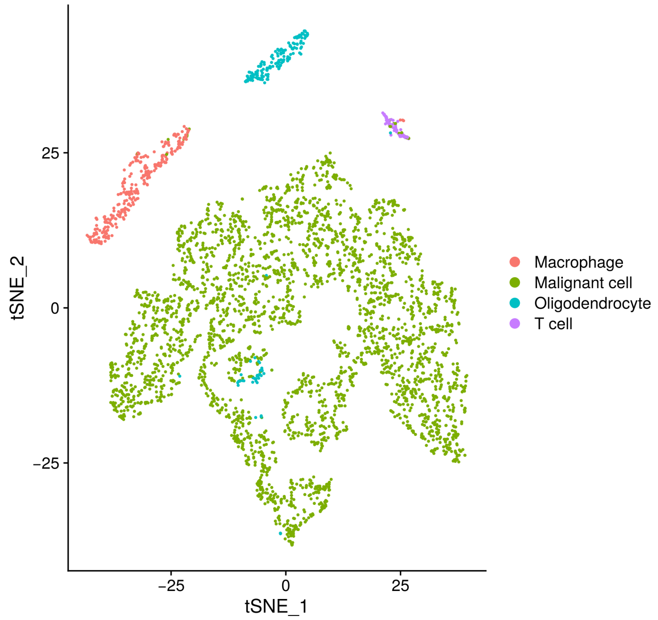

**Figure S3.** tSNE plot of single-cell RNA expression of five men with HGG (MGH102, MGH122, MGH125, MGH151, MGH152) from GEO study GSE131928. Cell clusters identified include malignant cells, macrophages, oligodendrocytes, and T cells.

**Table S1. Male-prevalent TCGA tumor types.**

| project | number of male samples | number of female samples | total samples | male:female ratio |
|---|---|---|---|---|
| TCGA-ESCA | 158 | 27 | 185 | 5.85 |
| TCGA-LUSC | 373 | 131 | 504 | 2.85 |
| TCGA-BLCA | 304 | 108 | 412 | 2.81 |
| TCGA-KIRP | 214 | 77 | 291 | 2.78 |
| TCGA-LIHC | 255 | 122 | 377 | 2.09 |
| TCGA-KIRC | 346 | 191 | 537 | 1.81 |
| TCGA-STAD | 285 | 158 | 443 | 1.8 |
| TCGA-SKCM | 186 | 112 | 298 | 1.66 |
| TCGA-HGG | 514 | 347 | 861 | 1.48 |
| TCGA-PAAD | 102 | 83 | 185 | 1.23 |
| TCGA-LGG | 136 | 113 | 249 | 1.2 |

**Table S2. HGG patients from the TCGA cohort.**

| Patient id | Gender | Age | Tumor type | Histology | Grade | IDH status | Codeletion 1p 19q | MGMT status | Gain chr7 & Loss chr10 | Cogain chr19/20 | TERT status | TERT expressed | PFI status | PFI time | SRY expression |
|---|---|---|---|---|---|---|---|---|---|---|---|---|---|---|---|
| TCGA-02-0047 | Male | 78 | Primary | Glioblastoma | IV | Wildtype | Non-codel | Unmethylated | No | No | [Not available] | Yes | 1 | 57 | 0.11610 |
| TCGA-02-2483 | Male | 43 | Primary | Astrocytoma | III | Mutant | Non-codel | Methylated | No | No | Wildtype | Yes | 0 | 466 | 0.00000 |
| TCGA-02-2485 | Male | 53 | Primary | Glioblastoma | IV | Wildtype | Non-codel | Unmethylated | Yes | No | Mutant | Yes | 1 | 186 | 2.10850 |
| TCGA-02-2486 | Male | 64 | Primary | Glioblastoma | IV | Wildtype | Non-codel | Unmethylated | Yes | No | [Not available] | Yes | 1 | 618 | 5.16450 |
| TCGA-06-0129 | Male | 30 | Primary | Astrocytoma | III | Mutant | Non-codel | Methylated | No | No | [Not available] | No | 1 | 148 | 0.00000 |
| TCGA-06-0130 | Male | 54 | Primary | Glioblastoma | IV | Wildtype | Non-codel | Unmethylated | No | No | [Not available] | Yes | 1 | 244 | 0.00000 |
| TCGA-06-0132 | Male | 49 | Primary | Glioblastoma | IV | Wildtype | Non-codel | [Not available] | No | No | [Not available] | No | 1 | 482 | 0.16350 |
| TCGA-06-0138 | Male | 43 | Primary | Glioblastoma | IV | Wildtype | Non-codel | [Not available] | Yes | No | [Not available] | Yes | 1 | 394 | 4.01550 |
| TCGA-06-0139 | Male | 40 | Primary | Glioblastoma | IV | Wildtype | Non-codel | Unmethylated | No | No | [Not available] | [Not available] | 1 | 152 | 0.80970 |
| TCGA-06-0141 | Male | 62 | Primary | Glioblastoma | IV | Wildtype | Non-codel | Unmethylated | No | No | [Not available] | Yes | 1 | 145 | 0.11590 |
| TCGA-06-0156 | Male | 57 | Primary | Astrocytoma | III | Mutant | [Not available] | [Not available] | [Not available] | [Not available] | [Not available] | Yes | 1 | 178 | 4.91130 |
| TCGA-06-0158 | Male | 73 | Primary | Glioblastoma | IV | Wildtype | Non-codel | [Not available] | No | Yes | [Not available] | Yes | 1 | 90 | 3.39460 |
| TCGA-06-0174 | Male | 54 | Primary | Glioblastoma | IV | Wildtype | Non-codel | [Not available] | Yes | No | [Not available] | Yes | 1 | 47 | 0.10580 |
| TCGA-06-0178 | Male | 38 | Primary | Astrocytoma | III | Mutant | Non-codel | [Not available] | No | No | [Not available] | No | 1 | 192 | 0.27200 |
| TCGA-06-0184 | Male | 63 | Primary | Glioblastoma | IV | Wildtype | Non-codel | [Not available] | Yes | No | [Not available] | Yes | 1 | 1276 | 0.29890 |
| TCGA-06-0187 | Male | 69 | Primary | Glioblastoma | IV | Wildtype | Non-codel | [Not available] | No | No | [Not available] | Yes | 1 | 531 | 0.57280 |
| TCGA-06-0190 | Male | 62 | Primary | Glioblastoma | IV | Wildtype | Non-codel | [Not available] | Yes | No | Mutant | Yes | 1 | 88 | 2.16620 |
| TCGA-06-0211 | Male | 47 | Primary | Glioblastoma | IV | Wildtype | Non-codel | [Not available] | Yes | Yes | Mutant | Yes | 1 | 53 | 0.00000 |
| TCGA-06-0219 | Male | 67 | Primary | Glioblastoma | IV | Wildtype | Non-codel | [Not available] | No | No | [Not available] | Yes | 1 | 22 | 9.94540 |
| TCGA-06-0238 | Male | 46 | Primary | Glioblastoma | IV | Wildtype | Non-codel | [Not available] | Yes | No | [Not available] | Yes | 1 | 311 | 1.98490 |
| TCGA-06-0644 | Male | 71 | Primary | Glioblastoma | IV | Wildtype | Non-codel | [Not available] | Yes | No | [Not available] | Yes | 1 | 85 | 0.35260 |
| TCGA-06-0646 | Male | 60 | Primary | Glioblastoma | IV | Wildtype | Non-codel | [Not available] | Yes | No | [Not available] | Yes | 1 | 90 | 2.84710 |
| TCGA-06-0686 | Male | 53 | Primary | Glioblastoma | IV | Wildtype | Non-codel | [Not available] | Yes | No | Mutant | Yes | 1 | 160 | 0.00000 |
| TCGA-06-0743 | Male | 69 | Primary | Glioblastoma | IV | Wildtype | Non-codel | [Not available] | Yes | Yes | [Not available] | Yes | 1 | 176 | 2.51430 |
| TCGA-06-0744 | Male | 66 | Primary | Glioblastoma | IV | Wildtype | Non-codel | [Not available] | Yes | No | Mutant | Yes | 1 | 1277 | 0.00000 |
| TCGA-06-0745 | Male | 59 | Primary | Glioblastoma | IV | Wildtype | Non-codel | [Not available] | Yes | No | Mutant | Yes | 1 | 92 | 3.85930 |
| TCGA-06-0747 | Male | 53 | Primary | Glioblastoma | IV | Wildtype | Non-codel | [Not available] | No | No | [Not available] | Yes | 1 | 82 | 2.73630 |
| TCGA-06-0749 | Male | 50 | Primary | Glioblastoma | IV | Wildtype | Non-codel | [Not available] | Yes | No | [Not available] | Yes | 1 | 82 | 0.00000 |
| TCGA-06-0750 | Male | 43 | Primary | Glioblastoma | IV | Wildtype | Non-codel | [Not available] | Yes | No | [Not available] | Yes | 1 | 28 | 0.15900 |
| TCGA-06-0878 | Male | 74 | Primary | Glioblastoma | IV | Wildtype | Non-codel | Unmethylated | Yes | No | [Not available] | Yes | 1 | 66 | 6.13500 |
| TCGA-06-0882 | Male | 30 | Primary | Glioblastoma | IV | Wildtype | Non-codel | Unmethylated | Yes | No | [Not available] | No | 1 | 213 | 0.98820 |
| TCGA-06-2557 | Male | 76 | Primary | Glioblastoma | IV | Wildtype | Non-codel | Unmethylated | No | No | Mutant | Yes | 1 | 33 | 0.16370 |
| TCGA-06-2559 | Male | 83 | Primary | Glioblastoma | IV | Wildtype | Non-codel | Methylated | No | No | [Not available] | Yes | 1 | 150 | 0.00000 |
| TCGA-06-2562 | Male | 81 | Primary | Glioblastoma | IV | Wildtype | Non-codel | Unmethylated | Yes | No | [Not available] | Yes | 1 | 151 | 0.00000 |
| TCGA-06-2564 | Male | 50 | Primary | Glioblastoma | IV | Wildtype | Non-codel | Unmethylated | Yes | No | [Not available] | Yes | 0 | 181 | 2.19410 |
| TCGA-06-2565 | Male | 59 | Primary | Glioblastoma | IV | Wildtype | Non-codel | Methylated | No | Yes | [Not available] | Yes | 1 | 178 | 1.20240 |
| TCGA-06-2567 | Male | 65 | Primary | Glioblastoma | IV | Wildtype | Non-codel | Methylated | Yes | No | [Not available] | Yes | 1 | 133 | 0.00000 |
| TCGA-06-5411 | Male | 51 | Primary | Glioblastoma | IV | Wildtype | Non-codel | Unmethylated | Yes | No | Mutant | Yes | 1 | 214 | 0.07460 |
| TCGA-06-5413 | Male | 67 | Primary | Glioblastoma | IV | Wildtype | Non-codel | Unmethylated | Yes | No | [Not available] | Yes | 1 | 195 | 0.00000 |
| TCGA-06-5414 | Male | 61 | Primary | Glioblastoma | IV | Wildtype | Non-codel | Unmethylated | Yes | Yes | [Not available] | Yes | 1 | 167 | 1.09790 |
| TCGA-06-5415 | Male | 60 | Primary | Glioblastoma | IV | Wildtype | Non-codel | Unmethylated | Yes | Yes | Mutant | Yes | 0 | 260 | 0.00000 |
| TCGA-06-5856 | Male | 58 | Primary | Glioblastoma | IV | Wildtype | Non-codel | Unmethylated | Yes | No | [Not available] | Yes | 1 | 114 | 0.19220 |
| TCGA-06-5859 | Male | 63 | Primary | Glioblastoma | IV | Wildtype | Non-codel | Unmethylated | Yes | No | [Not available] | Yes | 0 | 139 | 13.04940 |
| TCGA-08-0386 | Male | 74 | Primary | Glioblastoma | IV | Wildtype | Non-codel | [Not available] | Yes | No | [Not available] | Yes | 1 | 427 | 5.08580 |
| TCGA-12-0618 | Male | 49 | Primary | Glioblastoma | IV | Wildtype | Non-codel | [Not available] | Yes | No | [Not available] | Yes | 0 | 395 | 1.85270 |
| TCGA-12-0619 | Male | 60 | Primary | Glioblastoma | IV | Wildtype | Non-codel | [Not available] | Yes | No | [Not available] | Yes | 1 | 203 | 0.40910 |
| TCGA-12-0821 | Male | 62 | Primary | Glioblastoma | IV | Wildtype | Non-codel | Unmethylated | Yes | No | [Not available] | Yes | 1 | 259 | 3.48080 |

| | | | | | | | | | | | | | | | |
|---|---|---|---|---|---|---|---|---|---|---|---|---|---|---|---|
| TCGA-12-3650 | Male | 46 | Primary | Glioblastoma | IV | Wildtype | Non-codel | Unmethylated | Yes | No | [Not available] | Yes | 1 | 239 | 0.00000 |
| TCGA-12-3652 | Male | 60 | Primary | Glioblastoma | IV | Wildtype | Non-codel | Unmethylated | No | Yes | [Not available] | Yes | 1 | 203 | 6.62780 |
| TCGA-14-0781 | Male | 49 | Primary | Glioblastoma | IV | Wildtype | Non-codel | Unmethylated | No | No | [Not available] | Yes | 1 | 29 | 1.47310 |
| TCGA-14-0787 | Male | 69 | Primary | Glioblastoma | IV | Wildtype | Non-codel | Methylated | Yes | No | [Not available] | Yes | 0 | 68 | 3.75450 |
| TCGA-14-0789 | Male | 54 | Primary | Glioblastoma | IV | Wildtype | Non-codel | Methylated | Yes | No | [Not available] | Yes | 1 | 105 | 0.59240 |
| TCGA-14-1825 | Male | 70 | Primary | Glioblastoma | IV | Wildtype | Non-codel | Unmethylated | Yes | No | [Not available] | Yes | 1 | 82 | 1.66800 |
| TCGA-14-1829 | Male | 57 | Primary | Glioblastoma | IV | Wildtype | Non-codel | Unmethylated | Yes | No | [Not available] | Yes | 0 | 218 | 1.48720 |
| TCGA-15-0742 | Male | 65 | Primary | Glioblastoma | IV | Wildtype | Non-codel | [Not available] | Yes | No | [Not available] | Yes | 1 | 232 | 0.18430 |
| TCGA-15-1444 | Male | 21 | Primary | Astrocytoma | III | Mutant | Non-codel | Methylated | No | No | Wildtype | No | 1 | 1550 | 0.39810 |
| TCGA-16-0846 | Male | 85 | Primary | Glioblastoma | IV | Wildtype | Non-codel | Methylated | Yes | No | [Not available] | No | 1 | 119 | 2.82420 |
| TCGA-19-1787 | Male | 48 | Primary | [Not available] | III | [Not available] | Non-codel | Methylated | Yes | No | [Not available] | No | 1 | 308 | 4.41950 |
| TCGA-19-2620 | Male | 70 | Primary | Glioblastoma | IV | Wildtype | Non-codel | Methylated | No | No | Mutant | Yes | 0 | 148 | 0.00000 |
| TCGA-19-2624 | Male | 51 | Primary | Glioblastoma | IV | Wildtype | Non-codel | Unmethylated | Yes | No | Mutant | Yes | 0 | 5 | 0.67310 |
| TCGA-19-2629 | Male | 60 | Primary | Astrocytoma | III | Mutant | Non-codel | Unmethylated | No | No | Wildtype | No | 1 | 145 | 0.25450 |
| TCGA-19-4065 | Male | 36 | Primary | [Not available] | III | [Not available] | Non-codel | Unmethylated | Yes | No | [Not available] | Yes | 1 | 70 | 3.25240 |
| TCGA-19-5960 | Male | 56 | Primary | Glioblastoma | IV | Wildtype | Non-codel | Unmethylated | Yes | No | Mutant | Yes | 1 | 382 | 12.66690 |
| TCGA-26-1442 | Male | 43 | Primary | Astrocytoma | III | Mutant | Non-codel | Methylated | No | No | [Not available] | No | 0 | 953 | 0.00000 |
| TCGA-26-5132 | Male | 74 | Primary | Glioblastoma | IV | Wildtype | Non-codel | Methylated | Yes | No | Mutant | Yes | 0 | 286 | 0.09280 |
| TCGA-26-5133 | Male | 59 | Primary | Glioblastoma | IV | Wildtype | Non-codel | Unmethylated | No | No | [Not available] | No | 1 | 370 | 0.50110 |
| TCGA-26-5134 | Male | 74 | Primary | Glioblastoma | IV | Wildtype | Non-codel | Unmethylated | No | No | [Not available] | Yes | 0 | 167 | 0.59470 |
| TCGA-27-1830 | Male | 57 | Primary | Glioblastoma | IV | Wildtype | Non-codel | Unmethylated | Yes | No | [Not available] | Yes | 1 | 124 | 3.95570 |
| TCGA-27-1831 | Male | 66 | Primary | Glioblastoma | IV | Wildtype | Non-codel | Unmethylated | Yes | No | Mutant | Yes | 1 | 144 | 0.00000 |
| TCGA-27-1834 | Male | 56 | Primary | Glioblastoma | IV | Wildtype | Non-codel | Methylated | Yes | No | [Not available] | No | 1 | 335 | 3.21860 |
| TCGA-27-1837 | Male | 36 | Primary | Glioblastoma | IV | Wildtype | Non-codel | Methylated | No | No | [Not available] | Yes | 1 | 136 | 0.20200 |
| TCGA-27-2519 | Male | 48 | Primary | Glioblastoma | IV | Wildtype | Non-codel | Unmethylated | Yes | No | [Not available] | No | 1 | 256 | 6.09980 |
| TCGA-27-2521 | Male | 34 | Primary | Astrocytoma | III | Mutant | Non-codel | Methylated | No | No | [Not available] | No | 1 | 510 | 2.58930 |
| TCGA-27-2523 | Male | 63 | Primary | Glioblastoma | IV | Wildtype | Non-codel | Methylated | Yes | No | Mutant | Yes | 1 | 402 | 0.36840 |
| TCGA-27-2524 | Male | 56 | Primary | Glioblastoma | IV | Wildtype | Non-codel | Unmethylated | Yes | No | [Not available] | Yes | 1 | 231 | 2.00220 |
| TCGA-27-2528 | Male | 62 | Primary | Glioblastoma | IV | Wildtype | Non-codel | Methylated | No | Yes | Mutant | Yes | 1 | 72 | 2.47860 |
| TCGA-28-1747 | Male | 44 | Primary | Glioblastoma | IV | Wildtype | Non-codel | Methylated | No | No | [Not available] | Yes | 0 | 77 | 2.77730 |
| TCGA-28-1753 | Male | 53 | Primary | Glioblastoma | IV | Wildtype | Non-codel | Unmethylated | Yes | No | [Not available] | Yes | 0 | 37 | 0.19810 |
| TCGA-28-2499 | Male | 59 | Primary | Glioblastoma | IV | Wildtype | [Not available] | Unmethylated | [Not available] | [Not available] | [Not available] | No | 0 | 95 | 1.02890 |
| TCGA-28-2514 | Male | 45 | Primary | Glioblastoma | IV | Wildtype | Non-codel | Unmethylated | No | No | [Not available] | Yes | 0 | 160 | 4.04040 |
| TCGA-28-5204 | Male | 72 | Primary | Glioblastoma | IV | Wildtype | Non-codel | Unmethylated | Yes | No | [Not available] | Yes | 1 | 454 | 0.32900 |
| TCGA-28-5207 | Male | 71 | Primary | Glioblastoma | IV | Wildtype | Non-codel | Unmethylated | Yes | No | [Not available] | Yes | 1 | 343 | 3.81310 |
| TCGA-28-5208 | Male | 52 | Primary | Glioblastoma | IV | Wildtype | Non-codel | Methylated | No | No | [Not available] | Yes | 1 | 148 | 5.56200 |
| TCGA-28-5213 | Male | 72 | Primary | Glioblastoma | IV | Wildtype | Non-codel | Unmethylated | Yes | No | [Not available] | Yes | 0 | 951 | 0.19900 |
| TCGA-28-5216 | Male | 52 | Primary | Glioblastoma | IV | Wildtype | Non-codel | Unmethylated | No | No | [Not available] | Yes | 0 | 415 | 1.56920 |
| TCGA-28-5218 | Male | 63 | Primary | Glioblastoma | IV | Wildtype | Non-codel | Unmethylated | No | Yes | [Not available] | No | 0 | 157 | 0.31740 |
| TCGA-28-5220 | Male | 67 | Primary | Glioblastoma | IV | Wildtype | Non-codel | Unmethylated | Yes | No | [Not available] | Yes | 1 | 262 | 0.00000 |
| TCGA-32-1970 | Male | 59 | Primary | Glioblastoma | IV | Wildtype | Non-codel | Unmethylated | Yes | No | Mutant | Yes | 1 | 408 | 3.85840 |
| TCGA-32-1980 | Male | 72 | Primary | Glioblastoma | IV | Wildtype | Non-codel | Unmethylated | No | No | [Not available] | No | 1 | 36 | 0.16200 |
| TCGA-32-2615 | Male | 62 | Primary | Glioblastoma | IV | Wildtype | Non-codel | Unmethylated | No | No | [Not available] | Yes | 1 | 131 | 2.11130 |
| TCGA-32-2632 | Male | 80 | Primary | Glioblastoma | IV | Wildtype | Non-codel | Unmethylated | No | No | [Not available] | Yes | 1 | 269 | 0.96130 |
| TCGA-32-2634 | Male | 82 | Primary | Glioblastoma | IV | Wildtype | Non-codel | Methylated | No | No | [Not available] | Yes | 0 | 693 | 20.26840 |
| TCGA-32-2638 | Male | 67 | Primary | Glioblastoma | IV | Wildtype | Non-codel | Methylated | Yes | Yes | [Not available] | Yes | 1 | 766 | 0.00000 |
| TCGA-32-5222 | Male | 66 | Primary | Glioblastoma | IV | Wildtype | Non-codel | Methylated | Yes | Yes | [Not available] | Yes | 1 | 118 | 10.93390 |
| TCGA-41-2571 | Male | 89 | Primary | Glioblastoma | IV | Wildtype | Non-codel | Unmethylated | Yes | No | [Not available] | Yes | 1 | 26 | 0.00000 |
| TCGA-41-2572 | Male | 67 | Primary | Glioblastoma | IV | Wildtype | Non-codel | Unmethylated | Yes | Yes | [Not available] | Yes | 1 | 122 | 6.28730 |
| TCGA-41-3915 | Male | 48 | Primary | Glioblastoma | IV | Wildtype | Non-codel | Methylated | Yes | No | [Not available] | Yes | 1 | 288 | 3.58850 |

| | | | | | | | | | | | | | | | |
|---|---|---|---|---|---|---|---|---|---|---|---|---|---|---|---|
| TCGA-76-4925 | Male | 76 | Primary | Glioblastoma | IV | Wildtype | Non-codel | Methylated | Yes | No | [Not available] | Yes | 1 | 88 | 0.08140 |
| TCGA-76-4926 | Male | 68 | Primary | Glioblastoma | IV | Wildtype | Non-codel | Unmethylated | Yes | No | [Not available] | Yes | 1 | 34 | 2.49670 |
| TCGA-76-4927 | Male | 58 | Primary | Glioblastoma | IV | Wildtype | [Not available] | Unmethylated | [Not available] | [Not available] | [Not available] | Yes | 1 | 416 | 0.00000 |
| TCGA-CS-4941 | Male | 67 | Primary | Astrocytoma | III | Wildtype | Non-codel | Methylated | Yes | No | Mutant | Yes | 1 | 9 | 8.57350 |
| TCGA-CS-4943 | Male | 37 | Primary | Astrocytoma | III | Mutant | Non-codel | Methylated | No | No | Wildtype | No | 1 | 1106 | 0.07900 |
| TCGA-CS-5393 | Male | 39 | Primary | Astrocytoma | III | Mutant | Non-codel | Methylated | No | No | Wildtype | No | 0 | 1222 | 0.00000 |
| TCGA-CS-5394 | Male | 40 | Primary | Astrocytoma | III | Mutant | Non-codel | Methylated | No | No | Wildtype | No | 0 | 8 | 0.00000 |
| TCGA-CS-6186 | Male | 58 | Primary | Oligoastrocytoma | III | Wildtype | Non-codel | Unmethylated | Yes | No | Mutant | Yes | 1 | 188 | 1.89180 |
| TCGA-CS-6188 | Male | 48 | Primary | Astrocytoma | III | Wildtype | Non-codel | Unmethylated | Yes | No | Mutant | Yes | 1 | 647 | 0.73130 |
| TCGA-CS-6290 | Male | 31 | Primary | Astrocytoma | III | Mutant | Non-codel | Methylated | No | No | Wildtype | No | 0 | 1137 | 0.00000 |
| TCGA-CS-6666 | Male | 22 | Primary | Astrocytoma | III | Mutant | Non-codel | Methylated | No | No | Wildtype | No | 0 | 1428 | 0.00000 |
| TCGA-CS-6670 | Male | 43 | Primary | Oligodendroglioma | III | Mutant | Codel | Methylated | No | No | [Not available] | Yes | 0 | 1426 | 0.06240 |
| TCGA-DB-5273 | Male | 33 | Primary | Astrocytoma | III | Mutant | Non-codel | Unmethylated | No | No | Wildtype | No | 0 | 1850 | 0.08070 |
| TCGA-DB-5275 | Male | 36 | Primary | Oligoastrocytoma | III | Mutant | Non-codel | Methylated | No | No | Wildtype | No | 0 | 1458 | 0.00000 |
| TCGA-DB-5276 | Male | 32 | Primary | Oligoastrocytoma | III | Mutant | Non-codel | Methylated | No | No | Wildtype | No | 0 | 1850 | 0.25640 |
| TCGA-DB-5277 | Male | 34 | Primary | Astrocytoma | III | Mutant | Non-codel | Methylated | No | No | Wildtype | No | 1 | 1156 | 0.00000 |
| TCGA-DB-5281 | Male | 61 | Primary | Oligoastrocytoma | III | Mutant | Non-codel | Methylated | No | No | Wildtype | No | 0 | 1850 | 0.00000 |
| TCGA-DB-A4XB | Male | 38 | Primary | Astrocytoma | III | Mutant | Non-codel | Methylated | No | No | Wildtype | No | 0 | 919 | 0.00000 |
| TCGA-DB-A4XD | Male | 32 | Primary | Astrocytoma | III | Mutant | Non-codel | Methylated | No | No | Wildtype | No | 0 | 1210 | 0.00000 |
| TCGA-DB-A4XG | Male | 34 | Primary | Oligodendroglioma | III | Mutant | Codel | Methylated | No | No | Mutant | Yes | 0 | 1850 | 0.00000 |
| TCGA-DB-A64P | Male | 40 | Primary | Oligodendroglioma | III | Mutant | Codel | Methylated | No | No | Mutant | Yes | 0 | 916 | 0.00000 |
| TCGA-DB-A75O | Male | 29 | Primary | Astrocytoma | III | Mutant | Non-codel | Methylated | No | No | [Not available] | No | 0 | 935 | 0.00000 |
| TCGA-DH-5141 | Male | 32 | Primary | Oligodendroglioma | III | Mutant | Codel | Methylated | No | No | Mutant | Yes | 0 | 968 | 0.15500 |
| TCGA-DH-5142 | Male | 29 | Primary | Astrocytoma | III | Mutant | Non-codel | Methylated | No | No | Wildtype | No | 1 | 1627 | 0.08600 |
| TCGA-DH-5143 | Male | 30 | Primary | Oligoastrocytoma | III | Mutant | Non-codel | Methylated | No | No | Wildtype | No | 0 | 1401 | 0.00000 |
| TCGA-DH-A669 | Male | 70 | Primary | Oligodendroglioma | III | Mutant | Codel | Methylated | No | No | Mutant | Yes | 1 | 260 | 0.00000 |
| TCGA-DH-A66B | Male | 52 | Primary | Astrocytoma | III | Mutant | Non-codel | Methylated | No | No | Wildtype | No | 0 | 1279 | 0.00000 |
| TCGA-DH-A7UT | Male | 30 | Primary | Astrocytoma | III | Mutant | Non-codel | Methylated | No | No | [Not available] | No | 1 | 404 | 0.29180 |
| TCGA-DH-A7UU | Male | 43 | Primary | Astrocytoma | III | Mutant | Non-codel | Methylated | No | No | [Not available] | No | 0 | 417 | 0.00000 |
| TCGA-DH-A7UV | Male | 49 | Primary | Astrocytoma | III | Mutant | Non-codel | Methylated | No | No | [Not available] | No | 0 | 566 | 0.00000 |
| TCGA-DU-6393 | Male | 66 | Primary | Oligodendroglioma | III | Mutant | Codel | Methylated | No | No | Mutant | Yes | 0 | 1585 | 0.00000 |
| TCGA-DU-6394 | Male | 53 | Primary | Oligodendroglioma | III | Mutant | Codel | Methylated | No | No | Mutant | No | 1 | 353 | 0.00000 |
| TCGA-DU-6397 | Male | 45 | Primary | Oligodendroglioma | III | Mutant | Codel | Methylated | No | No | Mutant | Yes | 1 | 837 | 0.00000 |
| TCGA-DU-6402 | Male | 52 | Primary | Astrocytoma | III | Wildtype | Non-codel | Unmethylated | Yes | No | Mutant | Yes | 1 | 103 | 1.34210 |
| TCGA-DU-6410 | Male | 56 | Primary | Oligodendroglioma | III | Mutant | Codel | Methylated | No | No | Mutant | Yes | 0 | 242 | 0.00000 |
| TCGA-DU-6542 | Male | 25 | Primary | Oligoastrocytoma | III | Mutant | Non-codel | Methylated | No | No | Wildtype | No | 0 | 73 | 0.00000 |
| TCGA-DU-7013 | Male | 59 | Primary | Astrocytoma | III | Wildtype | Non-codel | Unmethylated | Yes | No | Mutant | Yes | 1 | 187 | 0.00000 |
| TCGA-DU-7292 | Male | 69 | Primary | Astrocytoma | III | Wildtype | Non-codel | Methylated | No | No | Wildtype | No | 1 | 91 | 1.06450 |
| TCGA-DU-7299 | Male | 33 | Primary | Astrocytoma | III | Mutant | Non-codel | Methylated | No | No | Wildtype | No | 1 | 675 | 0.06050 |
| TCGA-DU-7304 | Male | 43 | Primary | Oligoastrocytoma | III | Mutant | Non-codel | Methylated | No | No | Wildtype | No | 1 | 309 | 0.00000 |
| TCGA-DU-8163 | Male | 29 | Primary | Oligoastrocytoma | III | Mutant | Non-codel | Unmethylated | No | No | Wildtype | No | 1 | 509 | 0.08480 |
| TCGA-DU-A5TP | Male | 33 | Primary | Astrocytoma | III | Mutant | Non-codel | Methylated | No | No | Wildtype | No | 1 | 203 | 0.09870 |
| TCGA-DU-A5TT | Male | 70 | Primary | Oligodendroglioma | III | Wildtype | Non-codel | Methylated | No | No | Mutant | Yes | 1 | 584 | 4.17790 |
| TCGA-DU-A76L | Male | 54 | Primary | Oligodendroglioma | III | Wildtype | Non-codel | Methylated | Yes | No | [Not available] | Yes | 1 | 410 | 0.27060 |
| TCGA-DU-A76R | Male | 51 | Primary | Oligodendroglioma | III | Mutant | Codel | Methylated | No | No | [Not available] | Yes | 1 | 338 | 0.08110 |
| TCGA-DU-A7T8 | Male | 35 | Primary | Oligoastrocytoma | III | Mutant | Non-codel | Methylated | No | No | [Not available] | No | 0 | 1850 | 0.00000 |
| TCGA-DU-A7TD | Male | 52 | Primary | Oligoastrocytoma | III | Wildtype | Non-codel | Unmethylated | Yes | No | [Not available] | Yes | 1 | 188 | 0.51040 |
| TCGA-DU-A7TI | Male | 32 | Primary | Astrocytoma | III | [Not available] | Non-codel | Methylated | No | No | [Not available] | No | 1 | 822 | 0.09350 |
| TCGA-DU-A7TJ | Male | 55 | Primary | Astrocytoma | III | Wildtype | Non-codel | Methylated | Yes | No | [Not available] | Yes | 0 | 17 | 2.93840 |
| TCGA-E1-5302 | Male | 41 | Primary | Astrocytoma | III | Mutant | Non-codel | Methylated | No | No | Wildtype | No | 1 | 1242 | 0.38880 |

| | | | | | | | | | | | | | | | |
|---|---|---|---|---|---|---|---|---|---|---|---|---|---|---|---|
| TCGA-E1-5303 | Male | 38 | Primary | Astrocytoma | III | Mutant | Non-codel | Methylated | No | No | Wildtype | No | 1 | 156 | 0.24540 |
| TCGA-E1-5304 | Male | 42 | Primary | Astrocytoma | III | Mutant | Non-codel | Unmethylated | No | No | Wildtype | No | 1 | 903 | 0.00000 |
| TCGA-E1-5305 | Male | 34 | Primary | Astrocytoma | III | Mutant | Non-codel | Methylated | No | No | Wildtype | No | 1 | 1604 | 0.00000 |
| TCGA-E1-5311 | Male | 31 | Primary | Oligodendroglioma | III | Mutant | Codel | Methylated | No | No | Mutant | Yes | 0 | 1850 | 0.00000 |
| TCGA-E1-A7YD | Male | 57 | Primary | Astrocytoma | III | Wildtype | Non-codel | Unmethylated | Yes | No | [Not available] | Yes | 1 | 284 | 3.00950 |
| TCGA-E1-A7YJ | Male | 55 | Primary | Astrocytoma | III | Wildtype | Non-codel | Unmethylated | Yes | Yes | [Not available] | Yes | 1 | 516 | 2.02360 |
| TCGA-E1-A7YK | Male | 52 | Primary | Astrocytoma | III | Mutant | Non-codel | Methylated | No | No | [Not available] | No | 0 | 378 | 0.82170 |
| TCGA-E1-A7YL | Male | 46 | Primary | Astrocytoma | III | Wildtype | Non-codel | Unmethylated | Yes | Yes | [Not available] | Yes | 1 | 91 | 7.26320 |
| TCGA-E1-A7YM | Male | 63 | Primary | Astrocytoma | III | Wildtype | Non-codel | Unmethylated | Yes | No | [Not available] | Yes | 1 | 88 | 3.62120 |
| TCGA-E1-A7YO | Male | 45 | Primary | Oligodendroglioma | III | Mutant | Codel | Methylated | No | No | [Not available] | Yes | 0 | 1850 | 0.00000 |
| TCGA-E1-A7YS | Male | 71 | Primary | Oligodendroglioma | III | Mutant | Codel | Methylated | No | No | [Not available] | Yes | 1 | 466 | 0.00000 |
| TCGA-E1-A7YU | Male | 42 | Primary | Oligoastrocytoma | III | Mutant | Non-codel | Methylated | No | No | [Not available] | No | 1 | 23 | 0.15500 |
| TCGA-FG-5962 | Male | 54 | Primary | Oligodendroglioma | III | Mutant | Codel | Methylated | No | No | Mutant | Yes | 0 | 1453 | 0.00000 |
| TCGA-FG-5963 | Male | 23 | Primary | Astrocytoma | III | Wildtype | Non-codel | Unmethylated | No | No | Wildtype | No | 1 | 395 | 0.50500 |
| TCGA-FG-6692 | Male | 63 | Primary | Oligodendroglioma | III | Wildtype | Non-codel | Methylated | Yes | No | Mutant | Yes | 1 | 561 | 0.34210 |
| TCGA-FG-7636 | Male | 48 | Primary | Astrocytoma | III | Mutant | Non-codel | Methylated | No | No | Wildtype | No | 0 | 544 | 0.00000 |
| TCGA-FG-8181 | Male | 23 | Primary | Oligoastrocytoma | III | Wildtype | Non-codel | Unmethylated | No | No | Wildtype | No | 0 | 862 | 0.00000 |
| TCGA-FG-8185 | Male | 37 | Primary | Astrocytoma | III | Mutant | Non-codel | Methylated | No | No | Wildtype | No | 0 | 433 | 0.05850 |
| TCGA-FG-8191 | Male | 30 | Primary | Oligodendroglioma | III | Mutant | Non-codel | Unmethylated | No | No | Wildtype | No | 0 | 992 | 0.07450 |
| TCGA-FG-A4MU | Male | 58 | Primary | Oligoastrocytoma | III | Wildtype | Non-codel | Methylated | Yes | Yes | Mutant | Yes | 0 | 326 | 2.16260 |
| TCGA-FG-A4MW | Male | 63 | Primary | Oligoastrocytoma | III | Wildtype | Non-codel | Methylated | Yes | No | Mutant | Yes | 1 | 226 | 0.00000 |
| TCGA-FN-7833 | Male | 25 | Primary | Oligoastrocytoma | III | Mutant | Non-codel | Methylated | No | No | Wildtype | No | 0 | 837 | 0.00000 |
| TCGA-HT-7468 | Male | 30 | Primary | Oligodendroglioma | III | Mutant | Codel | Methylated | No | No | Mutant | Yes | 0 | 203 | 0.00000 |
| TCGA-HT-7469 | Male | 30 | Primary | Oligodendroglioma | III | Wildtype | Non-codel | Methylated | No | No | Wildtype | No | 1 | 92 | 0.09610 |
| TCGA-HT-7470 | Male | 37 | Primary | Oligodendroglioma | III | Mutant | Non-codel | Methylated | No | No | Wildtype | No | 0 | 1220 | 0.06040 |
| TCGA-HT-7475 | Male | 67 | Primary | Oligoastrocytoma | III | Mutant | Non-codel | Methylated | No | No | Wildtype | No | 0 | 530 | 0.00000 |
| TCGA-HT-7477 | Male | 62 | Primary | Astrocytoma | III | Mutant | Non-codel | Methylated | No | No | Wildtype | No | 0 | 738 | 0.19010 |
| TCGA-HT-7479 | Male | 44 | Primary | Astrocytoma | III | Mutant | Non-codel | Methylated | No | No | Mutant | No | 1 | 705 | 0.13160 |
| TCGA-HT-7609 | Male | 34 | Primary | Oligoastrocytoma | III | Mutant | Non-codel | Methylated | No | No | Wildtype | No | 0 | 1399 | 0.00000 |
| TCGA-HT-7616 | Male | 75 | Primary | Oligodendroglioma | III | Mutant | Codel | Methylated | No | No | Mutant | No | 1 | 7 | 0.00000 |
| TCGA-HT-7620 | Male | 40 | Primary | Oligodendroglioma | III | Mutant | Codel | Methylated | No | No | Mutant | Yes | 1 | 280 | 0.00000 |
| TCGA-HT-7677 | Male | 53 | Primary | Oligodendroglioma | III | Mutant | Codel | Methylated | No | No | Mutant | Yes | 0 | 494 | 0.07770 |
| TCGA-HT-7684 | Male | 58 | Primary | Oligoastrocytoma | III | Mutant | Non-codel | Methylated | No | No | Mutant | Yes | 0 | 184 | 0.07180 |
| TCGA-HT-7687 | Male | 74 | Primary | Oligodendroglioma | III | Mutant | Codel | Methylated | No | No | Mutant | Yes | 0 | 3 | 0.08250 |
| TCGA-HT-7688 | Male | 59 | Primary | Oligodendroglioma | III | Mutant | Non-codel | Methylated | No | No | Wildtype | No | 0 | 964 | 0.00000 |
| TCGA-HT-7690 | Male | 29 | Primary | Oligoastrocytoma | III | Mutant | Non-codel | Methylated | No | No | Wildtype | No | 0 | 3 | 0.07070 |
| TCGA-HT-7694 | Male | 60 | Primary | Oligodendroglioma | III | Mutant | Codel | Methylated | No | No | Mutant | No | 0 | 210 | 0.06850 |
| TCGA-HT-7855 | Male | 39 | Primary | Astrocytoma | III | Mutant | Non-codel | Methylated | No | No | Wildtype | No | 0 | 585 | 0.00000 |
| TCGA-HT-7856 | Male | 35 | Primary | Oligodendroglioma | III | Mutant | Codel | Methylated | No | No | Mutant | Yes | 0 | 1189 | 0.22290 |
| TCGA-HT-7879 | Male | 31 | Primary | Oligoastrocytoma | III | Mutant | Non-codel | Methylated | No | No | Wildtype | No | 0 | 112 | 1.54770 |
| TCGA-HT-7882 | Male | 66 | Primary | Oligodendroglioma | III | Wildtype | Non-codel | Methylated | No | No | Mutant | Yes | 1 | 113 | 0.24960 |
| TCGA-HT-8011 | Male | 55 | Primary | Astrocytoma | III | Wildtype | Non-codel | Unmethylated | Yes | No | Mutant | Yes | 1 | 278 | 0.47310 |
| TCGA-HT-8105 | Male | 54 | Primary | Oligodendroglioma | III | Mutant | Codel | Methylated | No | No | Mutant | Yes | 0 | 190 | 0.07310 |
| TCGA-HT-8106 | Male | 53 | Primary | Astrocytoma | III | Mutant | Non-codel | Methylated | No | No | Wildtype | No | 0 | 3 | 0.00000 |
| TCGA-HT-8109 | Male | 64 | Primary | Oligodendroglioma | III | Mutant | Codel | Methylated | No | No | Mutant | Yes | 0 | 169 | 0.07150 |
| TCGA-HT-8110 | Male | 57 | Primary | Astrocytoma | III | Wildtype | Non-codel | Methylated | Yes | No | Mutant | Yes | 0 | 419 | 3.26010 |
| TCGA-HT-8111 | Male | 32 | Primary | Oligoastrocytoma | III | Mutant | Non-codel | Methylated | No | No | Wildtype | No | 0 | 7 | 0.37460 |
| TCGA-HT-8114 | Male | 36 | Primary | Oligoastrocytoma | III | Mutant | Non-codel | Methylated | No | No | Wildtype | No | 0 | 1040 | 0.14740 |
| TCGA-HT-8564 | Male | 47 | Primary | Astrocytoma | III | Wildtype | Non-codel | Unmethylated | No | No | Wildtype | No | 1 | 120 | 0.41220 |
| TCGA-HT-A61B | Male | 22 | Primary | Astrocytoma | III | Mutant | Non-codel | Methylated | No | No | Wildtype | Yes | 0 | 533 | 0.00000 |

| | | | | | | | | | | | | | | | |
|---|---|---|---|---|---|---|---|---|---|---|---|---|---|---|---|
| TCGA-HT-A61C | Male | 66 | Primary | Oligodendroglioma | III | Wildtype | Non-codel | Unmethylated | Yes | No | Mutant | Yes | 1 | 476 | 0.17640 |
| TCGA-HT-A74H | Male | 62 | Primary | Astrocytoma | III | Wildtype | Non-codel | Unmethylated | Yes | No | [Not available] | No | 0 | 74 | 0.13070 |
| TCGA-HT-A74O | Male | 34 | Primary | Astrocytoma | III | Mutant | Non-codel | Methylated | No | No | [Not available] | No | 0 | 3 | 0.14440 |
| TCGA-HW-8320 | Male | 36 | Primary | Astrocytoma | III | Mutant | Non-codel | Methylated | No | No | Wildtype | No | 0 | 1217 | 0.00000 |
| TCGA-HW-8321 | Male | 31 | Primary | Astrocytoma | III | Mutant | Non-codel | Methylated | No | No | Wildtype | No | 0 | 1294 | 0.06570 |
| TCGA-HW-A5KJ | Male | 68 | Primary | Oligodendroglioma | III | Mutant | Codel | Methylated | No | No | Mutant | Yes | 1 | 615 | 0.00000 |
| TCGA-HW-A5KK | Male | 64 | Primary | Astrocytoma | III | Wildtype | Non-codel | Methylated | No | No | Mutant | Yes | 1 | 388 | 2.83990 |
| TCGA-IK-8125 | Male | 62 | Primary | Oligoastrocytoma | III | Mutant | Codel | Methylated | No | No | Mutant | Yes | 0 | 1301 | 0.05480 |
| TCGA-KT-A74X | Male | 26 | Primary | Oligoastrocytoma | III | Mutant | Codel | Methylated | No | No | [Not available] | Yes | 0 | 438 | 0.00000 |
| TCGA-P5-A5EU | Male | 35 | Primary | Astrocytoma | III | Mutant | Non-codel | Unmethylated | No | No | Wildtype | No | 0 | 0 | 0.00000 |
| TCGA-P5-A5EZ | Male | 39 | Primary | Astrocytoma | III | Mutant | Non-codel | Methylated | No | No | Wildtype | No | 0 | 70 | 0.27710 |
| TCGA-P5-A72W | Male | 35 | Primary | Astrocytoma | III | Mutant | Non-codel | Unmethylated | No | No | [Not available] | No | 0 | 317 | 0.00000 |
| TCGA-P5-A72X | Male | 21 | Primary | Astrocytoma | III | Mutant | Non-codel | Methylated | No | No | [Not available] | No | 0 | 403 | 0.07200 |
| TCGA-P5-A730 | Male | 22 | Primary | Oligoastrocytoma | III | Mutant | Codel | Methylated | No | No | [Not available] | Yes | 0 | 333 | 0.00000 |
| TCGA-QH-A6CS | Male | 41 | Primary | Astrocytoma | III | Wildtype | Non-codel | Unmethylated | Yes | No | [Not available] | [Not available] | 1 | 498 | 1.89710 |
| TCGA-QH-A6CV | Male | 51 | Primary | Oligoastrocytoma | III | Wildtype | Non-codel | Unmethylated | Yes | Yes | [Not available] | Yes | 0 | 442 | 0.08790 |
| TCGA-QH-A6CW | Male | 43 | Primary | Oligoastrocytoma | III | Mutant | Non-codel | Methylated | No | No | [Not available] | No | 0 | 414 | 0.37660 |
| TCGA-QH-A6CY | Male | 38 | Primary | Oligoastrocytoma | III | Mutant | Codel | Methylated | No | No | [Not available] | Yes | 0 | 408 | 0.00000 |
| TCGA-QH-A6X4 | Male | 47 | Primary | Oligoastrocytoma | III | Mutant | Codel | Methylated | No | No | [Not available] | Yes | 0 | 442 | 0.07330 |
| TCGA-QH-A6XC | Male | 48 | Primary | Astrocytoma | III | Wildtype | Non-codel | Methylated | Yes | Yes | [Not available] | Yes | 1 | 438 | 3.60280 |
| TCGA-R8-A6ML | Male | 52 | Primary | Oligodendroglioma | III | Mutant | Codel | Methylated | No | No | [Not available] | Yes | 0 | 1850 | 0.07490 |
| TCGA-RY-A840 | Male | 47 | Primary | Oligodendroglioma | III | Mutant | Codel | Methylated | No | No | [Not available] | Yes | 0 | 854 | 0.00000 |
| TCGA-RY-A843 | Male | 30 | Primary | Astrocytoma | III | Mutant | Non-codel | Methylated | No | No | [Not available] | Yes | 0 | 63 | 0.05850 |
| TCGA-S9-A6TV | Male | 50 | Primary | Oligoastrocytoma | III | Mutant | Non-codel | Methylated | No | No | [Not available] | No | 0 | 571 | 0.06740 |
| TCGA-S9-A6TW | Male | 40 | Primary | Oligodendroglioma | III | Mutant | Codel | Methylated | No | No | [Not available] | Yes | 1 | 861 | 0.00000 |
| TCGA-S9-A6TX | Male | 46 | Primary | Oligodendroglioma | III | Mutant | Codel | Methylated | No | No | [Not available] | Yes | 1 | 856 | 0.06980 |
| TCGA-S9-A6U0 | Male | 46 | Primary | Astrocytoma | III | Wildtype | Non-codel | Methylated | Yes | No | [Not available] | Yes | 1 | 692 | 0.33240 |
| TCGA-S9-A6U6 | Male | 28 | Primary | Astrocytoma | III | Mutant | Non-codel | Methylated | No | No | [Not available] | No | 0 | 1069 | 0.16480 |
| TCGA-S9-A6U9 | Male | 36 | Primary | Astrocytoma | III | Mutant | Non-codel | Methylated | No | No | [Not available] | No | 0 | 1850 | 0.07710 |
| TCGA-S9-A6UA | Male | 66 | Primary | Astrocytoma | III | Wildtype | Non-codel | Methylated | No | No | [Not available] | No | 1 | 161 | 1.33270 |
| TCGA-S9-A6WD | Male | 58 | Primary | Oligodendroglioma | III | Mutant | Codel | Methylated | No | No | [Not available] | Yes | 0 | 1850 | 0.29300 |
| TCGA-S9-A6WG | Male | 31 | Primary | Astrocytoma | III | Mutant | Non-codel | Methylated | No | No | [Not available] | No | 0 | 1850 | 0.00000 |
| TCGA-S9-A6WL | Male | 52 | Primary | Astrocytoma | III | Mutant | Codel | Methylated | No | No | [Not available] | Yes | 1 | 240 | 0.00000 |
| TCGA-S9-A6WP | Male | 42 | Primary | Oligoastrocytoma | III | Mutant | Codel | Methylated | No | No | [Not available] | No | 0 | 567 | 0.60230 |
| TCGA-S9-A7IX | Male | 57 | Primary | Astrocytoma | III | Wildtype | Non-codel | Unmethylated | Yes | No | [Not available] | No | 1 | 350 | 3.33330 |
| TCGA-S9-A7IY | Male | 39 | Primary | Oligoastrocytoma | III | Mutant | Codel | Methylated | No | No | [Not available] | Yes | 0 | 715 | 0.08820 |
| TCGA-S9-A7J2 | Male | 25 | Primary | Oligodendroglioma | III | Mutant | Codel | Methylated | No | No | [Not available] | Yes | 0 | 62 | 0.08100 |
| TCGA-S9-A7R2 | Male | 69 | Primary | Astrocytoma | III | Wildtype | Non-codel | Unmethylated | Yes | No | [Not available] | Yes | 1 | 316 | 6.54170 |
| TCGA-S9-A7R4 | Male | 46 | Primary | Astrocytoma | III | Mutant | Non-codel | Methylated | No | No | [Not available] | No | 0 | 914 | 0.00000 |
| TCGA-S9-A89V | Male | 70 | Primary | Astrocytoma | III | Wildtype | Non-codel | Methylated | No | No | [Not available] | No | 1 | 129 | 0.10410 |
| TCGA-S9-A89Z | Male | 40 | Primary | Astrocytoma | III | Mutant | Non-codel | Methylated | No | No | [Not available] | No | 0 | 623 | 0.31820 |
| TCGA-TM-A84B | Male | 40 | Primary | Astrocytoma | III | Wildtype | Non-codel | Unmethylated | Yes | No | [Not available] | Yes | 1 | 758 | 14.50060 |
| TCGA-TM-A84F | Male | 48 | Primary | Astrocytoma | III | Mutant | Non-codel | Methylated | No | No | [Not available] | Yes | 0 | 1796 | 0.37140 |
| TCGA-TM-A84I | Male | 30 | Primary | Astrocytoma | III | Mutant | Non-codel | Methylated | No | No | [Not available] | No | 0 | 854 | 0.00000 |
| TCGA-TM-A84J | Male | 63 | Primary | Oligodendroglioma | III | Wildtype | Non-codel | Unmethylated | No | No | [Not available] | Yes | 1 | 454 | 0.00000 |
| TCGA-TM-A84M | Male | 40 | Primary | Oligodendroglioma | III | Mutant | Codel | Methylated | No | No | [Not available] | Yes | 0 | 754 | 0.00000 |
| TCGA-TM-A84S | Male | 36 | Primary | Oligodendroglioma | III | Mutant | Codel | Methylated | No | No | [Not available] | Yes | 0 | 454 | 1.14600 |
| TCGA-VM-A8CB | Male | 33 | Primary | Oligodendroglioma | III | Mutant | Codel | Methylated | No | No | [Not available] | Yes | 0 | 3 | 0.00000 |
| TCGA-VM-A8CD | Male | 58 | Primary | Astrocytoma | III | Wildtype | Non-codel | Unmethylated | No | No | [Not available] | Yes | 1 | 240 | 1.87400 |
| TCGA-VV-A829 | Male | 44 | Primary | Oligoastrocytoma | III | Mutant | Codel | Methylated | No | No | [Not available] | Yes | 0 | 1127 | 0.00000 |

| | | | | | | | | | | | | | | | |
|---|---|---|---|---|---|---|---|---|---|---|---|---|---|---|---|
| TCGA-VW-A8FI | Male | 66 | Primary | Astrocytoma | III | Wildtype | Non-codel | Unmethylated | Yes | No | [Not available] | Yes | 1 | 245 | 3.11520 |

**Table S3. HGG patients from the CGGA cohort.**

| Patient id | Gender | Age | Tumor type | Histology | Grade | IDH status | Codeletion 1p 19q | MGMT status | OS status | OS time | SRY expression |
|---|---|---|---|---|---|---|---|---|---|---|---|
| CGGA_1001 | Male | 11 | Primary | Glioblastoma | IV | Wildtype | Non-codel | Unmethylated | 0 | 1850 | 0.00000 |
| CGGA_1006 | Male | 42 | Primary | Astrocytoma | III | Wildtype | Non-codel | Unmethylated | 1 | 254 | 2.04749 |
| CGGA_1015 | Male | 62 | Primary | Glioblastoma | IV | Wildtype | Non-codel | Unmethylated | 1 | 164 | 0.00000 |
| CGGA_1024 | Male | 64 | Primary | Glioblastoma | IV | Wildtype | Non-codel | Methylated | 0 | 1850 | 0.18886 |
| CGGA_1026 | Male | 57 | Primary | Glioblastoma | IV | Wildtype | Non-codel | Methylated | 1 | 1570 | 3.32560 |
| CGGA_1027 | Male | 37 | Primary | Oligodendroglioma | III | Mutant | Codel | Methylated | 0 | 1850 | 0.00000 |
| CGGA_103 | Male | 56 | Primary | Astrocytoma | III | Mutant | Non-codel | Unmethylated | 1 | 1188 | 0.00000 |
| CGGA_1034 | Male | 23 | Primary | Astrocytoma | III | Wildtype | Non-codel | Unmethylated | 1 | 753 | 0.00000 |
| CGGA_1036 | Male | 41 | Primary | Glioblastoma | IV | Wildtype | Non-codel | Methylated | 1 | 806 | 0.52828 |
| CGGA_1039 | Male | 40 | Primary | Glioblastoma | IV | Wildtype | Non-codel | Methylated | 1 | 727 | 0.22368 |
| CGGA_1041 | Male | 58 | Primary | Glioblastoma | IV | Wildtype | Non-codel | Unmethylated | 0 | 1850 | 0.12558 |
| CGGA_1045 | Male | 79 | Primary | Glioblastoma | IV | Wildtype | Non-codel | Unmethylated | 1 | 348 | 2.42109 |
| CGGA_1049 | Male | 58 | Primary | Glioblastoma | IV | Wildtype | Non-codel | Unmethylated | 1 | 809 | 0.00000 |
| CGGA_1055 | Male | 43 | Primary | Astrocytoma | III | Mutant | Non-codel | Unmethylated | 1 | 1611 | 0.00000 |
| CGGA_1070 | Male | 37 | Primary | Glioblastoma | IV | Wildtype | Non-codel | Unmethylated | 1 | 333 | 0.00000 |
| CGGA_1072 | Male | 26 | Primary | Glioblastoma | IV | Wildtype | Non-codel | Unmethylated | 1 | 1090 | 0.00000 |
| CGGA_1074 | Male | 50 | Primary | Glioblastoma | IV | Wildtype | Non-codel | Methylated | 1 | 86 | 8.20683 |
| CGGA_1075 | Male | 72 | Primary | Glioblastoma | IV | Wildtype | Non-codel | Unmethylated | 1 | 398 | 0.35789 |
| CGGA_1077 | Male | 47 | Primary | Glioblastoma | IV | Wildtype | Non-codel | Unmethylated | 1 | 591 | 0.00000 |
| CGGA_1079 | Male | 38 | Primary | Astrocytoma | III | Wildtype | Non-codel | Unmethylated | 1 | 471 | 2.71559 |
| CGGA_1083 | Male | 59 | Primary | Glioblastoma | IV | Wildtype | Non-codel | Unmethylated | 1 | 450 | 1.77043 |
| CGGA_1106 | Male | 37 | Primary | Glioblastoma | IV | Wildtype | Non-codel | Methylated | 1 | 420 | 0.00000 |
| CGGA_1109 | Male | 42 | Primary | Glioblastoma | IV | Wildtype | Non-codel | Unmethylated | 1 | 1348 | 0.00000 |
| CGGA_1111 | Male | 45 | Primary | Astrocytoma | III | Mutant | Non-codel | Methylated | 0 | 1850 | 0.11171 |
| CGGA_1113 | Male | 34 | Primary | Oligodendroglioma | III | Mutant | Codel | Methylated | 1 | 1265 | 0.00000 |
| CGGA_1114 | Male | 53 | Primary | Glioblastoma | IV | Wildtype | Non-codel | Unmethylated | 1 | 460 | 0.60142 |
| CGGA_1124 | Male | 47 | Primary | Glioblastoma | IV | Wildtype | Non-codel | Unmethylated | 1 | 484 | 2.07371 |
| CGGA_1126 | Male | 44 | Primary | Astrocytoma | III | Mutant | Non-codel | Methylated | 1 | 784 | 0.00000 |
| CGGA_1135 | Male | 40 | Primary | Glioblastoma | IV | Wildtype | Non-codel | Unmethylated | 1 | 1244 | 0.27346 |
| CGGA_1137 | Male | 58 | Primary | Astrocytoma | III | Wildtype | [Not available] | Methylated | 0 | 1850 | 0.00000 |
| CGGA_1138 | Male | 54 | Primary | Glioblastoma | IV | Wildtype | Non-codel | Unmethylated | 1 | 411 | 0.00000 |
| CGGA_1141 | Male | 65 | Primary | Astrocytoma | III | Wildtype | Non-codel | Unmethylated | 1 | 432 | 0.06718 |
| CGGA_1142 | Male | 60 | Primary | Glioblastoma | IV | Wildtype | Non-codel | Unmethylated | 1 | 1005 | 0.00000 |

| | | | | | | | | | | | |
|---|---|---|---|---|---|---|---|---|---|---|---|
| CGGA_1146 | Male | 35 | Primary | Astrocytoma | III | Mutant | Non-codel | Methylated | 0 | 1850 | 3.73041 |
| CGGA_1155 | Male | 43 | Primary | Astrocytoma | III | Mutant | Codel | Methylated | 1 | 1373 | 0.34654 |
| CGGA_1161 | Male | 36 | Primary | Astrocytoma | III | Mutant | Non-codel | Unmethylated | 0 | 1850 | 0.00000 |
| CGGA_1162 | Male | 39 | Primary | Oligoastrocytoma | III | [Not available] | Codel | Unmethylated | 0 | 1850 | 0.00000 |
| CGGA_1169 | Male | 30 | Primary | Astrocytoma | III | Mutant | Non-codel | Unmethylated | 1 | 1765 | 0.09924 |
| CGGA_1171 | Male | 40 | Primary | Glioblastoma | IV | Wildtype | Non-codel | Methylated | 1 | 412 | 1.24893 |
| CGGA_1191 | Male | 53 | Primary | Oligodendroglioma | III | Mutant | Codel | Methylated | 0 | 1850 | 0.00000 |
| CGGA_1195 | Male | 43 | Primary | Astrocytoma | III | Mutant | Non-codel | Unmethylated | 1 | 476 | 4.40175 |
| CGGA_1212 | Male | 31 | Primary | Astrocytoma | III | Mutant | Non-codel | Methylated | 0 | 1850 | 0.15158 |
| CGGA_1216 | Male | 44 | Primary | Glioblastoma | IV | Wildtype | Non-codel | Methylated | 1 | 312 | 2.82780 |
| CGGA_1223 | Male | 42 | Primary | Astrocytoma | III | Wildtype | Non-codel | Unmethylated | 1 | 682 | 3.08919 |
| CGGA_1224 | Male | 57 | Primary | Glioblastoma | IV | Wildtype | Non-codel | Unmethylated | 1 | 271 | 1.21767 |
| CGGA_1232 | Male | 39 | Primary | Astrocytoma | III | Mutant | Non-codel | Methylated | 0 | 1850 | 2.36398 |
| CGGA_1234 | Male | 32 | Primary | Glioblastoma | IV | Wildtype | Non-codel | Unmethylated | 1 | 705 | 2.81023 |
| CGGA_1237 | Male | 62 | Primary | Glioblastoma | IV | Wildtype | Non-codel | Unmethylated | 1 | 296 | 4.01197 |
| CGGA_1240 | Male | 33 | Primary | Glioblastoma | IV | Wildtype | Non-codel | Unmethylated | 1 | 19 | 1.08215 |
| CGGA_1246 | Male | 53 | Primary | Oligodendroglioma | III | Mutant | Codel | Methylated | 0 | 1850 | 0.00000 |
| CGGA_1258 | Male | 42 | Primary | Glioblastoma | IV | Wildtype | Non-codel | Unmethylated | 1 | 114 | 10.70848 |
| CGGA_1263 | Male | 50 | Primary | Astrocytoma | III | Wildtype | Non-codel | Unmethylated | 1 | 772 | 0.31014 |
| CGGA_1275 | Male | 70 | Primary | Glioblastoma | IV | Wildtype | Non-codel | Unmethylated | 1 | 183 | 0.00000 |
| CGGA_1280 | Male | 74 | Primary | Astrocytoma | III | Wildtype | Non-codel | Unmethylated | 1 | 447 | 4.93628 |
| CGGA_1281 | Male | 45 | Primary | Astrocytoma | III | Wildtype | Codel | Methylated | 0 | 1850 | 0.00000 |
| CGGA_1284 | Male | 73 | Primary | Astrocytoma | III | Wildtype | Non-codel | Unmethylated | 1 | 97 | 0.00000 |
| CGGA_1287 | Male | 29 | Primary | Astrocytoma | IV | Mutant | Non-codel | Unmethylated | 1 | 533 | 0.00000 |
| CGGA_1291 | Male | 41 | Primary | Oligodendroglioma | III | [Not available] | Codel | Unmethylated | 0 | 1850 | 0.00000 |
| CGGA_1292 | Male | 17 | Primary | Astrocytoma | III | Wildtype | Non-codel | Methylated | 0 | 1850 | 0.00000 |
| CGGA_1299 | Male | 54 | Primary | Glioblastoma | IV | Wildtype | Non-codel | Unmethylated | 1 | 550 | 0.00000 |
| CGGA_1300 | Male | 35 | Primary | Astrocytoma | III | Mutant | Non-codel | Methylated | 1 | 1793 | 0.65550 |
| CGGA_1305 | Male | 43 | Primary | Astrocytoma | III | Mutant | Non-codel | Methylated | 0 | 1850 | 0.10195 |
| CGGA_1309 | Male | 65 | Primary | Oligodendroglioma | III | [Not available] | Codel | Methylated | 0 | 964 | 0.00000 |
| CGGA_1313 | Male | 42 | Primary | Glioblastoma | IV | Wildtype | Non-codel | Unmethylated | 1 | 379 | 0.87382 |
| CGGA_1320 | Male | 19 | Primary | Glioblastoma | IV | Wildtype | Non-codel | Unmethylated | 1 | 59 | 44.48827 |
| CGGA_1321 | Male | 46 | Primary | Astrocytoma | III | Mutant | Non-codel | Methylated | 1 | 702 | 0.00000 |
| CGGA_1322 | Male | 10 | Primary | Astrocytoma | III | Mutant | Non-codel | Methylated | 1 | 253 | 0.00000 |
| CGGA_1326 | Male | 45 | Primary | Astrocytoma | IV | Mutant | Non-codel | Unmethylated | 1 | 322 | 2.02871 |
| CGGA_1340 | Male | 35 | Primary | Astrocytoma | III | Wildtype | Non-codel | Unmethylated | 1 | 1109 | 0.00000 |

| | | | | | | | | | | | |
|---|---|---|---|---|---|---|---|---|---|---|---|
| CGGA_1342 | Male | 47 | Primary | Glioblastoma | IV | Wildtype | Non-codel | Unmethylated | 1 | 186 | 1.46900 |
| CGGA_1353 | Male | 65 | Primary | Glioblastoma | IV | Wildtype | Non-codel | Unmethylated | 1 | 1022 | 0.00000 |
| CGGA_1365 | Male | 55 | Primary | Glioblastoma | IV | Wildtype | [Not available] | Methylated | 1 | 253 | 0.00000 |
| CGGA_1371 | Male | 68 | Primary | Glioblastoma | IV | Wildtype | Non-codel | Methylated | 0 | 1850 | 0.00000 |
| CGGA_1377 | Male | 28 | Primary | Astrocytoma | III | Mutant | Non-codel | Methylated | 0 | 1850 | 0.00000 |
| CGGA_1378 | Male | 47 | Primary | Glioblastoma | IV | Wildtype | [Not available] | Unmethylated | 1 | 378 | 0.00000 |
| CGGA_1380 | Male | 46 | Primary | Glioblastoma | IV | Wildtype | Non-codel | Methylated | 1 | 291 | 3.86675 |
| CGGA_1382 | Male | 57 | Primary | Glioblastoma | IV | Wildtype | Non-codel | Methylated | 1 | 284 | 0.27722 |
| CGGA_1386 | Male | 34 | Primary | Oligodendroglioma | III | [Not available] | Codel | Methylated | 0 | 1850 | 0.00000 |
| CGGA_139 | Male | 59 | Primary | Astrocytoma | IV | Mutant | Non-codel | Unmethylated | 1 | 694 | 0.00000 |
| CGGA_1391 | Male | 62 | Primary | Glioblastoma | IV | Wildtype | Non-codel | Unmethylated | 0 | 426 | 3.94185 |
| CGGA_1392 | Male | 62 | Primary | Glioblastoma | IV | Wildtype | Non-codel | Unmethylated | 1 | 473 | 0.91962 |
| CGGA_1401 | Male | 39 | Primary | Astrocytoma | III | Mutant | Non-codel | Methylated | 1 | 1680 | 0.13052 |
| CGGA_1402 | Male | 30 | Primary | Glioblastoma | IV | Wildtype | Non-codel | Methylated | 0 | 1850 | 2.71820 |
| CGGA_1413 | Male | 35 | Primary | Oligodendroglioma | III | Mutant | Codel | Methylated | 0 | 1850 | 0.50666 |
| CGGA_1416 | Male | 42 | Primary | Oligodendroglioma | III | Mutant | Codel | Unmethylated | 0 | 1850 | 0.00000 |
| CGGA_1417 | Male | 56 | Primary | Oligodendroglioma | III | Mutant | Codel | Methylated | 0 | 1850 | 0.00000 |
| CGGA_1420 | Male | 60 | Primary | Glioblastoma | IV | Wildtype | Non-codel | Methylated | 1 | 364 | 0.00000 |
| CGGA_1421 | Male | 35 | Primary | Oligodendroglioma | III | [Not available] | Non-codel | Methylated | 0 | 1850 | 0.00000 |
| CGGA_1422 | Male | 76 | Primary | Glioblastoma | IV | Wildtype | [Not available] | Unmethylated | 1 | 204 | 0.00000 |
| CGGA_1431 | Male | 48 | Primary | Astrocytoma | III | Mutant | Non-codel | Methylated | 0 | 1850 | 0.00000 |
| CGGA_1441 | Male | 70 | Primary | Glioblastoma | IV | Wildtype | Non-codel | Methylated | 0 | 1850 | 1.25536 |
| CGGA_1452 | Male | 53 | Primary | Glioblastoma | IV | Wildtype | Non-codel | Methylated | 1 | 468 | 3.33621 |
| CGGA_1457 | Male | 60 | Primary | Glioblastoma | IV | Wildtype | Non-codel | Methylated | 1 | 312 | 1.37511 |
| CGGA_1462 | Male | 49 | Primary | Glioblastoma | IV | Wildtype | Non-codel | Methylated | 1 | 174 | 1.58244 |
| CGGA_1467 | Male | 58 | Primary | Astrocytoma | IV | Mutant | [Not available] | Unmethylated | 1 | 866 | 0.00000 |
| CGGA_1469 | Male | 46 | Primary | Oligoastrocytoma | III | Mutant | [Not available] | Methylated | 0 | 1850 | 0.00000 |
| CGGA_1472 | Male | 34 | Primary | [Not available] | IV | [Not available] | Non-codel | Methylated | 1 | 1025 | 2.06704 |
| CGGA_1481 | Male | 55 | Primary | Glioblastoma | IV | Wildtype | [Not available] | Methylated | 1 | 131 | 0.00000 |
| CGGA_1482 | Male | 38 | Primary | Oligodendroglioma | III | [Not available] | Codel | Methylated | 0 | 1850 | 0.00000 |
| CGGA_1486 | Male | 45 | Primary | Glioblastoma | IV | Wildtype | [Not available] | Methylated | 1 | 184 | 0.00000 |
| CGGA_1491 | Male | 29 | Primary | Astrocytoma | IV | Mutant | Non-codel | Unmethylated | 1 | 246 | 0.11918 |
| CGGA_1494 | Male | 21 | Primary | Glioblastoma | IV | Wildtype | [Not available] | Methylated | 1 | 269 | 0.00000 |
| CGGA_1501 | Male | 58 | Primary | [Not available] | IV | [Not available] | Non-codel | Methylated | 1 | 222 | 0.00000 |
| CGGA_1503 | Male | 47 | Primary | Glioblastoma | IV | Wildtype | [Not available] | Unmethylated | 1 | 777 | 0.01672 |
| CGGA_1508 | Male | 39 | Primary | Astrocytoma | III | Mutant | Non-codel | Methylated | 1 | 1049 | 0.00000 |

| | | | | | | | | | | | |
|---|---|---|---|---|---|---|---|---|---|---|---|
| CGGA_1525 | Male | 41 | Primary | Oligoastrocytoma | III | [Not available] | Non-codel | Methylated | 1 | 1686 | 0.16079 |
| CGGA_1526 | Male | 39 | Primary | Oligodendroglioma | III | Mutant | Codel | Methylated | 0 | 1850 | 0.25089 |
| CGGA_1527 | Male | 47 | Primary | Astrocytoma | III | Mutant | Non-codel | Methylated | 1 | 1714 | 0.00000 |
| CGGA_1529 | Male | 63 | Primary | Glioblastoma | IV | Wildtype | Non-codel | Unmethylated | 1 | 583 | 0.00000 |
| CGGA_1530 | Male | 41 | Primary | Oligoastrocytoma | III | [Not available] | Non-codel | Unmethylated | 0 | 1850 | 0.00000 |
| CGGA_1537 | Male | 73 | Primary | Glioblastoma | IV | Wildtype | Non-codel | Methylated | 1 | 97 | 1.89079 |
| CGGA_1539 | Male | 61 | Primary | Astrocytoma | IV | Mutant | Non-codel | Methylated | 0 | 1850 | 0.00000 |
| CGGA_1542 | Male | 26 | Primary | Astrocytoma | IV | Mutant | [Not available] | Unmethylated | 0 | 184 | 0.00000 |
| CGGA_1543 | Male | 57 | Primary | Astrocytoma | IV | Mutant | Non-codel | Unmethylated | 1 | 723 | 3.01250 |
| CGGA_1546 | Male | 56 | Primary | Glioblastoma | IV | Wildtype | Non-codel | Unmethylated | 1 | 223 | 0.23099 |
| CGGA_1548 | Male | 53 | Primary | Glioblastoma | IV | Wildtype | Non-codel | Unmethylated | 1 | 1054 | 3.82261 |
| CGGA_1552 | Male | 39 | Primary | Oligodendroglioma | III | Mutant | Codel | Methylated | 0 | 1850 | 0.00000 |
| CGGA_1557 | Male | 47 | Primary | Oligodendroglioma | III | Mutant | Codel | Methylated | 0 | 1850 | 0.00000 |
| CGGA_1559 | Male | 63 | Primary | Astrocytoma | IV | Mutant | Codel | Methylated | 1 | 603 | 0.00000 |
| CGGA_1562 | Male | 51 | Primary | Oligodendroglioma | III | Mutant | Codel | Methylated | 0 | 1850 | 0.00000 |
| CGGA_1564 | Male | 48 | Primary | Glioblastoma | IV | Wildtype | Non-codel | Unmethylated | 1 | 190 | 2.61443 |
| CGGA_1596 | Male | 63 | Primary | Glioblastoma | IV | Wildtype | Non-codel | Methylated | 1 | 205 | 0.17608 |
| CGGA_1597 | Male | 58 | Primary | Glioblastoma | IV | Wildtype | Non-codel | Methylated | 1 | 174 | 1.79509 |
| CGGA_1598 | Male | 31 | Primary | Astrocytoma | III | Wildtype | Non-codel | Unmethylated | 1 | 564 | 0.36205 |
| CGGA_1601 | Male | 66 | Primary | Glioblastoma | IV | Wildtype | Non-codel | Methylated | 1 | 710 | 0.12347 |
| CGGA_1612 | Male | 68 | Primary | Glioblastoma | IV | Wildtype | Non-codel | Unmethylated | 1 | 718 | 4.75090 |
| CGGA_1613 | Male | 53 | Primary | Glioblastoma | IV | Wildtype | Non-codel | Methylated | 0 | 250 | 0.00000 |
| CGGA_1618 | Male | 31 | Primary | Astrocytoma | III | Mutant | Non-codel | Unmethylated | 0 | 1850 | 0.00000 |
| CGGA_1619 | Male | 44 | Primary | Astrocytoma | III | Mutant | Non-codel | Methylated | 1 | 766 | 0.00000 |
| CGGA_1620 | Male | 51 | Primary | Oligodendroglioma | III | Mutant | Codel | Methylated | 0 | 1850 | 0.00000 |
| CGGA_1626 | Male | 66 | Primary | Glioblastoma | IV | Wildtype | Non-codel | Unmethylated | 1 | 696 | 8.67767 |
| CGGA_1627 | Male | 37 | Primary | Astrocytoma | III | Mutant | Non-codel | Unmethylated | 0 | 1850 | 0.00000 |
| CGGA_1640 | Male | 55 | Primary | Oligoastrocytoma | III | [Not available] | Non-codel | Unmethylated | 0 | 1850 | 0.00000 |
| CGGA_1644 | Male | 48 | Primary | Glioblastoma | IV | Wildtype | Non-codel | Methylated | 1 | 173 | 0.00000 |
| CGGA_1645 | Male | 59 | Primary | Astrocytoma | III | Mutant | Non-codel | Methylated | 0 | 1850 | 0.22079 |
| CGGA_1650 | Male | 36 | Primary | Astrocytoma | IV | Mutant | Non-codel | Methylated | 1 | 1283 | 0.00000 |
| CGGA_1666 | Male | 60 | Primary | Glioblastoma | IV | Wildtype | [Not available] | Methylated | 1 | 249 | 0.00000 |
| CGGA_1675 | Male | 38 | Primary | Astrocytoma | III | Mutant | Non-codel | Methylated | 0 | 1850 | 0.00000 |
| CGGA_1678 | Male | 51 | Primary | Glioblastoma | IV | Wildtype | [Not available] | Methylated | 1 | 657 | 0.00000 |
| CGGA_1680 | Male | 45 | Primary | Astrocytoma | III | Mutant | Non-codel | Methylated | 1 | 596 | 0.00000 |
| CGGA_1686 | Male | 27 | Primary | Astrocytoma | III | Wildtype | Non-codel | Unmethylated | 1 | 745 | 0.00000 |

| | | | | | | | | | | | |
|---|---|---|---|---|---|---|---|---|---|---|---|
| CGGA_1687 | Male | 14 | Primary | Glioblastoma | IV | Wildtype | Non-codel | Unmethylated | 0 | 1850 | 1.85368 |
| CGGA_1690 | Male | 60 | Primary | Glioblastoma | IV | Wildtype | Non-codel | Unmethylated | 1 | 592 | 0.84352 |
| CGGA_1693 | Male | 61 | Primary | Oligodendroglioma | III | [Not available] | Non-codel | Unmethylated | 1 | 517 | 0.00000 |
| CGGA_1694 | Male | 55 | Primary | Glioblastoma | IV | Wildtype | Non-codel | Unmethylated | 1 | 624 | 6.33481 |
| CGGA_1696 | Male | 72 | Primary | Astrocytoma | III | Wildtype | [Not available] | Unmethylated | 0 | 1205 | 0.01621 |
| CGGA_1700 | Male | 49 | Primary | Oligodendroglioma | III | Mutant | Codel | Unmethylated | 0 | 1850 | 0.00000 |
| CGGA_1701 | Male | 44 | Primary | Oligodendroglioma | III | Mutant | Codel | Methylated | 0 | 1696 | 0.08395 |
| CGGA_1706 | Male | 60 | Primary | Glioblastoma | IV | Wildtype | Non-codel | [Not available] | 0 | 1850 | 0.89974 |
| CGGA_1709 | Male | 47 | Primary | Glioblastoma | IV | Wildtype | Non-codel | Unmethylated | 1 | 415 | 14.54265 |
| CGGA_1713 | Male | 62 | Primary | Glioblastoma | IV | Wildtype | Non-codel | Methylated | 1 | 332 | 0.00000 |
| CGGA_1727 | Male | 48 | Primary | Astrocytoma | IV | Mutant | Non-codel | Unmethylated | 0 | 1850 | 0.00000 |
| CGGA_1728 | Male | 45 | Primary | Astrocytoma | IV | Mutant | Non-codel | Unmethylated | 1 | 917 | 0.00000 |
| CGGA_1735 | Male | 54 | Primary | Glioblastoma | IV | Wildtype | Non-codel | Unmethylated | 1 | 813 | 0.63366 |
| CGGA_1737 | Male | 33 | Primary | Oligodendroglioma | III | Mutant | Codel | Methylated | 0 | 1850 | 0.00000 |
| CGGA_1738 | Male | 49 | Primary | Astrocytoma | III | Wildtype | Non-codel | Methylated | 1 | 469 | 0.13865 |
| CGGA_1744 | Male | 51 | Primary | [Not available] | IV | [Not available] | Non-codel | Methylated | 0 | 1850 | 0.45966 |
| CGGA_1764 | Male | 33 | Primary | Astrocytoma | IV | Mutant | Non-codel | Unmethylated | 1 | 710 | 0.00000 |
| CGGA_1767 | Male | 63 | Primary | Glioblastoma | IV | Wildtype | [Not available] | Methylated | 1 | 86 | 0.00000 |
| CGGA_1791 | Male | 55 | Primary | Astrocytoma | III | Wildtype | [Not available] | Unmethylated | 1 | 855 | 0.00000 |
| CGGA_1809 | Male | 27 | Primary | Oligodendroglioma | III | Wildtype | Non-codel | Unmethylated | 1 | 343 | 0.37226 |
| CGGA_1812 | Male | 65 | Primary | Glioblastoma | IV | Wildtype | Non-codel | Unmethylated | 1 | 780 | 0.00000 |
| CGGA_1819 | Male | 55 | Primary | Glioblastoma | IV | Wildtype | Non-codel | Methylated | 1 | 1005 | 0.46157 |
| CGGA_1854 | Male | 36 | Primary | Astrocytoma | III | Mutant | Non-codel | Methylated | 0 | 1626 | 0.00000 |
| CGGA_1862 | Male | 40 | Primary | Astrocytoma | III | Wildtype | Non-codel | Methylated | 1 | 110 | 2.00068 |
| CGGA_1866 | Male | 68 | Primary | Glioblastoma | IV | Wildtype | Non-codel | Unmethylated | 1 | 127 | 2.39965 |
| CGGA_1870 | Male | 62 | Primary | Astrocytoma | IV | Mutant | Codel | Unmethylated | 0 | 1556 | 0.07142 |
| CGGA_1882 | Male | 33 | Primary | Oligodendroglioma | III | Mutant | Codel | Methylated | 0 | 1531 | 0.10254 |
| CGGA_1901 | Male | 60 | Primary | Glioblastoma | IV | Wildtype | Non-codel | Methylated | 1 | 540 | 0.93171 |
| CGGA_247 | Male | 60 | Primary | Astrocytoma | III | Mutant | Non-codel | Methylated | 1 | 609 | 1.94978 |
| CGGA_277 | Male | 40 | Primary | Oligodendroglioma | III | Mutant | [Not available] | Methylated | 0 | 1850 | 1.38805 |
| CGGA_330 | Male | 45 | Primary | Astrocytoma | III | Wildtype | Non-codel | Unmethylated | 1 | 1121 | 3.20608 |
| CGGA_413 | Male | 54 | Primary | Glioblastoma | IV | Wildtype | Non-codel | Unmethylated | 0 | 183 | 0.00000 |
| CGGA_426 | Male | 50 | Primary | Astrocytoma | III | Wildtype | Non-codel | Unmethylated | 1 | 1560 | 0.00000 |
| CGGA_448 | Male | 64 | Primary | Astrocytoma | III | Wildtype | Non-codel | Unmethylated | 1 | 354 | 0.00000 |
| CGGA_474 | Male | [Not available] | Primary | Oligodendroglioma | III | Wildtype | Non-codel | Unmethylated | 0 | 1850 | 0.13626 |
| CGGA_488 | Male | 23 | Primary | Astrocytoma | III | Wildtype | Non-codel | Unmethylated | 1 | 435 | 1.60149 |

| | | | | | | | | | | | |
|---|---|---|---|---|---|---|---|---|---|---|---|
| CGGA_499 | Male | 51 | Primary | Glioblastoma | IV | Wildtype | Non-codel | Methylated | 1 | 122 | 3.22013 |
| CGGA_509 | Male | 38 | Primary | Glioblastoma | IV | Wildtype | Non-codel | Unmethylated | 1 | 623 | 3.35194 |
| CGGA_510 | Male | 42 | Primary | Oligodendroglioma | III | Mutant | Codel | Methylated | 0 | 1850 | 0.00000 |
| CGGA_521 | Male | 32 | Primary | Astrocytoma | III | Wildtype | Non-codel | Unmethylated | 1 | 514 | 3.89755 |
| CGGA_525 | Male | 61 | Primary | Glioblastoma | IV | Wildtype | Non-codel | Methylated | 1 | 138 | 0.87426 |
| CGGA_560 | Male | 43 | Primary | Astrocytoma | III | Mutant | Non-codel | Methylated | 0 | 1850 | 1.08696 |
| CGGA_564 | Male | 47 | Primary | Astrocytoma | III | Wildtype | Non-codel | Methylated | 1 | 679 | 5.25790 |
| CGGA_565 | Male | 51 | Primary | Astrocytoma | III | Wildtype | Non-codel | Unmethylated | 0 | 1850 | 0.00000 |
| CGGA_604 | Male | 46 | Primary | Glioblastoma | IV | Wildtype | Non-codel | Methylated | 1 | 381 | 3.21157 |
| CGGA_616 | Male | 43 | Primary | Astrocytoma | III | Wildtype | Non-codel | Methylated | 1 | 952 | 0.90296 |
| CGGA_643 | Male | 56 | Primary | Astrocytoma | III | Wildtype | Non-codel | Unmethylated | 1 | 331 | 2.19196 |
| CGGA_658 | Male | 55 | Primary | Glioblastoma | IV | Wildtype | Non-codel | Methylated | 1 | 372 | 0.55976 |
| CGGA_676 | Male | 60 | Primary | Glioblastoma | IV | Wildtype | Non-codel | Unmethylated | 1 | 376 | 2.97993 |
| CGGA_680 | Male | 44 | Primary | Glioblastoma | IV | Wildtype | Non-codel | Unmethylated | 0 | 1850 | 0.31869 |
| CGGA_705 | Male | 41 | Primary | Astrocytoma | III | Mutant | Non-codel | Unmethylated | 1 | 1507 | 0.26589 |
| CGGA_710 | Male | 41 | Primary | Astrocytoma | IV | Mutant | Non-codel | Methylated | 1 | 660 | 0.20179 |
| CGGA_727 | Male | 45 | Primary | Astrocytoma | III | Wildtype | Non-codel | Methylated | 1 | 1023 | 2.61519 |
| CGGA_731 | Male | 50 | Primary | Glioblastoma | IV | Wildtype | Non-codel | Unmethylated | 1 | 503 | 0.00000 |
| CGGA_747 | Male | 29 | Primary | Astrocytoma | IV | Mutant | Non-codel | Methylated | 1 | 970 | 0.00000 |
| CGGA_759 | Male | 30 | Primary | Astrocytoma | IV | Mutant | Non-codel | Methylated | 1 | 1263 | 0.00000 |
| CGGA_761 | Male | 45 | Primary | Astrocytoma | IV | Mutant | Non-codel | Methylated | 1 | 1840 | 0.00000 |
| CGGA_782 | Male | 60 | Primary | Glioblastoma | IV | Wildtype | Non-codel | Unmethylated | 1 | 289 | 3.13119 |
| CGGA_789 | Male | 50 | Primary | Glioblastoma | IV | Wildtype | Non-codel | Unmethylated | 1 | 345 | 1.77893 |
| CGGA_791 | Male | 55 | Primary | Astrocytoma | III | Wildtype | Non-codel | Unmethylated | 1 | 252 | 2.53199 |
| CGGA_802 | Male | 42 | Primary | Glioblastoma | IV | Wildtype | Non-codel | Unmethylated | 1 | 681 | 0.00000 |
| CGGA_808 | Male | 38 | Primary | Glioblastoma | IV | Wildtype | Non-codel | Unmethylated | 0 | 1167 | 0.22893 |
| CGGA_837 | Male | 57 | Primary | Glioblastoma | IV | Wildtype | Non-codel | Methylated | 1 | 432 | 0.00000 |
| CGGA_852 | Male | 40 | Primary | Astrocytoma | III | Wildtype | Non-codel | Unmethylated | 1 | 169 | 6.48398 |
| CGGA_859 | Male | 56 | Primary | Glioblastoma | IV | Wildtype | Non-codel | Unmethylated | 1 | 387 | 0.22631 |
| CGGA_861 | Male | 29 | Primary | Astrocytoma | III | Wildtype | Non-codel | [Not available] | 1 | 893 | 0.12197 |
| CGGA_878 | Male | 42 | Primary | Glioblastoma | IV | Wildtype | Non-codel | Unmethylated | 1 | 863 | 0.21559 |
| CGGA_890 | Male | 39 | Primary | Astrocytoma | III | Mutant | Non-codel | [Not available] | 0 | 1850 | 0.00000 |
| CGGA_901 | Male | 40 | Primary | Astrocytoma | III | Mutant | Non-codel | [Not available] | 1 | 929 | 0.00000 |
| CGGA_902 | Male | 54 | Primary | Glioblastoma | IV | Wildtype | Non-codel | Unmethylated | 1 | 193 | 0.00000 |
| CGGA_D03 | Male | 44 | Primary | Glioblastoma | IV | Wildtype | Non-codel | Methylated | 1 | 423 | 4.58467 |
| CGGA_D09 | Male | 57 | Primary | Glioblastoma | IV | Wildtype | Non-codel | Unmethylated | 1 | 233 | 5.05942 |

| | | | | | | | | | | | |
|---|---|---|---|---|---|---|---|---|---|---|---|
| CGGA_D28 | Male | 25 | Primary | Astrocytoma | III | Mutant | Non-codel | [Not available] | 1 | 860 | 0.00000 |
| CGGA_D30 | Male | 70 | Primary | Glioblastoma | IV | Wildtype | Non-codel | [Not available] | 1 | 215 | 0.44242 |
| CGGA_D37 | Male | 55 | Primary | Glioblastoma | IV | Wildtype | Non-codel | Unmethylated | 1 | 965 | 2.66397 |
| CGGA_D52 | Male | 46 | Primary | Oligodendroglioma | III | Mutant | Codel | Unmethylated | 0 | 1850 | 0.05599 |
| CGGA_D57 | Male | 40 | Primary | Glioblastoma | IV | Wildtype | Non-codel | [Not available] | 1 | 766 | 1.32427 |
| CGGA_J023 | Male | 36 | Primary | Oligodendroglioma | III | Mutant | Non-codel | [Not available] | 1 | 1028 | 0.00000 |
| CGGA_P100 | Male | 67 | Primary | Glioblastoma | IV | Wildtype | Non-codel | [Not available] | 1 | 268 | 3.02606 |
| CGGA_P102 | Male | 30 | Primary | Astrocytoma | IV | Mutant | Non-codel | [Not available] | 1 | 1269 | 0.00000 |
| CGGA_P112 | Male | 65 | Primary | Glioblastoma | IV | Wildtype | Non-codel | [Not available] | 1 | 834 | 0.44424 |
| CGGA_P116 | Male | 58 | Primary | Glioblastoma | IV | Wildtype | [Not available] | [Not available] | 1 | 305 | 0.00000 |
| CGGA_P142 | Male | 45 | Primary | Oligodendroglioma | III | Mutant | Codel | [Not available] | 0 | 1558 | 0.00000 |
| CGGA_P144 | Male | 44 | Primary | Astrocytoma | III | Mutant | Non-codel | [Not available] | 0 | 1559 | 0.00000 |
| CGGA_P15 | Male | 49 | Primary | [Not available] | IV | [Not available] | Non-codel | [Not available] | 1 | 723 | 0.00000 |
| CGGA_P159 | Male | 59 | Primary | Astrocytoma | III | Wildtype | Non-codel | [Not available] | 1 | 211 | 0.15376 |
| CGGA_P164 | Male | 27 | Primary | Glioblastoma | IV | Wildtype | [Not available] | [Not available] | 0 | 1553 | 0.00000 |
| CGGA_P17 | Male | 30 | Primary | Oligodendroglioma | III | Mutant | Codel | [Not available] | 0 | 1850 | 0.00000 |
| CGGA_P172 | Male | 33 | Primary | Astrocytoma | III | Mutant | [Not available] | [Not available] | 1 | 1049 | 0.00000 |
| CGGA_P18 | Male | 40 | Primary | Astrocytoma | III | Mutant | Non-codel | [Not available] | 1 | 387 | 0.25799 |
| CGGA_P180 | Male | 47 | Primary | Astrocytoma | IV | Mutant | [Not available] | [Not available] | 1 | 260 | 0.00000 |
| CGGA_P183 | Male | 58 | Primary | Oligodendroglioma | III | Mutant | Codel | [Not available] | 0 | 1512 | 0.11426 |
| CGGA_P205 | Male | 66 | Primary | Glioblastoma | IV | Wildtype | [Not available] | [Not available] | 1 | 583 | 0.01302 |
| CGGA_P22 | Male | 62 | Primary | Glioblastoma | IV | Wildtype | Non-codel | [Not available] | 1 | 406 | 4.38507 |
| CGGA_P25 | Male | 64 | Primary | Glioblastoma | IV | Wildtype | [Not available] | [Not available] | 1 | 147 | 0.00000 |
| CGGA_P265 | Male | 36 | Primary | Astrocytoma | III | Wildtype | [Not available] | Methylated | 1 | 779 | 0.00000 |
| CGGA_P28 | Male | 61 | Primary | Glioblastoma | IV | Wildtype | Non-codel | [Not available] | 1 | 107 | 0.00000 |

**Table S4. List of SRY human target genes identified in chromatin immunoprecipitation experiments.**

| gene.id | gene.symbol |
|---|---|
| ENSG00000129673 | AANAT |
| ENSG00000181409 | AATK |
| ENSG00000167972 | ABCA3 |
| ENSG00000064687 | ABCA7 |
| ENSG00000073734 | ABCB11 |
| ENSG00000150967 | ABCB9 |
| ENSG00000033050 | ABCF2 |
| ENSG00000143921 | ABCG8 |
| ENSG00000114779 | ABHD14B |
| ENSG00000100439 | ABHD4 |
| ENSG00000138443 | ABI2 |
| ENSG00000097007 | ABL1 |
| ENSG00000114626 | ABTB1 |
| ENSG00000167315 | ACAA2 |
| ENSG00000072818 | ACAP1 |
| ENSG00000120437 | ACAT2 |
| ENSG00000159640 | ACE |
| ENSG00000177076 | ACER2 |
| ENSG00000087085 | ACHE |
| ENSG00000131473 | ACLY |
| ENSG00000172497 | ACOT12 |
| ENSG00000168306 | ACOX2 |
| ENSG00000103740 | ACSBG1 |
| ENSG00000130377 | ACSBG2 |
| ENSG00000167107 | ACSF2 |
| ENSG00000176715 | ACSF3 |
| ENSG00000164398 | ACSL6 |
| ENSG00000111058 | ACSS3 |
| ENSG00000143632 | ACTA1 |

| | |
|---|---|
| ENSG00000075624 | ACTB |
| ENSG00000169067 | ACTBL2 |
| ENSG00000136518 | ACTL6A |
| ENSG00000072110 | ACTN1 |
| ENSG00000115091 | ACTR3 |
| ENSG00000139567 | ACVRL1 |
| ENSG00000132744 | ACY3 |
| ENSG00000196839 | ADA |
| ENSG00000137845 | ADAM10 |
| ENSG00000073670 | ADAM11 |
| ENSG00000104755 | ADAM2 |
| ENSG00000142303 | ADAMTS10 |
| ENSG00000151388 | ADAMTS12 |
| ENSG00000166106 | ADAMTS15 |
| ENSG00000145536 | ADAMTS16 |
| ENSG00000140470 | ADAMTS17 |
| ENSG00000145808 | ADAMTS19 |
| ENSG00000087116 | ADAMTS2 |
| ENSG00000173157 | ADAMTS20 |
| ENSG00000154736 | ADAMTS5 |
| ENSG00000136378 | ADAMTS7 |
| ENSG00000197859 | ADAMTSL2 |
| ENSG00000184060 | ADAP2 |
| ENSG00000197381 | ADARB1 |
| ENSG00000129467 | ADCY4 |
| ENSG00000173175 | ADCY5 |
| ENSG00000121281 | ADCY7 |
| ENSG00000078549 | ADCYAP1R1 |
| ENSG00000197894 | ADH5 |
| ENSG00000147576 | ADHFE1 |
| ENSG00000182551 | ADI1 |

| | |
|---|---|
| ENSG00000282608 | ADORA3 |
| ENSG00000153531 | ADPRHL1 |
| ENSG00000150594 | ADRA2A |
| ENSG00000184160 | ADRA2C |
| ENSG00000188778 | ADRB3 |
| ENSG00000072364 | AFF4 |
| ENSG00000135439 | AGAP2 |
| ENSG00000133612 | AGAP3 |
| ENSG00000084693 | AGBL5 |
| ENSG00000106351 | AGFG2 |
| ENSG00000204310 | AGPAT1 |
| ENSG00000188157 | AGRN |
| ENSG00000135744 | AGT |
| ENSG00000144891 | AGTR1 |
| ENSG00000172482 | AGXT |
| ENSG00000126705 | AHDC1 |
| ENSG00000126878 | AIF1L |
| ENSG00000042286 | AIFM2 |
| ENSG00000183773 | AIFM3 |
| ENSG00000129221 | AIPL1 |
| ENSG00000160224 | AIRE |
| ENSG00000147853 | AK3 |
| ENSG00000162433 | AK4 |
| ENSG00000121057 | AKAP1 |
| ENSG00000111254 | AKAP3 |
| ENSG00000204673 | AKT1S1 |
| ENSG00000166971 | AKTIP |
| ENSG00000148218 | ALAD |
| ENSG00000006534 | ALDH3B1 |
| ENSG00000109107 | ALDOC |
| ENSG00000033011 | ALG1 |

| | |
|---|---|
| ENSG00000086848 | ALG9 |
| ENSG00000171094 | ALK |
| ENSG00000161905 | ALOX15 |
| ENSG00000106927 | AMBP |
| ENSG00000110497 | AMBRA1 |
| ENSG00000162066 | AMDHD2 |
| ENSG00000159461 | AMFR |
| ENSG00000176020 | AMIGO3 |
| ENSG00000166126 | AMN |
| ENSG00000114019 | AMOTL2 |
| ENSG00000029534 | ANK1 |
| ENSG00000160117 | ANKLE1 |
| ENSG00000101745 | ANKRD12 |
| ENSG00000206560 | ANKRD28 |
| ENSG00000135299 | ANKRD6 |
| ENSG00000171714 | ANO5 |
| ENSG00000163297 | ANTXR2 |
| ENSG00000196975 | ANXA4 |
| ENSG00000164111 | ANXA5 |
| ENSG00000106367 | AP1S1 |
| ENSG00000182287 | AP1S2 |
| ENSG00000103723 | AP3B2 |
| ENSG00000077420 | APBB1IP |
| ENSG00000115266 | APC2 |
| ENSG00000166181 | API5 |
| ENSG00000169621 | APLF |
| ENSG00000179750 | APOBEC3B |
| ENSG00000234906 | APOC2 |
| ENSG00000267467 | APOC4 |
| ENSG00000130203 | APOE |
| ENSG00000062725 | APPBP2 |

| | |
|---|---|
| ENSG00000178301 | AQP11 |
| ENSG00000184945 | AQP12A |
| ENSG00000185176 | AQP12B |
| ENSG00000165269 | AQP7 |
| ENSG00000103375 | AQP8 |
| ENSG00000169083 | AR |
| ENSG00000186635 | ARAP1 |
| ENSG00000198576 | ARC |
| ENSG00000095139 | ARCN1 |
| ENSG00000143761 | ARF1 |
| ENSG00000134287 | ARF3 |
| ENSG00000242247 | ARFGAP3 |
| ENSG00000081181 | ARG2 |
| ENSG00000175220 | ARHGAP1 |
| ENSG00000163219 | ARHGAP25 |
| ENSG00000160007 | ARHGAP35 |
| ENSG00000047648 | ARHGAP6 |
| ENSG00000141522 | ARHGDIA |
| ENSG00000074964 | ARHGEF10L |
| ENSG00000142632 | ARHGEF19 |
| ENSG00000240771 | ARHGEF25 |
| ENSG00000165801 | ARHGEF40 |
| ENSG00000179361 | ARID3B |
| ENSG00000150347 | ARID5B |
| ENSG00000166233 | ARIH1 |
| ENSG00000172379 | ARNT2 |
| ENSG00000130429 | ARPC1B |
| ENSG00000197070 | ARRDC1 |
| ENSG00000100299 | ARSA |
| ENSG00000129744 | ART1 |
| ENSG00000111339 | ART4 |

| | |
|---|---|
| ENSG00000167311 | ART5 |
| ENSG00000117407 | ARTN |
| ENSG00000161664 | ASB16 |
| ENSG00000148331 | ASB6 |
| ENSG00000177981 | ASB8 |
| ENSG00000138303 | ASCC1 |
| ENSG00000183734 | ASCL2 |
| ENSG00000204653 | ASPDH |
| ENSG00000106819 | ASPN |
| ENSG00000152092 | ASTN1 |
| ENSG00000148219 | ASTN2 |
| ENSG00000171456 | ASXL1 |
| ENSG00000143970 | ASXL2 |
| ENSG00000137343 | ATAT1 |
| ENSG00000167654 | ATCAY |
| ENSG00000126775 | ATG14 |
| ENSG00000110046 | ATG2A |
| ENSG00000181652 | ATG9B |
| ENSG00000149311 | ATM |
| ENSG00000166454 | ATMIN |
| ENSG00000111676 | ATN1 |
| ENSG00000101974 | ATP11C |
| ENSG00000143153 | ATP1B1 |
| ENSG00000196296 | ATP2A1 |
| ENSG00000074370 | ATP2A3 |
| ENSG00000058668 | ATP2B4 |
| ENSG00000105675 | ATP4A |
| ENSG00000185344 | ATP6V0A2 |
| ENSG00000117410 | ATP6V0B |
| ENSG00000159720 | ATP6V0D1 |
| ENSG00000250565 | ATP6V1E2 |

| | |
|---|---|
| ENSG00000128524 | ATP6V1F |
| ENSG00000124406 | ATP8A1 |
| ENSG00000130270 | ATP8B3 |
| ENSG00000171953 | ATPAF2 |
| ENSG00000124788 | ATXN1 |
| ENSG00000135407 | AVIL |
| ENSG00000101200 | AVP |
| ENSG00000126895 | AVPR2 |
| ENSG00000163512 | AZI2 |
| ENSG00000162885 | B3GALNT2 |
| ENSG00000235863 | B3GALT4 |
| ENSG00000149541 | B3GAT3 |
| ENSG00000156966 | B3GNT7 |
| ENSG00000177191 | B3GNT8 |
| ENSG00000117411 | B4GALT2 |
| ENSG00000186318 | BACE1 |
| ENSG00000156273 | BACH1 |
| ENSG00000112182 | BACH2 |
| ENSG00000175866 | BAIAP2 |
| ENSG00000006453 | BAIAP2L1 |
| ENSG00000095739 | BAMBI |
| ENSG00000043039 | BARX2 |
| ENSG00000123685 | BATF3 |
| ENSG00000087088 | BAX |
| ENSG00000123636 | BAZ2B |
| ENSG00000174483 | BBS1 |
| ENSG00000187244 | BCAM |
| ENSG00000132692 | BCAN |
| ENSG00000185825 | BCAP31 |
| ENSG00000141376 | BCAS3 |
| ENSG00000060982 | BCAT1 |

| | |
|---|---|
| ENSG00000119866 | BCL11A |
| ENSG00000127152 | BCL11B |
| ENSG00000099968 | BCL2L13 |
| ENSG00000110987 | BCL7A |
| ENSG00000085185 | BCORL1 |
| ENSG00000100739 | BDKRB1 |
| ENSG00000127325 | BEST3 |
| ENSG00000242252 | BGLAP |
| ENSG00000182492 | BGN |
| ENSG00000122870 | BICC1 |
| ENSG00000136717 | BIN1 |
| ENSG00000125845 | BMP2 |
| ENSG00000101144 | BMP7 |
| ENSG00000113734 | BNIP1 |
| ENSG00000176171 | BNIP3 |
| ENSG00000104765 | BNIP3L |
| ENSG00000145919 | BOD1 |
| ENSG00000163170 | BOLA3 |
| ENSG00000186191 | BPIFB4 |
| ENSG00000089234 | BRAP |
| ENSG00000204256 | BRD2 |
| ENSG00000184992 | BRI3BP |
| ENSG00000174744 | BRMS1 |
| ENSG00000174672 | BRSK2 |
| ENSG00000172270 | BSG |
| ENSG00000132640 | BTBD3 |
| ENSG00000159388 | BTG2 |
| ENSG00000154640 | BTG3 |
| ENSG00000124508 | BTN2A2 |
| ENSG00000137656 | BUD13 |
| ENSG00000165863 | C10orf82 |

| | |
|---|---|
| ENSG00000171067 | C11orf24 |
| ENSG00000188070 | C11orf95 |
| ENSG00000130921 | C12orf65 |
| ENSG00000184601 | C14orf180 |
| ENSG00000100802 | C14orf93 |
| ENSG00000188277 | C15orf62 |
| ENSG00000155330 | C16orf87 |
| ENSG00000152242 | C18orf25 |
| ENSG00000188032 | C19orf67 |
| ENSG00000183397 | C19orf71 |
| ENSG00000000460 | C1orf112 |
| ENSG00000143793 | C1orf35 |
| ENSG00000144119 | C1QL2 |
| ENSG00000186897 | C1QL4 |
| ENSG00000173918 | C1QTNF1 |
| ENSG00000205863 | C1QTNF9B |
| ENSG00000101220 | C20orf27 |
| ENSG00000169314 | C22orf15 |
| ENSG00000172375 | C2CD2L |
| ENSG00000187068 | C3orf70 |
| ENSG00000197405 | C5AR1 |
| ENSG00000205765 | C5orf51 |
| ENSG00000112539 | C6orf118 |
| ENSG00000182307 | C8orf33 |
| ENSG00000156172 | C8orf37 |
| ENSG00000171159 | C9orf16 |
| ENSG00000135045 | C9orf40 |
| ENSG00000157653 | C9orf43 |
| ENSG00000063180 | CA11 |
| ENSG00000168748 | CA7 |
| ENSG00000107159 | CA9 |

| | |
|---|---|
| ENSG00000148408 | CACNA1B |
| ENSG00000151067 | CACNA1C |
| ENSG00000006283 | CACNA1G |
| ENSG00000151062 | CACNA2D4 |
| ENSG00000067191 | CACNB1 |
| ENSG00000167535 | CACNB3 |
| ENSG00000108878 | CACNG1 |
| ENSG00000075461 | CACNG4 |
| ENSG00000084774 | CAD |
| ENSG00000175161 | CADM2 |
| ENSG00000105767 | CADM4 |
| ENSG00000081803 | CADPS2 |
| ENSG00000183166 | CALN1 |
| ENSG00000008118 | CAMK1G |
| ENSG00000070808 | CAMK2A |
| ENSG00000145349 | CAMK2D |
| ENSG00000148660 | CAMK2G |
| ENSG00000042493 | CAPG |
| ENSG00000014216 | CAPN1 |
| ENSG00000142330 | CAPN10 |
| ENSG00000182472 | CAPN12 |
| ENSG00000149260 | CAPN5 |
| ENSG00000131375 | CAPN7 |
| ENSG00000135773 | CAPN9 |
| ENSG00000198898 | CAPZA2 |
| ENSG00000100065 | CARD10 |
| ENSG00000141527 | CARD14 |
| ENSG00000164326 | CARTPT |
| ENSG00000143318 | CASQ1 |
| ENSG00000099338 | CATSPERG |
| ENSG00000105971 | CAV2 |

| | |
|---|---|
| ENSG00000141668 | CBLN2 |
| ENSG00000141570 | CBX8 |
| ENSG00000135736 | CCDC102A |
| ENSG00000173581 | CCDC106 |
| ENSG00000105479 | CCDC114 |
| ENSG00000104957 | CCDC130 |
| ENSG00000185298 | CCDC137 |
| ENSG00000163492 | CCDC141 |
| ENSG00000160050 | CCDC28B |
| ENSG00000165972 | CCDC38 |
| ENSG00000160124 | CCDC58 |
| ENSG00000115355 | CCDC88A |
| ENSG00000105321 | CCDC9 |
| ENSG00000125633 | CCDC93 |
| ENSG00000173013 | CCDC96 |
| ENSG00000102970 | CCL17 |
| ENSG00000118971 | CCND2 |
| ENSG00000129315 | CCNT1 |
| ENSG00000112486 | CCR6 |
| ENSG00000173992 | CCS |
| ENSG00000163468 | CCT3 |
| ENSG00000135624 | CCT7 |
| ENSG00000156535 | CD109 |
| ENSG00000177697 | CD151 |
| ENSG00000012124 | CD22 |
| ENSG00000174807 | CD248 |
| ENSG00000139193 | CD27 |
| ENSG00000103855 | CD276 |
| ENSG00000169217 | CD2BP2 |
| ENSG00000204345 | CD300LD |
| ENSG00000167775 | CD320 |

| | |
|---|---|
| ENSG00000196776 | CD47 |
| ENSG00000110448 | CD5 |
| ENSG00000013725 | CD6 |
| ENSG00000135404 | CD63 |
| ENSG00000112149 | CD83 |
| ENSG00000081377 | CDC14B |
| ENSG00000106993 | CDC37L1 |
| ENSG00000198752 | CDC42BPB |
| ENSG00000149798 | CDC42EP2 |
| ENSG00000163171 | CDC42EP3 |
| ENSG00000179604 | CDC42EP4 |
| ENSG00000134371 | CDC73 |
| ENSG00000164649 | CDCA7L |
| ENSG00000140937 | CDH11 |
| ENSG00000129910 | CDH15 |
| ENSG00000149654 | CDH22 |
| ENSG00000107736 | CDH23 |
| ENSG00000062038 | CDH3 |
| ENSG00000167258 | CDK12 |
| ENSG00000058091 | CDK14 |
| ENSG00000117266 | CDK18 |
| ENSG00000176749 | CDK5R1 |
| ENSG00000108465 | CDK5RAP3 |
| ENSG00000136807 | CDK9 |
| ENSG00000237190 | CDKN2AIPNL |
| ENSG00000167513 | CDT1 |
| ENSG00000091527 | CDV3 |
| ENSG00000213892 | CEACAM16 |
| ENSG00000186567 | CEACAM19 |
| ENSG00000245848 | CEBPA |
| ENSG00000092067 | CEBPE |

| | |
|---|---|
| ENSG00000170835 | CEL |
| ENSG00000139610 | CELA1 |
| ENSG00000142615 | CELA2A |
| ENSG00000219073 | CELA3B |
| ENSG00000159409 | CELF3 |
| ENSG00000143126 | CELSR2 |
| ENSG00000008300 | CELSR3 |
| ENSG00000184524 | CEND1 |
| ENSG00000168944 | CEP120 |
| ENSG00000138180 | CEP55 |
| ENSG00000183137 | CEP57L1 |
| ENSG00000114107 | CEP70 |
| ENSG00000167123 | CERCAM |
| ENSG00000100422 | CERK |
| ENSG00000159398 | CES5A |
| ENSG00000197766 | CFD |
| ENSG00000205403 | CFI |
| ENSG00000001626 | CFTR |
| ENSG00000100532 | CGRRF1 |
| ENSG00000159259 | CHAF1B |
| ENSG00000106153 | CHCHD2 |
| ENSG00000124177 | CHD6 |
| ENSG00000177200 | CHD9 |
| ENSG00000100604 | CHGA |
| ENSG00000109220 | CHIC2 |
| ENSG00000177830 | CHID1 |
| ENSG00000100288 | CHKB |
| ENSG00000130724 | CHMP2A |
| ENSG00000090539 | CHRD |
| ENSG00000180720 | CHRM4 |
| ENSG00000080644 | CHRNA3 |

| | |
|---|---|
| ENSG00000170175 | CHRNB1 |
| ENSG00000135902 | CHRND |
| ENSG00000196811 | CHRNG |
| ENSG00000115526 | CHST10 |
| ENSG00000213341 | CHUK |
| ENSG00000079432 | CIC |
| ENSG00000125931 | CITED1 |
| ENSG00000104879 | CKM |
| ENSG00000223572 | CKMT1A |
| ENSG00000237289 | CKMT1B |
| ENSG00000175505 | CLCF1 |
| ENSG00000186510 | CLCNKA |
| ENSG00000134873 | CLDN10 |
| ENSG00000013297 | CLDN11 |
| ENSG00000157224 | CLDN12 |
| ENSG00000066405 | CLDN18 |
| ENSG00000164007 | CLDN19 |
| ENSG00000165215 | CLDN3 |
| ENSG00000189143 | CLDN4 |
| ENSG00000181885 | CLDN7 |
| ENSG00000213937 | CLDN9 |
| ENSG00000080822 | CLDND1 |
| ENSG00000105472 | CLEC11A |
| ENSG00000140839 | CLEC18B |
| ENSG00000152672 | CLEC4F |
| ENSG00000169583 | CLIC3 |
| ENSG00000130779 | CLIP1 |
| ENSG00000165959 | CLMN |
| ENSG00000188603 | CLN3 |
| ENSG00000134852 | CLOCK |
| ENSG00000137392 | CLPS |

| | |
|---|---|
| ENSG00000049656 | CLPTM1L |
| ENSG00000171603 | CLSTN1 |
| ENSG00000120885 | CLU |
| ENSG00000174600 | CMKLR1 |
| ENSG00000134326 | CMPK2 |
| ENSG00000140931 | CMTM3 |
| ENSG00000183723 | CMTM4 |
| ENSG00000166091 | CMTM5 |
| ENSG00000144191 | CNGA3 |
| ENSG00000132259 | CNGA4 |
| ENSG00000174871 | CNIH2 |
| ENSG00000143771 | CNIH4 |
| ENSG00000064666 | CNN2 |
| ENSG00000257727 | CNPY2 |
| ENSG00000137161 | CNPY3 |
| ENSG00000166997 | CNPY4 |
| ENSG00000122756 | CNTFR |
| ENSG00000068120 | COASY |
| ENSG00000100473 | COCH |
| ENSG00000121058 | COIL |
| ENSG00000204248 | COL11A2 |
| ENSG00000108821 | COL1A1 |
| ENSG00000050767 | COL23A1 |
| ENSG00000196739 | COL27A1 |
| ENSG00000139219 | COL2A1 |
| ENSG00000134871 | COL4A2 |
| ENSG00000081052 | COL4A4 |
| ENSG00000130635 | COL5A1 |
| ENSG00000142173 | COL6A2 |
| ENSG00000110442 | COMMD9 |
| ENSG00000105664 | COMP |

| | |
|---|---|
| ENSG00000129083 | COPB1 |
| ENSG00000168090 | COPS6 |
| ENSG00000111652 | COPS7A |
| ENSG00000144524 | COPS7B |
| ENSG00000173085 | COQ2 |
| ENSG00000088682 | COQ9 |
| ENSG00000102879 | CORO1A |
| ENSG00000172725 | CORO1B |
| ENSG00000240230 | COX19 |
| ENSG00000131055 | COX4I2 |
| ENSG00000112695 | COX7A2 |
| ENSG00000168993 | CPLX1 |
| ENSG00000085719 | CPNE3 |
| ENSG00000178773 | CPNE7 |
| ENSG00000139117 | CPNE8 |
| ENSG00000144550 | CPNE9 |
| ENSG00000165934 | CPSF2 |
| ENSG00000187959 | CPSF4L |
| ENSG00000205560 | CPT1B |
| ENSG00000088882 | CPXM1 |
| ENSG00000109625 | CPZ |
| ENSG00000143162 | CREG1 |
| ENSG00000175874 | CREG2 |
| ENSG00000163703 | CRELD1 |
| ENSG00000184164 | CRELD2 |
| ENSG00000213145 | CRIP1 |
| ENSG00000167193 | CRK |
| ENSG00000099942 | CRKL |
| ENSG00000006016 | CRLF1 |
| ENSG00000058453 | CROCC |
| ENSG00000109943 | CRTAM |

| | |
|---|---|
| ENSG00000140577 | CRTC3 |
| ENSG00000008405 | CRY1 |
| ENSG00000160202 | CRYAA |
| ENSG00000100122 | CRYBB1 |
| ENSG00000244752 | CRYBB2 |
| ENSG00000100053 | CRYBB3 |
| ENSG00000118231 | CRYGD |
| ENSG00000205758 | CRYZL1 |
| ENSG00000172346 | CSDC2 |
| ENSG00000164400 | CSF2 |
| ENSG00000108342 | CSF3 |
| ENSG00000204414 | CSHL1 |
| ENSG00000103653 | CSK |
| ENSG00000070770 | CSNK2A2 |
| ENSG00000110925 | CSRNP2 |
| ENSG00000077984 | CST7 |
| ENSG00000160213 | CSTB |
| ENSG00000175029 | CTBP2 |
| ENSG00000060069 | CTDP1 |
| ENSG00000144579 | CTDSP1 |
| ENSG00000175215 | CTDSP2 |
| ENSG00000144677 | CTDSPL |
| ENSG00000198730 | CTR9 |
| ENSG00000164733 | CTSB |
| ENSG00000174080 | CTSF |
| ENSG00000143387 | CTSK |
| ENSG00000136943 | CTSV |
| ENSG00000172543 | CTSW |
| ENSG00000085733 | CTTN |
| ENSG00000178531 | CTXN1 |
| ENSG00000107874 | CUEDC2 |

| | |
|---|---|
| ENSG00000112514 | CUTA |
| ENSG00000111249 | CUX2 |
| ENSG00000109182 | CWH43 |
| ENSG00000163464 | CXCR1 |
| ENSG00000204165 | CXorf65 |
| ENSG00000168772 | CXXC4 |
| ENSG00000008283 | CYB561 |
| ENSG00000167740 | CYB5D2 |
| ENSG00000051523 | CYBA |
| ENSG00000172115 | CYCS |
| ENSG00000055163 | CYFIP2 |
| ENSG00000179142 | CYP11B2 |
| ENSG00000138061 | CYP1B1 |
| ENSG00000095596 | CYP26A1 |
| ENSG00000003137 | CYP26B1 |
| ENSG00000135929 | CYP27A1 |
| ENSG00000134716 | CYP2J2 |
| ENSG00000167600 | CYP2S1 |
| ENSG00000186526 | CYP4F8 |
| ENSG00000108669 | CYTH1 |
| ENSG00000164535 | DAGLB |
| ENSG00000112977 | DAP |
| ENSG00000205944 | DAZ2 |
| ENSG00000183283 | DAZAP2 |
| ENSG00000123454 | DBH |
| ENSG00000113758 | DBN1 |
| ENSG00000105516 | DBP |
| ENSG00000137992 | DBT |
| ENSG00000109851 | DBX1 |
| ENSG00000175203 | DCTN2 |
| ENSG00000166847 | DCTN5 |

| | |
|---|---|
| ENSG00000169738 | DCXR |
| ENSG00000134574 | DDB2 |
| ENSG00000162733 | DDR2 |
| ENSG00000100201 | DDX17 |
| ENSG00000198563 | DDX39B |
| ENSG00000123064 | DDX54 |
| ENSG00000110367 | DDX6 |
| ENSG00000242612 | DECR2 |
| ENSG00000023892 | DEF6 |
| ENSG00000162777 | DENND2D |
| ENSG00000182108 | DEXI |
| ENSG00000070413 | DGCR2 |
| ENSG00000145214 | DGKQ |
| ENSG00000149091 | DGKZ |
| ENSG00000116133 | DHCR24 |
| ENSG00000139549 | DHH |
| ENSG00000100612 | DHRS7 |
| ENSG00000132153 | DHX30 |
| ENSG00000108771 | DHX58 |
| ENSG00000135829 | DHX9 |
| ENSG00000100697 | DICER1 |
| ENSG00000211452 | DIO1 |
| ENSG00000197406 | DIO3 |
| ENSG00000176490 | DIRAS1 |
| ENSG00000162946 | DISC1 |
| ENSG00000170579 | DLGAP1 |
| ENSG00000116544 | DLGAP3 |
| ENSG00000080845 | DLGAP4 |
| ENSG00000128917 | DLL4 |
| ENSG00000197587 | DMBX1 |
| ENSG00000161249 | DMKN |

| | |
|---|---|
| ENSG00000104936 | DMPK |
| ENSG00000135164 | DMTF1 |
| ENSG00000185800 | DMWD |
| ENSG00000086061 | DNAJA1 |
| ENSG00000090520 | DNAJB11 |
| ENSG00000148719 | DNAJB12 |
| ENSG00000179407 | DNAJB8 |
| ENSG00000128590 | DNAJB9 |
| ENSG00000136770 | DNAJC1 |
| ENSG00000135392 | DNAJC14 |
| ENSG00000178401 | DNAJC22 |
| ENSG00000059769 | DNAJC25 |
| ENSG00000116675 | DNAJC6 |
| ENSG00000167968 | DNASE1L2 |
| ENSG00000187957 | DNER |
| ENSG00000197959 | DNM3 |
| ENSG00000142182 | DNMT3L |
| ENSG00000123992 | DNPEP |
| ENSG00000088538 | DOCK3 |
| ENSG00000129932 | DOHH |
| ENSG00000146094 | DOK3 |
| ENSG00000104885 | DOT1L |
| ENSG00000172269 | DPAGT1 |
| ENSG00000015413 | DPEP1 |
| ENSG00000108963 | DPH1 |
| ENSG00000132768 | DPH2 |
| ENSG00000179085 | DPM3 |
| ENSG00000175497 | DPP10 |
| ENSG00000130226 | DPP6 |
| ENSG00000203909 | DPPA5 |
| ENSG00000188641 | DPYD |

| | |
|---|---|
| ENSG00000147647 | DPYS |
| ENSG00000113657 | DPYSL3 |
| ENSG00000175550 | DRAP1 |
| ENSG00000184845 | DRD1 |
| ENSG00000151577 | DRD3 |
| ENSG00000102385 | DRP2 |
| ENSG00000171587 | DSCAM |
| ENSG00000047579 | DTNBP1 |
| ENSG00000178498 | DTX3 |
| ENSG00000110042 | DTX4 |
| ENSG00000140254 | DUOXA1 |
| ENSG00000141994 | DUS3L |
| ENSG00000081721 | DUSP12 |
| ENSG00000158050 | DUSP2 |
| ENSG00000158716 | DUSP23 |
| ENSG00000188542 | DUSP28 |
| ENSG00000108861 | DUSP3 |
| ENSG00000120875 | DUSP4 |
| ENSG00000139318 | DUSP6 |
| ENSG00000130829 | DUSP9 |
| ENSG00000144635 | DYNC1LI1 |
| ENSG00000135720 | DYNC1LI2 |
| ENSG00000135636 | DYSF |
| ENSG00000221818 | EBF2 |
| ENSG00000145194 | ECE2 |
| ENSG00000171551 | ECEL1 |
| ENSG00000121310 | ECHDC2 |
| ENSG00000167969 | ECI1 |
| ENSG00000143369 | ECM1 |
| ENSG00000088298 | EDEM2 |
| ENSG00000164176 | EDIL3 |

| | |
|---|---|
| ENSG00000136160 | EDNRB |
| ENSG00000156508 | EEF1A1 |
| ENSG00000103319 | EEF2K |
| ENSG00000172638 | EFEMP2 |
| ENSG00000169242 | EFNA1 |
| ENSG00000184349 | EFNA5 |
| ENSG00000108947 | EFNB3 |
| ENSG00000100842 | EFS |
| ENSG00000172889 | EGFL7 |
| ENSG00000146648 | EGFR |
| ENSG00000269858 | EGLN2 |
| ENSG00000135625 | EGR4 |
| ENSG00000024422 | EHD2 |
| ENSG00000175376 | EIF1AD |
| ENSG00000173674 | EIF1AX |
| ENSG00000055332 | EIF2AK2 |
| ENSG00000134001 | EIF2S1 |
| ENSG00000130811 | EIF3G |
| ENSG00000084623 | EIF3I |
| ENSG00000243056 | EIF4EBP3 |
| ENSG00000106682 | EIF4H |
| ENSG00000197561 | ELANE |
| ENSG00000162374 | ELAVL4 |
| ENSG00000163435 | ELF3 |
| ENSG00000105656 | ELL |
| ENSG00000128886 | ELL3 |
| ENSG00000062598 | ELMO2 |
| ENSG00000130165 | ELOF1 |
| ENSG00000066322 | ELOVL1 |
| ENSG00000183798 | EMILIN3 |
| ENSG00000134531 | EMP1 |

| | |
|---|---|
| ENSG00000142227 | EMP3 |
| ENSG00000170370 | EMX2 |
| ENSG00000167136 | ENDOG |
| ENSG00000111405 | ENDOU |
| ENSG00000106991 | ENG |
| ENSG00000111674 | ENO2 |
| ENSG00000182156 | ENPP7 |
| ENSG00000197586 | ENTPD6 |
| ENSG00000159023 | EPB41 |
| ENSG00000088367 | EPB41L1 |
| ENSG00000166947 | EPB42 |
| ENSG00000116106 | EPHA4 |
| ENSG00000133216 | EPHB2 |
| ENSG00000196411 | EPHB4 |
| ENSG00000072134 | EPN2 |
| ENSG00000261150 | EPPK1 |
| ENSG00000127527 | EPS15L1 |
| ENSG00000177106 | EPS8L2 |
| ENSG00000198758 | EPS8L3 |
| ENSG00000133106 | EPSTI1 |
| ENSG00000082805 | ERC1 |
| ENSG00000104884 | ERCC2 |
| ENSG00000049167 | ERCC8 |
| ENSG00000105722 | ERF |
| ENSG00000087502 | ERGIC2 |
| ENSG00000204978 | ERICH4 |
| ENSG00000177459 | ERICH5 |
| ENSG00000147475 | ERLIN2 |
| ENSG00000164010 | ERMAP |
| ENSG00000089248 | ERP29 |
| ENSG00000187017 | ESPN |

| | |
|---|---|
| ENSG00000139641 | ESYT1 |
| ENSG00000143845 | ETNK2 |
| ENSG00000157557 | ETS2 |
| ENSG00000105672 | ETV2 |
| ENSG00000175832 | ETV4 |
| ENSG00000196405 | EVL |
| ENSG00000167880 | EVPL |
| ENSG00000123737 | EXOSC9 |
| ENSG00000158008 | EXTL1 |
| ENSG00000158161 | EYA3 |
| ENSG00000112319 | EYA4 |
| ENSG00000106462 | EZH2 |
| ENSG00000131187 | F12 |
| ENSG00000180210 | F2 |
| ENSG00000181104 | F2R |
| ENSG00000134824 | FADS2 |
| ENSG00000185104 | FAF1 |
| ENSG00000184083 | FAM120C |
| ENSG00000187866 | FAM122A |
| ENSG00000175182 | FAM131A |
| ENSG00000159784 | FAM131B |
| ENSG00000051009 | FAM160A2 |
| ENSG00000152102 | FAM168B |
| ENSG00000189320 | FAM180A |
| ENSG00000104059 | FAM189A1 |
| ENSG00000116199 | FAM20B |
| ENSG00000177706 | FAM20C |
| ENSG00000174137 | FAM53A |
| ENSG00000120709 | FAM53C |
| ENSG00000180043 | FAM71E2 |
| ENSG00000196550 | FAM72A |

| | |
|---|---|
| ENSG00000147689 | FAM83A |
| ENSG00000180921 | FAM83H |
| ENSG00000181544 | FANCB |
| ENSG00000197601 | FAR1 |
| ENSG00000064763 | FAR2 |
| ENSG00000117560 | FASLG |
| ENSG00000164896 | FASTK |
| ENSG00000188878 | FBF1 |
| ENSG00000105202 | FBL |
| ENSG00000162458 | FBLIM1 |
| ENSG00000166147 | FBN1 |
| ENSG00000138829 | FBN2 |
| ENSG00000165140 | FBP1 |
| ENSG00000171823 | FBXL14 |
| ENSG00000155034 | FBXL18 |
| ENSG00000147364 | FBXO25 |
| ENSG00000132879 | FBXO44 |
| ENSG00000116663 | FBXO6 |
| ENSG00000107829 | FBXW4 |
| ENSG00000159069 | FBXW5 |
| ENSG00000174989 | FBXW8 |
| ENSG00000072694 | FCGR2B |
| ENSG00000137478 | FCHSD2 |
| ENSG00000160752 | FDPS |
| ENSG00000168496 | FEN1 |
| ENSG00000182511 | FES |
| ENSG00000090512 | FETUB |
| ENSG00000149557 | FEZ1 |
| ENSG00000171055 | FEZ2 |
| ENSG00000146192 | FGD2 |
| ENSG00000113578 | FGF1 |

| | |
|---|---|
| ENSG00000161958 | FGF11 |
| ENSG00000129682 | FGF13 |
| ENSG00000156427 | FGF18 |
| ENSG00000162344 | FGF19 |
| ENSG00000138685 | FGF2 |
| ENSG00000078579 | FGF20 |
| ENSG00000102678 | FGF9 |
| ENSG00000174721 | FGFBP3 |
| ENSG00000160867 | FGFR4 |
| ENSG00000000938 | FGR |
| ENSG00000137460 | FHDC1 |
| ENSG00000183733 | FIGLA |
| ENSG00000139914 | FITM1 |
| ENSG00000173486 | FKBP2 |
| ENSG00000122025 | FLT3 |
| ENSG00000090554 | FLT3LG |
| ENSG00000248905 | FMN1 |
| ENSG00000115414 | FN1 |
| ENSG00000167363 | FN3K |
| ENSG00000102531 | FNDC3A |
| ENSG00000170608 | FOXA3 |
| ENSG00000186564 | FOXD2 |
| ENSG00000137273 | FOXF2 |
| ENSG00000168269 | FOXI1 |
| ENSG00000129654 | FOXJ1 |
| ENSG00000176678 | FOXL1 |
| ENSG00000109101 | FOXN1 |
| ENSG00000139445 | FOXN4 |
| ENSG00000118689 | FOXO3 |
| ENSG00000136877 | FPGS |
| ENSG00000171877 | FRMD5 |

| | |
|---|---|
| ENSG00000139926 | FRMD6 |
| ENSG00000137218 | FRS3 |
| ENSG00000162998 | FRZB |
| ENSG00000075618 | FSCN1 |
| ENSG00000186765 | FSCN2 |
| ENSG00000106701 | FSD1L |
| ENSG00000134363 | FST |
| ENSG00000160282 | FTCD |
| ENSG00000087086 | FTL |
| ENSG00000174951 | FUT1 |
| ENSG00000172461 | FUT9 |
| ENSG00000266964 | FXYD1 |
| ENSG00000150201 | FXYD4 |
| ENSG00000089327 | FXYD5 |
| ENSG00000221946 | FXYD7 |
| ENSG00000163820 | FYCO1 |
| ENSG00000122068 | FYTTD1 |
| ENSG00000163251 | FZD5 |
| ENSG00000177283 | FZD8 |
| ENSG00000105325 | FZR1 |
| ENSG00000171298 | GAA |
| ENSG00000166206 | GABRB3 |
| ENSG00000187730 | GABRD |
| ENSG00000094755 | GABRP |
| ENSG00000128683 | GAD1 |
| ENSG00000136750 | GAD2 |
| ENSG00000069482 | GAL |
| ENSG00000197093 | GAL3ST4 |
| ENSG00000117308 | GALE |
| ENSG00000119514 | GALNT12 |
| ENSG00000144278 | GALNT13 |

| | |
|---|---|
| ENSG00000158089 | GALNT14 |
| ENSG00000109586 | GALNT7 |
| ENSG00000128310 | GALR3 |
| ENSG00000213930 | GALT |
| ENSG00000111640 | GAPDH |
| ENSG00000183087 | GAS6 |
| ENSG00000102145 | GATA1 |
| ENSG00000107862 | GBF1 |
| ENSG00000183347 | GBP6 |
| ENSG00000164900 | GBX1 |
| ENSG00000168505 | GBX2 |
| ENSG00000105607 | GCDH |
| ENSG00000215644 | GCGR |
| ENSG00000137880 | GCHFR |
| ENSG00000084734 | GCKR |
| ENSG00000111846 | GCNT2 |
| ENSG00000124194 | GDAP1L1 |
| ENSG00000006007 | GDE1 |
| ENSG00000130283 | GDF1 |
| ENSG00000130513 | GDF15 |
| ENSG00000125965 | GDF5 |
| ENSG00000156466 | GDF6 |
| ENSG00000057608 | GDI2 |
| ENSG00000168621 | GDNF |
| ENSG00000164949 | GEM |
| ENSG00000131095 | GFAP |
| ENSG00000162676 | GFI1 |
| ENSG00000131459 | GFPT2 |
| ENSG00000168546 | GFRA2 |
| ENSG00000146013 | GFRA3 |
| ENSG00000100083 | GGA1 |

| | |
|---|---|
| ENSG00000152904 | GGPS1 |
| ENSG00000099998 | GGT5 |
| ENSG00000167741 | GGT6 |
| ENSG00000157017 | GHRL |
| ENSG00000281887 | GIMAP1-GIMAP5 |
| ENSG00000108262 | GIT1 |
| ENSG00000121743 | GJA3 |
| ENSG00000187513 | GJA4 |
| ENSG00000189433 | GJB4 |
| ENSG00000189280 | GJB5 |
| ENSG00000198835 | GJC2 |
| ENSG00000196475 | GK2 |
| ENSG00000149328 | GLB1L2 |
| ENSG00000174842 | GLMN |
| ENSG00000112164 | GLP1R |
| ENSG00000145451 | GLRA3 |
| ENSG00000109738 | GLRB |
| ENSG00000173221 | GLRX |
| ENSG00000151948 | GLT1D1 |
| ENSG00000204007 | GLT6D1 |
| ENSG00000120820 | GLT8D2 |
| ENSG00000140632 | GLYR1 |
| ENSG00000144591 | GMPPA |
| ENSG00000173540 | GMPPB |
| ENSG00000088256 | GNA11 |
| ENSG00000156049 | GNA14 |
| ENSG00000065135 | GNAI3 |
| ENSG00000087258 | GNAO1 |
| ENSG00000128266 | GNAZ |
| ENSG00000111664 | GNB3 |
| ENSG00000114450 | GNB4 |

| | |
|---|---|
| ENSG00000127588 | GNG13 |
| ENSG00000162188 | GNG3 |
| ENSG00000111670 | GNPTAB |
| ENSG00000144674 | GOLGA4 |
| ENSG00000135052 | GOLM1 |
| ENSG00000174567 | GOLT1A |
| ENSG00000108433 | GOSR2 |
| ENSG00000179921 | GPBAR1 |
| ENSG00000063660 | GPC1 |
| ENSG00000076716 | GPC4 |
| ENSG00000171723 | GPHN |
| ENSG00000165370 | GPR101 |
| ENSG00000147262 | GPR119 |
| ENSG00000183484 | GPR132 |
| ENSG00000077585 | GPR137B |
| ENSG00000257008 | GPR142 |
| ENSG00000164849 | GPR146 |
| ENSG00000178015 | GPR150 |
| ENSG00000175697 | GPR156 |
| ENSG00000151025 | GPR158 |
| ENSG00000204882 | GPR20 |
| ENSG00000181773 | GPR3 |
| ENSG00000170075 | GPR37L1 |
| ENSG00000135973 | GPR45 |
| ENSG00000119714 | GPR68 |
| ENSG00000123901 | GPR83 |
| ENSG00000170412 | GPRC5C |
| ENSG00000185477 | GPRIN3 |
| ENSG00000167701 | GPT |
| ENSG00000166123 | GPT2 |
| ENSG00000167468 | GPX4 |

| | |
|---|---|
| ENSG00000198704 | GPX6 |
| ENSG00000075240 | GRAMD4 |
| ENSG00000180875 | GREM2 |
| ENSG00000137106 | GRHPR |
| ENSG00000182771 | GRID1 |
| ENSG00000149403 | GRIK4 |
| ENSG00000105737 | GRIK5 |
| ENSG00000105464 | GRIN2D |
| ENSG00000185974 | GRK1 |
| ENSG00000198873 | GRK5 |
| ENSG00000109519 | GRPEL1 |
| ENSG00000126010 | GRPR |
| ENSG00000105447 | GRWD1 |
| ENSG00000167914 | GSDMA |
| ENSG00000082701 | GSK3B |
| ENSG00000148180 | GSN |
| ENSG00000168765 | GSTM4 |
| ENSG00000134201 | GSTM5 |
| ENSG00000148834 | GSTO1 |
| ENSG00000169840 | GSX1 |
| ENSG00000180613 | GSX2 |
| ENSG00000130299 | GTPBP3 |
| ENSG00000178605 | GTPBP6 |
| ENSG00000178605 | GTPBP6 |
| ENSG00000075218 | GTSE1 |
| ENSG00000197273 | GUCA2A |
| ENSG00000044012 | GUCA2B |
| ENSG00000151233 | GXYLT1 |
| ENSG00000125812 | GZF1 |
| ENSG00000197540 | GZMM |
| ENSG00000131373 | HACL1 |

| | |
|---|---|
| ENSG00000084110 | HAL |
| ENSG00000140511 | HAPLN3 |
| ENSG00000213397 | HAUS7 |
| ENSG00000163517 | HDAC11 |
| ENSG00000108840 | HDAC5 |
| ENSG00000140287 | HDC |
| ENSG00000119285 | HEATR1 |
| ENSG00000148634 | HERC4 |
| ENSG00000069812 | HES2 |
| ENSG00000144485 | HES6 |
| ENSG00000179111 | HES7 |
| ENSG00000135547 | HEY2 |
| ENSG00000163909 | HEYL |
| ENSG00000162669 | HFM1 |
| ENSG00000185359 | HGS |
| ENSG00000054392 | HHAT |
| ENSG00000010282 | HHATL |
| ENSG00000152804 | HHEX |
| ENSG00000169635 | HIC2 |
| ENSG00000169567 | HINT1 |
| ENSG00000163349 | HIPK1 |
| ENSG00000010818 | HIVEP2 |
| ENSG00000206503 | HLA-A |
| ENSG00000204257 | HLA-DMA |
| ENSG00000179344 | HLA-DQB1 |
| ENSG00000137309 | HMGA1 |
| ENSG00000112972 | HMGCS1 |
| ENSG00000135100 | HNF1A |
| ENSG00000101076 | HNF4A |
| ENSG00000104824 | HNRNPL |
| ENSG00000095066 | HOOK2 |

| | |
|---|---|
| ENSG00000143452 | HORMAD1 |
| ENSG00000253293 | HOXA10 |
| ENSG00000106031 | HOXA13 |
| ENSG00000106004 | HOXA5 |
| ENSG00000106006 | HOXA6 |
| ENSG00000120094 | HOXB1 |
| ENSG00000159184 | HOXB13 |
| ENSG00000173917 | HOXB2 |
| ENSG00000182742 | HOXB4 |
| ENSG00000180818 | HOXC10 |
| ENSG00000172789 | HOXC5 |
| ENSG00000197757 | HOXC6 |
| ENSG00000128710 | HOXD10 |
| ENSG00000170178 | HOXD12 |
| ENSG00000128709 | HOXD9 |
| ENSG00000168453 | HR |
| ENSG00000196639 | HRH1 |
| ENSG00000134489 | HRH4 |
| ENSG00000153936 | HS2ST1 |
| ENSG00000162040 | HS3ST6 |
| ENSG00000230989 | HSBP1 |
| ENSG00000176387 | HSD11B2 |
| ENSG00000108786 | HSD17B1 |
| ENSG00000099377 | HSD3B7 |
| ENSG00000102878 | HSF4 |
| ENSG00000166598 | HSP90B1 |
| ENSG00000165868 | HSPA12A |
| ENSG00000132622 | HSPA12B |
| ENSG00000044574 | HSPA5 |
| ENSG00000173641 | HSPB7 |
| ENSG00000152137 | HSPB8 |

| | |
|---|---|
| ENSG00000120694 | HSPH1 |
| ENSG00000166736 | HTR3A |
| ENSG00000136273 | HUS1 |
| ENSG00000086758 | HUWE1 |
| ENSG00000122986 | HVCN1 |
| ENSG00000090339 | ICAM1 |
| ENSG00000105371 | ICAM4 |
| ENSG00000105376 | ICAM5 |
| ENSG00000182054 | IDH2 |
| ENSG00000166411 | IDH3A |
| ENSG00000068079 | IFI35 |
| ENSG00000204010 | IFIT1B |
| ENSG00000159128 | IFNGR2 |
| ENSG00000214706 | IFRD2 |
| ENSG00000138002 | IFT172 |
| ENSG00000118096 | IFT46 |
| ENSG00000159217 | IGF2BP1 |
| ENSG00000073792 | IGF2BP2 |
| ENSG00000099769 | IGFALS |
| ENSG00000141753 | IGFBP4 |
| ENSG00000188624 | IGFL3 |
| ENSG00000117154 | IGSF21 |
| ENSG00000143061 | IGSF3 |
| ENSG00000163501 | IHH |
| ENSG00000104365 | IKBKB |
| ENSG00000110324 | IL10RA |
| ENSG00000243646 | IL10RB |
| ENSG00000095752 | IL11 |
| ENSG00000137070 | IL11RA |
| ENSG00000113302 | IL12B |
| ENSG00000096996 | IL12RB1 |

| | |
|---|---|
| ENSG00000164136 | IL15 |
| ENSG00000134470 | IL15RA |
| ENSG00000172349 | IL16 |
| ENSG00000124391 | IL17C |
| ENSG00000137496 | IL18BP |
| ENSG00000115604 | IL18R1 |
| ENSG00000016402 | IL20RA |
| ENSG00000174564 | IL20RB |
| ENSG00000103522 | IL21R |
| ENSG00000166090 | IL25 |
| ENSG00000100385 | IL2RB |
| ENSG00000147168 | IL2RG |
| ENSG00000157368 | IL34 |
| ENSG00000104951 | IL4I1 |
| ENSG00000104432 | IL7 |
| ENSG00000124334 | IL9R |
| ENSG00000124334 | IL9R |
| ENSG00000129351 | ILF3 |
| ENSG00000132305 | IMMT |
| ENSG00000133731 | IMPA1 |
| ENSG00000203485 | INF2 |
| ENSG00000123999 | INHA |
| ENSG00000139269 | INHBE |
| ENSG00000241644 | INMT |
| ENSG00000128908 | INO80 |
| ENSG00000115274 | INO80B |
| ENSG00000153391 | INO80C |
| ENSG00000068383 | INPP5A |
| ENSG00000198825 | INPP5F |
| ENSG00000185133 | INPP5J |
| ENSG00000186480 | INSIG1 |

| | |
|---|---|
| ENSG00000125629 | INSIG2 |
| ENSG00000102786 | INTS6 |
| ENSG00000119509 | INVS |
| ENSG00000161896 | IP6K3 |
| ENSG00000196497 | IPO4 |
| ENSG00000065150 | IPO5 |
| ENSG00000160051 | IQCC |
| ENSG00000125347 | IRF1 |
| ENSG00000126456 | IRF3 |
| ENSG00000137265 | IRF4 |
| ENSG00000169047 | IRS1 |
| ENSG00000170549 | IRX1 |
| ENSG00000177508 | IRX3 |
| ENSG00000135070 | ISCA1 |
| ENSG00000172183 | ISG20 |
| ENSG00000161638 | ITGA5 |
| ENSG00000150093 | ITGB1 |
| ENSG00000055957 | ITIH1 |
| ENSG00000162267 | ITIH3 |
| ENSG00000055955 | ITIH4 |
| ENSG00000078596 | ITM2A |
| ENSG00000100605 | ITPK1 |
| ENSG00000123104 | ITPR2 |
| ENSG00000182264 | IZUMO1 |
| ENSG00000105639 | JAK3 |
| ENSG00000161999 | JMJD8 |
| ENSG00000161677 | JOSD2 |
| ENSG00000154118 | JPH3 |
| ENSG00000092051 | JPH4 |
| ENSG00000234616 | JRK |
| ENSG00000130522 | JUND |

| | |
|---|---|
| ENSG00000189337 | KAZN |
| ENSG00000111262 | KCNA1 |
| ENSG00000177272 | KCNA3 |
| ENSG00000166006 | KCNC2 |
| ENSG00000102057 | KCND1 |
| ENSG00000184408 | KCND2 |
| ENSG00000171385 | KCND3 |
| ENSG00000159197 | KCNE2 |
| ENSG00000055118 | KCNH2 |
| ENSG00000173826 | KCNH6 |
| ENSG00000187486 | KCNJ11 |
| ENSG00000153822 | KCNJ16 |
| ENSG00000123700 | KCNJ2 |
| ENSG00000157542 | KCNJ6 |
| ENSG00000182450 | KCNK4 |
| ENSG00000099337 | KCNK6 |
| ENSG00000173338 | KCNK7 |
| ENSG00000156113 | KCNMA1 |
| ENSG00000145936 | KCNMB1 |
| ENSG00000135643 | KCNMB4 |
| ENSG00000105642 | KCNN1 |
| ENSG00000104783 | KCNN4 |
| ENSG00000053918 | KCNQ1 |
| ENSG00000184156 | KCNQ3 |
| ENSG00000135253 | KCP |
| ENSG00000180901 | KCTD2 |
| ENSG00000167977 | KCTD5 |
| ENSG00000168301 | KCTD6 |
| ENSG00000104756 | KCTD9 |
| ENSG00000105438 | KDELR1 |
| ENSG00000089094 | KDM2B |

| | |
|---|---|
| ENSG00000147050 | KDM6A |
| ENSG00000132510 | KDM6B |
| ENSG00000128052 | KDR |
| ENSG00000121774 | KHDRBS1 |
| ENSG00000138030 | KHK |
| ENSG00000100441 | KHNYN |
| ENSG00000198920 | KIAA0753 |
| ENSG00000196123 | KIAA0895L |
| ENSG00000100364 | KIAA0930 |
| ENSG00000162522 | KIAA1522 |
| ENSG00000117245 | KIF17 |
| ENSG00000129250 | KIF1C |
| ENSG00000068796 | KIF2A |
| ENSG00000155980 | KIF5A |
| ENSG00000164627 | KIF6 |
| ENSG00000167702 | KIFC2 |
| ENSG00000116014 | KISS1R |
| ENSG00000104892 | KLC3 |
| ENSG00000155090 | KLF10 |
| ENSG00000169926 | KLF13 |
| ENSG00000163884 | KLF15 |
| ENSG00000171872 | KLF17 |
| ENSG00000179023 | KLHDC7A |
| ENSG00000185909 | KLHDC8B |
| ENSG00000114648 | KLHL18 |
| ENSG00000167487 | KLHL26 |
| ENSG00000186231 | KLHL32 |
| ENSG00000135686 | KLHL36 |
| ENSG00000129451 | KLK10 |
| ENSG00000167757 | KLK11 |
| ENSG00000169035 | KLK7 |

| | |
|---|---|
| ENSG00000129455 | KLK8 |
| ENSG00000213022 | KLK9 |
| ENSG00000188883 | KLRG2 |
| ENSG00000171798 | KNDC1 |
| ENSG00000118162 | KPTN |
| ENSG00000133619 | KRBA1 |
| ENSG00000183762 | KREMEN1 |
| ENSG00000131650 | KREMEN2 |
| ENSG00000001631 | KRIT1 |
| ENSG00000167768 | KRT1 |
| ENSG00000171346 | KRT15 |
| ENSG00000186832 | KRT16 |
| ENSG00000128422 | KRT17 |
| ENSG00000171345 | KRT19 |
| ENSG00000094796 | KRT31 |
| ENSG00000135480 | KRT7 |
| ENSG00000139648 | KRT71 |
| ENSG00000186049 | KRT73 |
| ENSG00000170454 | KRT75 |
| ENSG00000185069 | KRT76 |
| ENSG00000196156 | KRTAP4-3 |
| ENSG00000157992 | KRTCAP3 |
| ENSG00000198910 | L1CAM |
| ENSG00000147592 | LACTB2 |
| ENSG00000089692 | LAG3 |
| ENSG00000167531 | LALBA |
| ENSG00000101680 | LAMA1 |
| ENSG00000005893 | LAMP2 |
| ENSG00000213658 | LAT |
| ENSG00000213398 | LCAT |
| ENSG00000187922 | LCN10 |

| | |
|---|---|
| ENSG00000184925 | LCN12 |
| ENSG00000148346 | LCN2 |
| ENSG00000148386 | LCN9 |
| ENSG00000198728 | LDB1 |
| ENSG00000134333 | LDHA |
| ENSG00000182195 | LDOC1 |
| ENSG00000243709 | LEFTY1 |
| ENSG00000161904 | LEMD2 |
| ENSG00000163352 | LENEP |
| ENSG00000275183 | LENG9 |
| ENSG00000174697 | LEP |
| ENSG00000213625 | LEPROT |
| ENSG00000205076 | LGALS7 |
| ENSG00000178934 | LGALS7B |
| ENSG00000168481 | LGI3 |
| ENSG00000153902 | LGI4 |
| ENSG00000139292 | LGR5 |
| ENSG00000104826 | LHB |
| ENSG00000089116 | LHX5 |
| ENSG00000106852 | LHX6 |
| ENSG00000128342 | LIF |
| ENSG00000105486 | LIG1 |
| ENSG00000105370 | LIM2 |
| ENSG00000064042 | LIMCH1 |
| ENSG00000203896 | LIME1 |
| ENSG00000106683 | LIMK1 |
| ENSG00000169756 | LIMS1 |
| ENSG00000205659 | LIN52 |
| ENSG00000104863 | LIN7B |
| ENSG00000107798 | LIPA |
| ENSG00000189067 | LITAF |

| | |
|---|---|
| ENSG00000073350 | LLGL2 |
| ENSG00000074695 | LMAN1 |
| ENSG00000113368 | LMNB1 |
| ENSG00000142235 | LMTK3 |
| ENSG00000136944 | LMX1B |
| ENSG00000129038 | LOXL1 |
| ENSG00000134013 | LOXL2 |
| ENSG00000198121 | LPAR1 |
| ENSG00000064547 | LPAR2 |
| ENSG00000134324 | LPIN1 |
| ENSG00000136141 | LRCH1 |
| ENSG00000128011 | LRFN1 |
| ENSG00000173621 | LRFN4 |
| ENSG00000165379 | LRFN5 |
| ENSG00000171236 | LRG1 |
| ENSG00000139263 | LRIG3 |
| ENSG00000134569 | LRP4 |
| ENSG00000162337 | LRP5 |
| ENSG00000137269 | LRRC1 |
| ENSG00000163827 | LRRC2 |
| ENSG00000172731 | LRRC20 |
| ENSG00000254402 | LRRC24 |
| ENSG00000160233 | LRRC3 |
| ENSG00000183908 | LRRC55 |
| ENSG00000108829 | LRRC59 |
| ENSG00000136802 | LRRC8A |
| ENSG00000171492 | LRRC8D |
| ENSG00000124831 | LRRFIP1 |
| ENSG00000166159 | LRTM2 |
| ENSG00000149657 | LSM14B |
| ENSG00000130592 | LSP1 |

| | |
|---|---|
| ENSG00000105699 | LSR |
| ENSG00000160285 | LSS |
| ENSG00000226979 | LTA |
| ENSG00000111144 | LTA4H |
| ENSG00000227507 | LTB |
| ENSG00000213903 | LTB4R |
| ENSG00000213906 | LTB4R2 |
| ENSG00000049323 | LTBP1 |
| ENSG00000111321 | LTBR |
| ENSG00000213316 | LTC4S |
| ENSG00000160932 | LY6E |
| ENSG00000104903 | LYL1 |
| ENSG00000124466 | LYPD3 |
| ENSG00000150556 | LYPD6B |
| ENSG00000011009 | LYPLA2 |
| ENSG00000143353 | LYPLAL1 |
| ENSG00000102897 | LYRM1 |
| ENSG00000083099 | LYRM2 |
| ENSG00000214113 | LYRM4 |
| ENSG00000186687 | LYRM7 |
| ENSG00000183060 | LYSMD4 |
| ENSG00000110514 | MADD |
| ENSG00000178573 | MAF |
| ENSG00000105695 | MAG |
| ENSG00000179222 | MAGED1 |
| ENSG00000172005 | MAL |
| ENSG00000196782 | MAML3 |
| ENSG00000140400 | MAN2C1 |
| ENSG00000189221 | MAOA |
| ENSG00000076984 | MAP2K7 |
| ENSG00000095015 | MAP3K1 |

| | |
|---|---|
| ENSG00000006062 | MAP3K14 |
| ENSG00000169967 | MAP3K2 |
| ENSG00000198909 | MAP3K3 |
| ENSG00000100030 | MAPK1 |
| ENSG00000188130 | MAPK12 |
| ENSG00000156711 | MAPK13 |
| ENSG00000141639 | MAPK4 |
| ENSG00000121653 | MAPK8IP1 |
| ENSG00000138834 | MAPK8IP3 |
| ENSG00000089022 | MAPKAPK5 |
| ENSG00000186868 | MAPT |
| ENSG00000140832 | MARVELD3 |
| ENSG00000086015 | MAST2 |
| ENSG00000132561 | MATN2 |
| ENSG00000132031 | MATN3 |
| ENSG00000124159 | MATN4 |
| ENSG00000088888 | MAVS |
| ENSG00000176055 | MBLAC2 |
| ENSG00000076770 | MBNL3 |
| ENSG00000076706 | MCAM |
| ENSG00000100294 | MCAT |
| ENSG00000153898 | MCOLN2 |
| ENSG00000110492 | MDK |
| ENSG00000082212 | ME2 |
| ENSG00000124733 | MEA1 |
| ENSG00000085276 | MECOM |
| ENSG00000184634 | MED12 |
| ENSG00000180182 | MED14 |
| ENSG00000175221 | MED16 |
| ENSG00000130772 | MED18 |
| ENSG00000164758 | MED30 |

| | |
|---|---|
| ENSG00000068305 | MEF2A |
| ENSG00000145794 | MEGF10 |
| ENSG00000169519 | METTL15 |
| ENSG00000127804 | METTL16 |
| ENSG00000165792 | METTL17 |
| ENSG00000087995 | METTL2A |
| ENSG00000101574 | METTL4 |
| ENSG00000185432 | METTL7A |
| ENSG00000123600 | METTL8 |
| ENSG00000254726 | MEX3A |
| ENSG00000181588 | MEX3D |
| ENSG00000117122 | MFAP2 |
| ENSG00000205639 | MFSD2B |
| ENSG00000167700 | MFSD3 |
| ENSG00000151690 | MFSD6 |
| ENSG00000185156 | MFSD6L |
| ENSG00000161013 | MGAT4B |
| ENSG00000074416 | MGLL |
| ENSG00000164877 | MICALL2 |
| ENSG00000167470 | MIDN |
| ENSG00000155545 | MIER3 |
| ENSG00000164654 | MIOS |
| ENSG00000027001 | MIPEP |
| ENSG00000185155 | MIXL1 |
| ENSG00000099875 | MKNK2 |
| ENSG00000130382 | MLLT1 |
| ENSG00000108788 | MLX |
| ENSG00000009950 | MLXIPL |
| ENSG00000151611 | MMAA |
| ENSG00000136297 | MMD2 |
| ENSG00000099953 | MMP11 |

| | |
|---|---|
| ENSG00000157227 | MMP14 |
| ENSG00000154485 | MMP21 |
| ENSG00000189409 | MMP23B |
| ENSG00000008516 | MMP25 |
| ENSG00000271447 | MMP28 |
| ENSG00000121211 | MND1 |
| ENSG00000130675 | MNX1 |
| ENSG00000182208 | MOB2 |
| ENSG00000115275 | MOGS |
| ENSG00000133131 | MORC4 |
| ENSG00000188010 | MORN2 |
| ENSG00000101928 | MOSPD1 |
| ENSG00000103152 | MPG |
| ENSG00000158186 | MRAS |
| ENSG00000184350 | MRGPRE |
| ENSG00000172935 | MRGPRF |
| ENSG00000262814 | MRPL12 |
| ENSG00000158042 | MRPL17 |
| ENSG00000143314 | MRPL24 |
| ENSG00000243147 | MRPL33 |
| ENSG00000116221 | MRPL37 |
| ENSG00000172590 | MRPL52 |
| ENSG00000204822 | MRPL53 |
| ENSG00000162910 | MRPL55 |
| ENSG00000135972 | MRPS9 |
| ENSG00000053372 | MRTO4 |
| ENSG00000156738 | MS4A1 |
| ENSG00000172689 | MS4A10 |
| ENSG00000102854 | MSLN |
| ENSG00000147065 | MSN |
| ENSG00000149480 | MTA2 |

| | |
|---|---|
| ENSG00000057935 | MTA3 |
| ENSG00000147649 | MTDH |
| ENSG00000242114 | MTFP1 |
| ENSG00000100714 | MTHFD1 |
| ENSG00000177000 | MTHFR |
| ENSG00000103248 | MTHFSD |
| ENSG00000014914 | MTMR11 |
| ENSG00000112031 | MTRF1L |
| ENSG00000117983 | MUC5B |
| ENSG00000184956 | MUC6 |
| ENSG00000172732 | MUS81 |
| ENSG00000013364 | MVP |
| ENSG00000213347 | MXD3 |
| ENSG00000132382 | MYBBP1A |
| ENSG00000134571 | MYBPC3 |
| ENSG00000221986 | MYBPHL |
| ENSG00000134323 | MYCN |
| ENSG00000172936 | MYD88 |
| ENSG00000111049 | MYF5 |
| ENSG00000133026 | MYH10 |
| ENSG00000105357 | MYH14 |
| ENSG00000197616 | MYH6 |
| ENSG00000092054 | MYH7 |
| ENSG00000106436 | MYL10 |
| ENSG00000101608 | MYL12A |
| ENSG00000092841 | MYL6 |
| ENSG00000196465 | MYL6B |
| ENSG00000091536 | MYO15A |
| ENSG00000196535 | MYO18A |
| ENSG00000278259 | MYO19 |
| ENSG00000176658 | MYO1D |

| | |
|---|---|
| ENSG00000137474 | MYO7A |
| ENSG00000129152 | MYOD1 |
| ENSG00000101605 | MYOM1 |
| ENSG00000244754 | N4BP2L2 |
| ENSG00000145911 | N4BP3 |
| ENSG00000168060 | NAALADL1 |
| ENSG00000148411 | NACC2 |
| ENSG00000159593 | NAE1 |
| ENSG00000198951 | NAGA |
| ENSG00000161653 | NAGS |
| ENSG00000205531 | NAP1L4 |
| ENSG00000131400 | NAPSA |
| ENSG00000141562 | NARF |
| ENSG00000132780 | NASP |
| ENSG00000090971 | NAT14 |
| ENSG00000134369 | NAV1 |
| ENSG00000130287 | NCAN |
| ENSG00000146918 | NCAPG2 |
| ENSG00000025770 | NCAPH2 |
| ENSG00000158517 | NCF1 |
| ENSG00000176771 | NCKAP5 |
| ENSG00000213672 | NCKIPSD |
| ENSG00000125912 | NCLN |
| ENSG00000196498 | NCOR2 |
| ENSG00000166579 | NDEL1 |
| ENSG00000165795 | NDRG2 |
| ENSG00000103034 | NDRG4 |
| ENSG00000139180 | NDUFA9 |
| ENSG00000165264 | NDUFB6 |
| ENSG00000023228 | NDUFS1 |
| ENSG00000115286 | NDUFS7 |

| | |
|---|---|
| ENSG00000103154 | NECAB2 |
| ENSG00000111859 | NEDD9 |
| ENSG00000277586 | NEFL |
| ENSG00000140398 | NEIL1 |
| ENSG00000119408 | NEK6 |
| ENSG00000184613 | NELL2 |
| ENSG00000173848 | NET1 |
| ENSG00000166342 | NETO1 |
| ENSG00000162139 | NEU3 |
| ENSG00000215041 | NEURL4 |
| ENSG00000181965 | NEUROG1 |
| ENSG00000196712 | NF1 |
| ENSG00000235568 | NFAM1 |
| ENSG00000163531 | NFASC |
| ENSG00000131196 | NFATC1 |
| ENSG00000101096 | NFATC2 |
| ENSG00000100968 | NFATC4 |
| ENSG00000147862 | NFIB |
| ENSG00000141905 | NFIC |
| ENSG00000165030 | NFIL3 |
| ENSG00000077150 | NFKB2 |
| ENSG00000204498 | NFKBIL1 |
| ENSG00000064300 | NGFR |
| ENSG00000171786 | NHLH1 |
| ENSG00000187566 | NHLRC1 |
| ENSG00000257108 | NHLRC4 |
| ENSG00000170113 | NIPA1 |
| ENSG00000136783 | NIPSNAP3A |
| ENSG00000186416 | NKRF |
| ENSG00000119919 | NKX2-3 |
| ENSG00000136327 | NKX2-8 |

| | |
|---|---|
| ENSG00000163623 | NKX6-1 |
| ENSG00000169760 | NLGN1 |
| ENSG00000169992 | NLGN2 |
| ENSG00000103024 | NME3 |
| ENSG00000103202 | NME4 |
| ENSG00000171596 | NMUR1 |
| ENSG00000053438 | NNAT |
| ENSG00000140939 | NOL3 |
| ENSG00000111641 | NOP2 |
| ENSG00000164867 | NOS3 |
| ENSG00000104967 | NOVA2 |
| ENSG00000130751 | NPAS1 |
| ENSG00000151322 | NPAS3 |
| ENSG00000174576 | NPAS4 |
| ENSG00000183979 | NPB |
| ENSG00000015520 | NPC1L1 |
| ENSG00000139574 | NPFF |
| ENSG00000131697 | NPHP4 |
| ENSG00000161270 | NPHS1 |
| ENSG00000159899 | NPR2 |
| ENSG00000113389 | NPR3 |
| ENSG00000114388 | NPRL2 |
| ENSG00000164129 | NPY5R |
| ENSG00000174738 | NR1D2 |
| ENSG00000184162 | NR2C2AP |
| ENSG00000278570 | NR2E3 |
| ENSG00000185551 | NR2F2 |
| ENSG00000113580 | NR3C1 |
| ENSG00000123358 | NR4A1 |
| ENSG00000198435 | NRARP |
| ENSG00000091129 | NRCAM |

| | |
|---|---|
| ENSG00000185737 | NRG3 |
| ENSG00000175352 | NRIP3 |
| ENSG00000124785 | NRN1 |
| ENSG00000152954 | NRSN1 |
| ENSG00000171119 | NRTN |
| ENSG00000178694 | NSUN3 |
| ENSG00000135318 | NT5E |
| ENSG00000185652 | NTF3 |
| ENSG00000162068 | NTN3 |
| ENSG00000142233 | NTN5 |
| ENSG00000148053 | NTRK2 |
| ENSG00000101188 | NTSR1 |
| ENSG00000074590 | NUAK1 |
| ENSG00000104805 | NUCB1 |
| ENSG00000106268 | NUDT1 |
| ENSG00000168101 | NUDT16L1 |
| ENSG00000275074 | NUDT18 |
| ENSG00000213965 | NUDT19 |
| ENSG00000167799 | NUDT8 |
| ENSG00000105245 | NUMBL |
| ENSG00000124789 | NUP153 |
| ENSG00000138750 | NUP54 |
| ENSG00000110713 | NUP98 |
| ENSG00000143748 | NVL |
| ENSG00000182575 | NXPH3 |
| ENSG00000182379 | NXPH4 |
| ENSG00000188937 | NYX |
| ENSG00000111335 | OAS2 |
| ENSG00000111331 | OAS3 |
| ENSG00000104044 | OCA2 |
| ENSG00000115758 | ODC1 |

| | |
|---|---|
| ENSG00000136811 | ODF2 |
| ENSG00000177947 | ODF3 |
| ENSG00000182950 | ODF3L1 |
| ENSG00000181781 | ODF3L2 |
| ENSG00000114026 | OGG1 |
| ENSG00000205927 | OLIG2 |
| ENSG00000173391 | OLR1 |
| ENSG00000169856 | ONECUT1 |
| ENSG00000197430 | OPALIN |
| ENSG00000178814 | OPLAH |
| ENSG00000112038 | OPRM1 |
| ENSG00000180708 | OR10K2 |
| ENSG00000196248 | OR10S1 |
| ENSG00000204688 | OR2H1 |
| ENSG00000181609 | OR52D1 |
| ENSG00000184478 | OR56A3 |
| ENSG00000178586 | OR6B3 |
| ENSG00000181803 | OR6S1 |
| ENSG00000258083 | OR9A4 |
| ENSG00000123353 | ORMDL2 |
| ENSG00000135506 | OS9 |
| ENSG00000099985 | OSM |
| ENSG00000115155 | OTOF |
| ENSG00000089723 | OTUB2 |
| ENSG00000115507 | OTX1 |
| ENSG00000186912 | P2RY4 |
| ENSG00000171631 | P2RY6 |
| ENSG00000072682 | P4HA2 |
| ENSG00000170515 | PA2G4 |
| ENSG00000124507 | PACSIN1 |
| ENSG00000142623 | PADI1 |

| | |
|---|---|
| ENSG00000079462 | PAFAH1B3 |
| ENSG00000128050 | PAICS |
| ENSG00000152782 | PANK1 |
| ENSG00000100767 | PAPLN |
| ENSG00000162073 | PAQR4 |
| ENSG00000170915 | PAQR8 |
| ENSG00000188582 | PAQR9 |
| ENSG00000102981 | PARD6A |
| ENSG00000188677 | PARVB |
| ENSG00000138964 | PARVG |
| ENSG00000196844 | PATE2 |
| ENSG00000166889 | PATL1 |
| ENSG00000177425 | PAWR |
| ENSG00000075891 | PAX2 |
| ENSG00000106331 | PAX4 |
| ENSG00000196092 | PAX5 |
| ENSG00000007372 | PAX6 |
| ENSG00000009709 | PAX7 |
| ENSG00000125618 | PAX8 |
| ENSG00000105717 | PBX4 |
| ENSG00000166228 | PCBD1 |
| ENSG00000183570 | PCBP3 |
| ENSG00000090097 | PCBP4 |
| ENSG00000114054 | PCCB |
| ENSG00000113555 | PCDH12 |
| ENSG00000169851 | PCDH7 |
| ENSG00000136099 | PCDH8 |
| ENSG00000126226 | PCID2 |
| ENSG00000100982 | PCIF1 |
| ENSG00000203880 | PCMTD2 |
| ENSG00000106333 | PCOLCE |

| | |
|---|---|
| ENSG00000174788 | PCP2 |
| ENSG00000185813 | PCYT2 |
| ENSG00000150593 | PDCD4 |
| ENSG00000090470 | PDCD7 |
| ENSG00000154678 | PDE1C |
| ENSG00000065989 | PDE4A |
| ENSG00000105650 | PDE4C |
| ENSG00000178104 | PDE4DIP |
| ENSG00000156973 | PDE6D |
| ENSG00000185527 | PDE6G |
| ENSG00000100311 | PDGFB |
| ENSG00000113721 | PDGFRB |
| ENSG00000175087 | PDIK1L |
| ENSG00000004799 | PDK4 |
| ENSG00000131435 | PDLIM4 |
| ENSG00000162493 | PDPN |
| ENSG00000088356 | PDRG1 |
| ENSG00000139515 | PDX1 |
| ENSG00000162734 | PEA15 |
| ENSG00000162517 | PEF1 |
| ENSG00000152684 | PELO |
| ENSG00000179094 | PER1 |
| ENSG00000123349 | PFDN5 |
| ENSG00000158571 | PFKFB1 |
| ENSG00000114268 | PFKFB4 |
| ENSG00000141959 | PFKL |
| ENSG00000067057 | PFKP |
| ENSG00000196570 | PFN3 |
| ENSG00000176732 | PFN4 |
| ENSG00000177614 | PGBD5 |
| ENSG00000184207 | PGP |

| | |
|---|---|
| ENSG00000130517 | PGPEP1 |
| ENSG00000215021 | PHB2 |
| ENSG00000116273 | PHF13 |
| ENSG00000119403 | PHF19 |
| ENSG00000165462 | PHOX2A |
| ENSG00000054148 | PHPT1 |
| ENSG00000155252 | PI4K2A |
| ENSG00000078043 | PIAS2 |
| ENSG00000105229 | PIAS4 |
| ENSG00000073921 | PICALM |
| ENSG00000153823 | PID1 |
| ENSG00000151665 | PIGF |
| ENSG00000100564 | PIGH |
| ENSG00000197563 | PIGN |
| ENSG00000007541 | PIGQ |
| ENSG00000087111 | PIGS |
| ENSG00000101464 | PIGU |
| ENSG00000163964 | PIGX |
| ENSG00000119227 | PIGZ |
| ENSG00000105647 | PIK3R2 |
| ENSG00000117461 | PIK3R3 |
| ENSG00000137193 | PIM1 |
| ENSG00000166908 | PIP4K2C |
| ENSG00000167103 | PIP5KL1 |
| ENSG00000241878 | PISD |
| ENSG00000174238 | PITPNA |
| ENSG00000180957 | PITPNB |
| ENSG00000164093 | PITX2 |
| ENSG00000107859 | PITX3 |
| ENSG00000181191 | PJA1 |
| ENSG00000160447 | PKN3 |

| | |
|---|---|
| ENSG00000160199 | PKNOX1 |
| ENSG00000184363 | PKP3 |
| ENSG00000117215 | PLA2G2D |
| ENSG00000159337 | PLA2G4D |
| ENSG00000145287 | PLAC8 |
| ENSG00000181690 | PLAG1 |
| ENSG00000122861 | PLAU |
| ENSG00000182621 | PLCB1 |
| ENSG00000187091 | PLCD1 |
| ENSG00000161714 | PLCD3 |
| ENSG00000124181 | PLCG1 |
| ENSG00000178209 | PLEC |
| ENSG00000175895 | PLEKHF2 |
| ENSG00000171680 | PLEKHG5 |
| ENSG00000225190 | PLEKHM1 |
| ENSG00000167676 | PLIN4 |
| ENSG00000214456 | PLIN5 |
| ENSG00000166851 | PLK1 |
| ENSG00000102007 | PLP2 |
| ENSG00000187838 | PLSCR3 |
| ENSG00000161381 | PLXDC1 |
| ENSG00000130827 | PLXNA3 |
| ENSG00000164050 | PLXNB1 |
| ENSG00000160783 | PMF1 |
| ENSG00000260238 | PMF1-BGLAP |
| ENSG00000127838 | PNKD |
| ENSG00000176903 | PNMA1 |
| ENSG00000183837 | PNMA3 |
| ENSG00000141744 | PNMT |
| ENSG00000168081 | PNOC |
| ENSG00000180316 | PNPLA1 |

| | |
|---|---|
| ENSG00000177666 | PNPLA2 |
| ENSG00000189266 | PNRC2 |
| ENSG00000174348 | PODN |
| ENSG00000132000 | PODNL1 |
| ENSG00000114631 | PODXL2 |
| ENSG00000186866 | POFUT2 |
| ENSG00000106628 | POLD2 |
| ENSG00000175482 | POLD4 |
| ENSG00000100479 | POLE2 |
| ENSG00000101751 | POLI |
| ENSG00000186184 | POLR1D |
| ENSG00000137054 | POLR1E |
| ENSG00000047315 | POLR2B |
| ENSG00000144231 | POLR2D |
| ENSG00000132963 | POMP |
| ENSG00000172336 | POP7 |
| ENSG00000127948 | POR |
| ENSG00000137709 | POU2F3 |
| ENSG00000151615 | POU4F2 |
| ENSG00000091010 | POU4F3 |
| ENSG00000204531 | POU5F1 |
| ENSG00000184271 | POU6F1 |
| ENSG00000186951 | PPARA |
| ENSG00000084072 | PPIE |
| ENSG00000240344 | PPIL3 |
| ENSG00000131013 | PPIL4 |
| ENSG00000155367 | PPM1J |
| ENSG00000204569 | PPP1R10 |
| ENSG00000088808 | PPP1R13B |
| ENSG00000104881 | PPP1R13L |
| ENSG00000173457 | PPP1R14B |

| | |
|---|---|
| ENSG00000160972 | PPP1R16A |
| ENSG00000135447 | PPP1R1A |
| ENSG00000158528 | PPP1R9A |
| ENSG00000108819 | PPP1R9B |
| ENSG00000074211 | PPP2R2C |
| ENSG00000175470 | PPP2R2D |
| ENSG00000068971 | PPP2R5B |
| ENSG00000120910 | PPP3CC |
| ENSG00000105063 | PPP6R1 |
| ENSG00000108849 | PPY |
| ENSG00000137509 | PRCP |
| ENSG00000138078 | PREPL |
| ENSG00000012211 | PRICKLE3 |
| ENSG00000175785 | PRIMA1 |
| ENSG00000111725 | PRKAB1 |
| ENSG00000072062 | PRKACA |
| ENSG00000181929 | PRKAG1 |
| ENSG00000106617 | PRKAG2 |
| ENSG00000108946 | PRKAR1A |
| ENSG00000005249 | PRKAR2B |
| ENSG00000166501 | PRKCB |
| ENSG00000163932 | PRKCD |
| ENSG00000027075 | PRKCH |
| ENSG00000105287 | PRKD2 |
| ENSG00000185532 | PRKG1 |
| ENSG00000183943 | PRKX |
| ENSG00000250799 | PRODH2 |
| ENSG00000007062 | PROM1 |
| ENSG00000155066 | PROM2 |
| ENSG00000135406 | PRPH |
| ENSG00000167183 | PRR15L |

| | |
|---|---|
| ENSG00000204576 | PRR3 |
| ENSG00000186654 | PRR5 |
| ENSG00000131188 | PRR7 |
| ENSG00000130723 | PRRC2B |
| ENSG00000164099 | PRSS12 |
| ENSG00000005001 | PRSS22 |
| ENSG00000172382 | PRSS27 |
| ENSG00000215148 | PRSS41 |
| ENSG00000196415 | PRTN3 |
| ENSG00000167653 | PSCA |
| ENSG00000146005 | PSD2 |
| ENSG00000143801 | PSEN2 |
| ENSG00000205220 | PSMB10 |
| ENSG00000222028 | PSMB11 |
| ENSG00000013275 | PSMC4 |
| ENSG00000100519 | PSMC6 |
| ENSG00000175166 | PSMD2 |
| ENSG00000095261 | PSMD5 |
| ENSG00000092010 | PSME1 |
| ENSG00000134222 | PSRC1 |
| ENSG00000140368 | PSTPIP1 |
| ENSG00000152229 | PSTPIP2 |
| ENSG00000107317 | PTGDS |
| ENSG00000125384 | PTGER2 |
| ENSG00000050628 | PTGER3 |
| ENSG00000134247 | PTGFRN |
| ENSG00000160013 | PTGIR |
| ENSG00000124212 | PTGIS |
| ENSG00000106853 | PTGR1 |
| ENSG00000184489 | PTP4A3 |
| ENSG00000196396 | PTPN1 |

| | |
|---|---|
| ENSG00000152104 | PTPN14 |
| ENSG00000072135 | PTPN18 |
| ENSG00000127329 | PTPRB |
| ENSG00000213402 | PTPRCAP |
| ENSG00000144724 | PTPRG |
| ENSG00000179950 | PUF60 |
| ENSG00000134644 | PUM1 |
| ENSG00000146676 | PURB |
| ENSG00000110060 | PUS3 |
| ENSG00000130508 | PXDN |
| ENSG00000089159 | PXN |
| ENSG00000183010 | PYCR1 |
| ENSG00000100504 | PYGL |
| ENSG00000129646 | QRICH2 |
| ENSG00000084733 | RAB10 |
| ENSG00000135631 | RAB11FIP5 |
| ENSG00000139832 | RAB20 |
| ENSG00000132698 | RAB25 |
| ENSG00000157869 | RAB28 |
| ENSG00000137502 | RAB30 |
| ENSG00000168461 | RAB31 |
| ENSG00000172794 | RAB37 |
| ENSG00000167994 | RAB3IL1 |
| ENSG00000197562 | RAB40C |
| ENSG00000172780 | RAB43 |
| ENSG00000167578 | RAB4B |
| ENSG00000175582 | RAB6A |
| ENSG00000123570 | RAB9B |
| ENSG00000177548 | RABEP2 |
| ENSG00000136933 | RABEPK |
| ENSG00000100949 | RABGGTA |

| | |
|---|---|
| ENSG00000183155 | RABIF |
| ENSG00000144134 | RABL2A |
| ENSG00000136238 | RAC1 |
| ENSG00000169750 | RAC3 |
| ENSG00000101146 | RAE1 |
| ENSG00000108557 | RAI1 |
| ENSG00000039560 | RAI14 |
| ENSG00000006451 | RALA |
| ENSG00000017797 | RALBP1 |
| ENSG00000125970 | RALY |
| ENSG00000132329 | RAMP1 |
| ENSG00000099901 | RANBP1 |
| ENSG00000031823 | RANBP3 |
| ENSG00000138698 | RAP1GDS1 |
| ENSG00000165917 | RAPSN |
| ENSG00000172819 | RARG |
| ENSG00000155903 | RASA2 |
| ENSG00000105808 | RASA4 |
| ENSG00000108551 | RASD1 |
| ENSG00000172575 | RASGRP1 |
| ENSG00000068831 | RASGRP2 |
| ENSG00000100276 | RASL10A |
| ENSG00000099849 | RASSF7 |
| ENSG00000123094 | RASSF8 |
| ENSG00000139687 | RB1 |
| ENSG00000122257 | RBBP6 |
| ENSG00000171174 | RBKS |
| ENSG00000188739 | RBM34 |
| ENSG00000132819 | RBM38 |
| ENSG00000153250 | RBMS1 |
| ENSG00000144642 | RBMS3 |

| | |
|---|---|
| ENSG00000114115 | RBP1 |
| ENSG00000162444 | RBP7 |
| ENSG00000124232 | RBPJL |
| ENSG00000172348 | RCAN2 |
| ENSG00000117602 | RCAN3 |
| ENSG00000166965 | RCCD1 |
| ENSG00000173653 | RCE1 |
| ENSG00000117906 | RCN2 |
| ENSG00000142552 | RCN3 |
| ENSG00000089902 | RCOR1 |
| ENSG00000080511 | RDH8 |
| ENSG00000122707 | RECK |
| ENSG00000004700 | RECQL |
| ENSG00000165476 | REEP3 |
| ENSG00000168476 | REEP4 |
| ENSG00000115255 | REEP6 |
| ENSG00000173039 | RELA |
| ENSG00000214022 | REPIN1 |
| ENSG00000165731 | RET |
| ENSG00000092871 | RFFL |
| ENSG00000169733 | RFNG |
| ENSG00000168411 | RFWD3 |
| ENSG00000132005 | RFX1 |
| ENSG00000087903 | RFX2 |
| ENSG00000174136 | RGMB |
| ENSG00000182732 | RGS6 |
| ENSG00000182901 | RGS7 |
| ENSG00000108370 | RGS9 |
| ENSG00000007384 | RHBDF1 |
| ENSG00000129667 | RHBDF2 |
| ENSG00000103269 | RHBDL1 |

| | |
|---|---|
| ENSG00000163914 | RHO |
| ENSG00000155366 | RHOC |
| ENSG00000177105 | RHOG |
| ENSG00000119729 | RHOQ |
| ENSG00000140983 | RHOT2 |
| ENSG00000178796 | RIIAD1 |
| ENSG00000167705 | RILP |
| ENSG00000150977 | RILPL2 |
| ENSG00000079841 | RIMS1 |
| ENSG00000117016 | RIMS3 |
| ENSG00000174791 | RIN1 |
| ENSG00000204227 | RING1 |
| ENSG00000104312 | RIPK2 |
| ENSG00000129465 | RIPK3 |
| ENSG00000107018 | RLN1 |
| ENSG00000178966 | RMI1 |
| ENSG00000153561 | RMND5A |
| ENSG00000206150 | RNASE13 |
| ENSG00000258818 | RNASE4 |
| ENSG00000104889 | RNASEH2A |
| ENSG00000219200 | RNASEK |
| ENSG00000172602 | RND1 |
| ENSG00000164068 | RNF123 |
| ENSG00000070423 | RNF126 |
| ENSG00000133135 | RNF128 |
| ENSG00000170881 | RNF139 |
| ENSG00000137393 | RNF144B |
| ENSG00000176641 | RNF152 |
| ENSG00000158717 | RNF166 |
| ENSG00000158286 | RNF207 |
| ENSG00000212864 | RNF208 |

| | |
|---|---|
| ENSG00000099999 | RNF215 |
| ENSG00000233198 | RNF224 |
| ENSG00000092098 | RNF31 |
| ENSG00000204308 | RNF5 |
| ENSG00000176393 | RNPEP |
| ENSG00000142327 | RNPEPL1 |
| ENSG00000185483 | ROR1 |
| ENSG00000106399 | RPA3 |
| ENSG00000103494 | RPGRIP1L |
| ENSG00000089169 | RPH3A |
| ENSG00000197958 | RPL12 |
| ENSG00000142541 | RPL13A |
| ENSG00000105640 | RPL18A |
| ENSG00000131469 | RPL27 |
| ENSG00000108107 | RPL28 |
| ENSG00000156482 | RPL30 |
| ENSG00000136942 | RPL35 |
| ENSG00000140986 | RPL3L |
| ENSG00000148303 | RPL7A |
| ENSG00000177519 | RPRM |
| ENSG00000142534 | RPS11 |
| ENSG00000071242 | RPS6KA2 |
| ENSG00000177189 | RPS6KA3 |
| ENSG00000175634 | RPS6KB2 |
| ENSG00000007376 | RPUSD1 |
| ENSG00000165526 | RPUSD4 |
| ENSG00000083750 | RRAGB |
| ENSG00000133818 | RRAS2 |
| ENSG00000124782 | RREB1 |
| ENSG00000167325 | RRM1 |
| ENSG00000124541 | RRP36 |

| | |
|---|---|
| ENSG00000179041 | RRS1 |
| ENSG00000134321 | RSAD2 |
| ENSG00000104941 | RSPH6A |
| ENSG00000172426 | RSPH9 |
| ENSG00000115310 | RTN4 |
| ENSG00000108309 | RUNDC3A |
| ENSG00000079102 | RUNX1T1 |
| ENSG00000160753 | RUSC1 |
| ENSG00000013392 | RWDD2A |
| ENSG00000163602 | RYBP |
| ENSG00000196218 | RYR1 |
| ENSG00000198626 | RYR2 |
| ENSG00000189171 | S100A13 |
| ENSG00000189334 | S100A14 |
| ENSG00000196420 | S100A5 |
| ENSG00000197956 | S100A6 |
| ENSG00000267534 | S1PR2 |
| ENSG00000125910 | S1PR4 |
| ENSG00000173432 | SAA1 |
| ENSG00000130254 | SAFB2 |
| ENSG00000256463 | SALL3 |
| ENSG00000130590 | SAMD10 |
| ENSG00000167100 | SAMD14 |
| ENSG00000020577 | SAMD4A |
| ENSG00000156671 | SAMD8 |
| ENSG00000123453 | SARDH |
| ENSG00000175467 | SART1 |
| ENSG00000122122 | SASH3 |
| ENSG00000182568 | SATB1 |
| ENSG00000085365 | SCAMP1 |
| ENSG00000073060 | SCARB1 |

| | |
|---|---|
| ENSG00000074660 | SCARF1 |
| ENSG00000105711 | SCN1B |
| ENSG00000007314 | SCN4A |
| ENSG00000111319 | SCNN1A |
| ENSG00000168447 | SCNN1B |
| ENSG00000284194 | SCO2 |
| ENSG00000180900 | SCRIB |
| ENSG00000141295 | SCRN2 |
| ENSG00000261678 | SCRT1 |
| ENSG00000080293 | SCTR |
| ENSG00000175356 | SCUBE2 |
| ENSG00000115884 | SDC1 |
| ENSG00000054282 | SDCCAG8 |
| ENSG00000205138 | SDHAF1 |
| ENSG00000117118 | SDHB |
| ENSG00000069188 | SDK2 |
| ENSG00000170426 | SDR9C7 |
| ENSG00000139410 | SDSL |
| ENSG00000274529 | SEBOX |
| ENSG00000140612 | SEC11A |
| ENSG00000129657 | SEC14L1 |
| ENSG00000100012 | SEC14L3 |
| ENSG00000133488 | SEC14L4 |
| ENSG00000148396 | SEC16A |
| ENSG00000093183 | SEC22C |
| ENSG00000132432 | SEC61G |
| ENSG00000025796 | SEC63 |
| ENSG00000141574 | SECTM1 |
| ENSG00000091490 | SEL1L3 |
| ENSG00000012171 | SEMA3B |
| ENSG00000001617 | SEMA3F |

| | |
|---|---|
| ENSG00000010319 | SEMA3G |
| ENSG00000196189 | SEMA4A |
| ENSG00000167680 | SEMA6B |
| ENSG00000119231 | SENP5 |
| ENSG00000166192 | SENP8 |
| ENSG00000142864 | SERBP1 |
| ENSG00000140264 | SERF2 |
| ENSG00000196136 | SERPINA3 |
| ENSG00000170054 | SERPINA9 |
| ENSG00000206075 | SERPINB5 |
| ENSG00000170542 | SERPINB9 |
| ENSG00000106366 | SERPINE1 |
| ENSG00000132386 | SERPINF1 |
| ENSG00000163536 | SERPINI1 |
| ENSG00000149212 | SESN3 |
| ENSG00000183576 | SETD3 |
| ENSG00000168137 | SETD5 |
| ENSG00000063015 | SEZ6 |
| ENSG00000174938 | SEZ6L2 |
| ENSG00000087365 | SF3B2 |
| ENSG00000175793 | SFN |
| ENSG00000116560 | SFPQ |
| ENSG00000156384 | SFR1 |
| ENSG00000106483 | SFRP4 |
| ENSG00000120057 | SFRP5 |
| ENSG00000198818 | SFT2D1 |
| ENSG00000168484 | SFTPC |
| ENSG00000133661 | SFTPD |
| ENSG00000170624 | SGCD |
| ENSG00000118515 | SGK1 |
| ENSG00000166224 | SGPL1 |

| | |
|---|---|
| ENSG00000160999 | SH2B2 |
| ENSG00000178217 | SH2D4B |
| ENSG00000142669 | SH3BGRL3 |
| ENSG00000100092 | SH3BP1 |
| ENSG00000174705 | SH3PXD2B |
| ENSG00000156463 | SH3RF2 |
| ENSG00000169247 | SH3TC2 |
| ENSG00000035115 | SH3YL1 |
| ENSG00000105251 | SHD |
| ENSG00000164690 | SHH |
| ENSG00000198892 | SHISA4 |
| ENSG00000164054 | SHISA5 |
| ENSG00000188803 | SHISA6 |
| ENSG00000160410 | SHKBP1 |
| ENSG00000072858 | SIDT1 |
| ENSG00000185187 | SIGIRR |
| ENSG00000105366 | SIGLEC8 |
| ENSG00000170145 | SIK2 |
| ENSG00000169375 | SIN3A |
| ENSG00000198053 | SIRPA |
| ENSG00000096717 | SIRT1 |
| ENSG00000124523 | SIRT5 |
| ENSG00000137078 | SIT1 |
| ENSG00000100625 | SIX4 |
| ENSG00000154839 | SKA1 |
| ENSG00000141293 | SKAP1 |
| ENSG00000157933 | SKI |
| ENSG00000163950 | SLBP |
| ENSG00000253598 | SLC10A5 |
| ENSG00000124140 | SLC12A5 |
| ENSG00000007216 | SLC13A2 |

| | |
|---|---|
| ENSG00000158296 | SLC13A3 |
| ENSG00000141485 | SLC13A5 |
| ENSG00000141469 | SLC14A1 |
| ENSG00000110446 | SLC15A3 |
| ENSG00000174326 | SLC16A11 |
| ENSG00000141526 | SLC16A3 |
| ENSG00000101194 | SLC17A9 |
| ENSG00000165646 | SLC18A2 |
| ENSG00000117479 | SLC19A2 |
| ENSG00000135917 | SLC19A3 |
| ENSG00000105281 | SLC1A5 |
| ENSG00000162383 | SLC1A7 |
| ENSG00000144136 | SLC20A1 |
| ENSG00000144671 | SLC22A14 |
| ENSG00000149452 | SLC22A8 |
| ENSG00000074621 | SLC24A1 |
| ENSG00000183048 | SLC25A10 |
| ENSG00000102078 | SLC25A14 |
| ENSG00000102743 | SLC25A15 |
| ENSG00000100372 | SLC25A17 |
| ENSG00000125648 | SLC25A23 |
| ENSG00000144741 | SLC25A26 |
| ENSG00000162461 | SLC25A34 |
| ENSG00000151729 | SLC25A4 |
| ENSG00000181045 | SLC26A11 |
| ENSG00000174502 | SLC26A9 |
| ENSG00000140284 | SLC27A2 |
| ENSG00000156222 | SLC28A1 |
| ENSG00000197506 | SLC28A3 |
| ENSG00000117394 | SLC2A1 |
| ENSG00000160326 | SLC2A6 |

| | |
|---|---|
| ENSG00000136856 | SLC2A8 |
| ENSG00000170385 | SLC30A1 |
| ENSG00000145740 | SLC30A5 |
| ENSG00000014824 | SLC30A9 |
| ENSG00000198569 | SLC34A3 |
| ENSG00000121073 | SLC35B1 |
| ENSG00000116704 | SLC35D1 |
| ENSG00000130958 | SLC35D2 |
| ENSG00000127526 | SLC35E1 |
| ENSG00000151812 | SLC35F4 |
| ENSG00000186334 | SLC36A3 |
| ENSG00000157800 | SLC37A3 |
| ENSG00000169507 | SLC38A11 |
| ENSG00000166558 | SLC38A8 |
| ENSG00000196950 | SLC39A10 |
| ENSG00000133195 | SLC39A11 |
| ENSG00000141873 | SLC39A3 |
| ENSG00000139540 | SLC39A5 |
| ENSG00000112473 | SLC39A7 |
| ENSG00000114544 | SLC41A3 |
| ENSG00000167703 | SLC43A2 |
| ENSG00000070214 | SLC44A1 |
| ENSG00000143036 | SLC44A3 |
| ENSG00000204385 | SLC44A4 |
| ENSG00000211584 | SLC48A1 |
| ENSG00000080493 | SLC4A4 |
| ENSG00000169241 | SLC50A1 |
| ENSG00000100170 | SLC5A1 |
| ENSG00000198743 | SLC5A3 |
| ENSG00000105641 | SLC5A5 |
| ENSG00000138074 | SLC5A6 |

| | |
|---|---|
| ENSG00000117834 | SLC5A9 |
| ENSG00000157103 | SLC6A1 |
| ENSG00000174358 | SLC6A19 |
| ENSG00000130821 | SLC6A8 |
| ENSG00000130876 | SLC7A10 |
| ENSG00000003989 | SLC7A2 |
| ENSG00000165349 | SLC7A3 |
| ENSG00000099960 | SLC7A4 |
| ENSG00000103257 | SLC7A5 |
| ENSG00000092068 | SLC7A8 |
| ENSG00000118160 | SLC8A2 |
| ENSG00000065054 | SLC9A3R2 |
| ENSG00000197818 | SLC9A8 |
| ENSG00000174640 | SLCO2A1 |
| ENSG00000176463 | SLCO3A1 |
| ENSG00000101187 | SLCO4A1 |
| ENSG00000187122 | SLIT1 |
| ENSG00000126233 | SLURP1 |
| ENSG00000141646 | SMAD4 |
| ENSG00000102038 | SMARCA1 |
| ENSG00000080503 | SMARCA2 |
| ENSG00000138375 | SMARCAL1 |
| ENSG00000113810 | SMC4 |
| ENSG00000157106 | SMG1 |
| ENSG00000116698 | SMG7 |
| ENSG00000119953 | SMNDC1 |
| ENSG00000123415 | SMUG1 |
| ENSG00000135632 | SMYD5 |
| ENSG00000124216 | SNAI1 |
| ENSG00000185669 | SNAI3 |
| ENSG00000099940 | SNAP29 |

| | |
|---|---|
| ENSG00000104976 | SNAPC2 |
| ENSG00000165684 | SNAPC4 |
| ENSG00000064692 | SNCAIP |
| ENSG00000197157 | SND1 |
| ENSG00000144028 | SNRNP200 |
| ENSG00000161981 | SNRNP25 |
| ENSG00000184209 | SNRNP35 |
| ENSG00000104852 | SNRNP70 |
| ENSG00000100028 | SNRPD3 |
| ENSG00000172554 | SNTG2 |
| ENSG00000028528 | SNX1 |
| ENSG00000147164 | SNX12 |
| ENSG00000167208 | SNX20 |
| ENSG00000148158 | SNX30 |
| ENSG00000174226 | SNX31 |
| ENSG00000173548 | SNX33 |
| ENSG00000114520 | SNX4 |
| ENSG00000129515 | SNX6 |
| ENSG00000167780 | SOAT2 |
| ENSG00000112320 | SOBP |
| ENSG00000185338 | SOCS1 |
| ENSG00000274211 | SOCS7 |
| ENSG00000120896 | SORBS3 |
| ENSG00000108018 | SORCS1 |
| ENSG00000167941 | SOST |
| ENSG00000187808 | SOWAHD |
| ENSG00000176887 | SOX11 |
| ENSG00000177732 | SOX12 |
| ENSG00000143842 | SOX13 |
| ENSG00000129194 | SOX15 |
| ENSG00000110693 | SOX6 |

| | |
|---|---|
| ENSG00000171056 | SOX7 |
| ENSG00000005513 | SOX8 |
| ENSG00000164871 | SPAG11B |
| ENSG00000144451 | SPAG16 |
| ENSG00000123352 | SPATS2 |
| ENSG00000196141 | SPATS2L |
| ENSG00000124664 | SPDEF |
| ENSG00000090487 | SPG21 |
| ENSG00000197912 | SPG7 |
| ENSG00000063176 | SPHK2 |
| ENSG00000269404 | SPIB |
| ENSG00000229453 | SPINK8 |
| ENSG00000166145 | SPINT1 |
| ENSG00000107742 | SPOCK2 |
| ENSG00000159674 | SPON2 |
| ENSG00000166068 | SPRED1 |
| ENSG00000203772 | SPRN |
| ENSG00000187678 | SPRY4 |
| ENSG00000167778 | SPRYD3 |
| ENSG00000176422 | SPRYD4 |
| ENSG00000162032 | SPSB3 |
| ENSG00000277363 | SRCIN1 |
| ENSG00000277893 | SRD5A2 |
| ENSG00000116649 | SRM |
| ENSG00000139767 | SRRM4 |
| ENSG00000115875 | SRSF7 |
| ENSG00000271303 | SRXN1 |
| ENSG00000008324 | SS18L2 |
| ENSG00000157216 | SSBP3 |
| ENSG00000141298 | SSH2 |
| ENSG00000139874 | SSTR1 |

| | |
|---|---|
| ENSG00000278195 | SSTR3 |
| ENSG00000117155 | SSX2IP |
| ENSG00000008513 | ST3GAL1 |
| ENSG00000157350 | ST3GAL2 |
| ENSG00000070731 | ST6GALNAC2 |
| ENSG00000111728 | ST8SIA1 |
| ENSG00000140557 | ST8SIA2 |
| ENSG00000113532 | ST8SIA4 |
| ENSG00000148488 | ST8SIA6 |
| ENSG00000066923 | STAG3 |
| ENSG00000124356 | STAMBP |
| ENSG00000131748 | STARD3 |
| ENSG00000174448 | STARD6 |
| ENSG00000118804 | STBD1 |
| ENSG00000115107 | STEAP3 |
| ENSG00000118046 | STK11 |
| ENSG00000115694 | STK25 |
| ENSG00000117632 | STMN1 |
| ENSG00000104435 | STMN2 |
| ENSG00000067221 | STOML1 |
| ENSG00000133115 | STOML3 |
| ENSG00000115808 | STRN |
| ENSG00000196792 | STRN3 |
| ENSG00000134910 | STT3A |
| ENSG00000103266 | STUB1 |
| ENSG00000117758 | STX12 |
| ENSG00000103496 | STX4 |
| ENSG00000162236 | STX5 |
| ENSG00000168952 | STXBP6 |
| ENSG00000130540 | SULT4A1 |
| ENSG00000188612 | SUMO2 |

| | |
|---|---|
| ENSG00000184900 | SUMO3 |
| ENSG00000164828 | SUN1 |
| ENSG00000167098 | SUN5 |
| ENSG00000148291 | SURF2 |
| ENSG00000148248 | SURF4 |
| ENSG00000099994 | SUSD2 |
| ENSG00000157303 | SUSD3 |
| ENSG00000165124 | SVEP1 |
| ENSG00000197321 | SVIL |
| ENSG00000175854 | SWI5 |
| ENSG00000169895 | SYAP1 |
| ENSG00000117614 | SYF2 |
| ENSG00000008056 | SYN1 |
| ENSG00000162520 | SYNC |
| ENSG00000135316 | SYNCRIP |
| ENSG00000108639 | SYNGR2 |
| ENSG00000078269 | SYNJ2 |
| ENSG00000019505 | SYT13 |
| ENSG00000143469 | SYT14 |
| ENSG00000139973 | SYT16 |
| ENSG00000143858 | SYT2 |
| ENSG00000213023 | SYT3 |
| ENSG00000129990 | SYT5 |
| ENSG00000011347 | SYT7 |
| ENSG00000149043 | SYT8 |
| ENSG00000164674 | SYTL3 |
| ENSG00000135569 | TAAR5 |
| ENSG00000138162 | TACC2 |
| ENSG00000013810 | TACC3 |
| ENSG00000115353 | TACR1 |
| ENSG00000169836 | TACR3 |

| | |
|---|---|
| ENSG00000276234 | TADA2A |
| ENSG00000273841 | TAF9 |
| ENSG00000158710 | TAGLN2 |
| ENSG00000177156 | TALDO1 |
| ENSG00000168394 | TAP1 |
| ENSG00000231925 | TAPBP |
| ENSG00000139192 | TAPBPL |
| ENSG00000139546 | TARBP2 |
| ENSG00000173662 | TAS1R1 |
| ENSG00000169962 | TAS1R3 |
| ENSG00000102125 | TAZ |
| ENSG00000166220 | TBATA |
| ENSG00000132405 | TBC1D14 |
| ENSG00000121749 | TBC1D15 |
| ENSG00000162065 | TBC1D24 |
| ENSG00000105254 | TBCB |
| ENSG00000167800 | TBX10 |
| ENSG00000112837 | TBX18 |
| ENSG00000135111 | TBX3 |
| ENSG00000121075 | TBX4 |
| ENSG00000149922 | TBX6 |
| ENSG00000006638 | TBXA2R |
| ENSG00000173991 | TCAP |
| ENSG00000176769 | TCERG1L |
| ENSG00000125878 | TCF15 |
| ENSG00000141002 | TCF25 |
| ENSG00000071564 | TCF3 |
| ENSG00000100721 | TCL1A |
| ENSG00000166046 | TCP11L2 |
| ENSG00000188396 | TCTEX1D4 |
| ENSG00000241186 | TDGF1 |

| | |
|---|---|
| ENSG00000083544 | TDRD3 |
| ENSG00000167074 | TEF |
| ENSG00000129566 | TEP1 |
| ENSG00000147601 | TERF1 |
| ENSG00000166848 | TERF2IP |
| ENSG00000136478 | TEX2 |
| ENSG00000091513 | TF |
| ENSG00000087510 | TFAP2C |
| ENSG00000116819 | TFAP2E |
| ENSG00000029639 | TFB1M |
| ENSG00000198176 | TFDP1 |
| ENSG00000112561 | TFEB |
| ENSG00000041988 | THAP3 |
| ENSG00000178726 | THBD |
| ENSG00000169231 | THBS3 |
| ENSG00000051596 | THOC3 |
| ENSG00000100296 | THOC5 |
| ENSG00000131652 | THOC6 |
| ENSG00000172009 | THOP1 |
| ENSG00000090534 | THPO |
| ENSG00000187720 | THSD4 |
| ENSG00000145365 | TIFA |
| ENSG00000173825 | TIGD3 |
| ENSG00000099800 | TIMM13 |
| ENSG00000035862 | TIMP2 |
| ENSG00000100234 | TIMP3 |
| ENSG00000142910 | TINAGL1 |
| ENSG00000137221 | TJAP1 |
| ENSG00000104067 | TJP1 |
| ENSG00000105289 | TJP3 |
| ENSG00000140332 | TLE3 |

| | |
|---|---|
| ENSG00000164342 | TLR3 |
| ENSG00000149809 | TM7SF2 |
| ENSG00000135926 | TMBIM1 |
| ENSG00000133069 | TMCC2 |
| ENSG00000150403 | TMCO3 |
| ENSG00000113119 | TMCO6 |
| ENSG00000144339 | TMEFF2 |
| ENSG00000166292 | TMEM100 |
| ENSG00000181284 | TMEM102 |
| ENSG00000232258 | TMEM114 |
| ENSG00000183160 | TMEM119 |
| ENSG00000188735 | TMEM120B |
| ENSG00000184986 | TMEM121 |
| ENSG00000179178 | TMEM125 |
| ENSG00000171204 | TMEM126B |
| ENSG00000135956 | TMEM127 |
| ENSG00000181291 | TMEM132E |
| ENSG00000172663 | TMEM134 |
| ENSG00000166575 | TMEM135 |
| ENSG00000244187 | TMEM141 |
| ENSG00000096092 | TMEM14A |
| ENSG00000170006 | TMEM154 |
| ENSG00000164180 | TMEM161B |
| ENSG00000127419 | TMEM175 |
| ENSG00000185475 | TMEM179B |
| ENSG00000105518 | TMEM205 |
| ENSG00000150433 | TMEM218 |
| ENSG00000198133 | TMEM229B |
| ENSG00000149582 | TMEM25 |
| ENSG00000176142 | TMEM39A |
| ENSG00000169964 | TMEM42 |

| | |
|---|---|
| ENSG00000121900 | TMEM54 |
| ENSG00000105696 | TMEM59L |
| ENSG00000137842 | TMEM62 |
| ENSG00000164953 | TMEM67 |
| ENSG00000163472 | TMEM79 |
| ENSG00000162460 | TMEM82 |
| ENSG00000167874 | TMEM88 |
| ENSG00000109084 | TMEM97 |
| ENSG00000179104 | TMTC2 |
| ENSG00000164897 | TMUB1 |
| ENSG00000232810 | TNF |
| ENSG00000109079 | TNFAIP1 |
| ENSG00000183578 | TNFAIP8L3 |
| ENSG00000006327 | TNFRSF12A |
| ENSG00000159958 | TNFRSF13C |
| ENSG00000157873 | TNFRSF14 |
| ENSG00000215788 | TNFRSF25 |
| ENSG00000121858 | TNFSF10 |
| ENSG00000120659 | TNFSF11 |
| ENSG00000125735 | TNFSF14 |
| ENSG00000168884 | TNIP2 |
| ENSG00000159173 | TNNI1 |
| ENSG00000130598 | TNNI2 |
| ENSG00000105048 | TNNT1 |
| ENSG00000130595 | TNNT3 |
| ENSG00000064419 | TNPO3 |
| ENSG00000131746 | TNS4 |
| ENSG00000025772 | TOMM34 |
| ENSG00000130204 | TOMM40 |
| ENSG00000214736 | TOMM6 |
| ENSG00000196683 | TOMM7 |

| | |
|---|---|
| ENSG00000136827 | TOR1A |
| ENSG00000160404 | TOR2A |
| ENSG00000103460 | TOX3 |
| ENSG00000067369 | TP53BP1 |
| ENSG00000175274 | TP53I11 |
| ENSG00000146242 | TPBG |
| ENSG00000186815 | TPCN1 |
| ENSG00000111669 | TPI1 |
| ENSG00000159713 | TPPP3 |
| ENSG00000163870 | TPRA1 |
| ENSG00000158109 | TPRG1L |
| ENSG00000172236 | TPSAB1 |
| ENSG00000197253 | TPSB2 |
| ENSG00000116176 | TPSG1 |
| ENSG00000102871 | TRADD |
| ENSG00000135148 | TRAFD1 |
| ENSG00000054116 | TRAPPC3 |
| ENSG00000007255 | TRAPPC6A |
| ENSG00000153339 | TRAPPC8 |
| ENSG00000167632 | TRAPPC9 |
| ENSG00000124731 | TREM1 |
| ENSG00000213689 | TREX1 |
| ENSG00000255690 | TRIL |
| ENSG00000106785 | TRIM14 |
| ENSG00000234127 | TRIM26 |
| ENSG00000137699 | TRIM29 |
| ENSG00000110171 | TRIM3 |
| ENSG00000204599 | TRIM39 |
| ENSG00000132256 | TRIM5 |
| ENSG00000121236 | TRIM6 |
| ENSG00000158022 | TRIM63 |

| | |
|---|---|
| ENSG00000141569 | TRIM65 |
| ENSG00000185880 | TRIM69 |
| ENSG00000100106 | TRIOBP |
| ENSG00000125733 | TRIP10 |
| ENSG00000153827 | TRIP12 |
| ENSG00000087077 | TRIP6 |
| ENSG00000126814 | TRMT5 |
| ENSG00000100416 | TRMU |
| ENSG00000104321 | TRPA1 |
| ENSG00000133107 | TRPC4 |
| ENSG00000100991 | TRPC4AP |
| ENSG00000083067 | TRPM3 |
| ENSG00000130529 | TRPM4 |
| ENSG00000119121 | TRPM6 |
| ENSG00000167723 | TRPV3 |
| ENSG00000127412 | TRPV5 |
| ENSG00000157514 | TSC22D3 |
| ENSG00000166925 | TSC22D4 |
| ENSG00000154743 | TSEN2 |
| ENSG00000123297 | TSFM |
| ENSG00000102904 | TSNAXIP1 |
| ENSG00000140391 | TSPAN3 |
| ENSG00000064201 | TSPAN32 |
| ENSG00000158457 | TSPAN33 |
| ENSG00000100300 | TSPO |
| ENSG00000187189 | TSPYL4 |
| ENSG00000184281 | TSSC4 |
| ENSG00000206203 | TSSK2 |
| ENSG00000162526 | TSSK3 |
| ENSG00000167094 | TTC16 |
| ENSG00000006555 | TTC22 |

| | |
|---|---|
| ENSG00000172425 | TTC36 |
| ENSG00000085831 | TTC39A |
| ENSG00000155158 | TTC39B |
| ENSG00000068724 | TTC7A |
| ENSG00000124120 | TTPAL |
| ENSG00000183785 | TUBA8 |
| ENSG00000178462 | TUBAL3 |
| ENSG00000137267 | TUBB2A |
| ENSG00000137285 | TUBB2B |
| ENSG00000131462 | TUBG1 |
| ENSG00000130640 | TUBGCP2 |
| ENSG00000198680 | TUSC1 |
| ENSG00000233608 | TWIST2 |
| ENSG00000128791 | TWSG1 |
| ENSG00000086712 | TXLNG |
| ENSG00000087301 | TXNDC16 |
| ENSG00000129235 | TXNDC17 |
| ENSG00000239264 | TXNDC5 |
| ENSG00000115514 | TXNDC9 |
| ENSG00000091164 | TXNL1 |
| ENSG00000184470 | TXNRD2 |
| ENSG00000105397 | TYK2 |
| ENSG00000137831 | UACA |
| ENSG00000197355 | UAP1L1 |
| ENSG00000081307 | UBA5 |
| ENSG00000134882 | UBAC2 |
| ENSG00000175063 | UBE2C |
| ENSG00000109332 | UBE2D3 |
| ENSG00000103275 | UBE2I |
| ENSG00000130725 | UBE2M |
| ENSG00000108106 | UBE2S |

| | |
|---|---|
| ENSG00000244687 | UBE2V1 |
| ENSG00000159202 | UBE2Z |
| ENSG00000130939 | UBE4B |
| ENSG00000122042 | UBL3 |
| ENSG00000198258 | UBL5 |
| ENSG00000118900 | UBN1 |
| ENSG00000160803 | UBQLN4 |
| ENSG00000159459 | UBR1 |
| ENSG00000108312 | UBTF |
| ENSG00000173960 | UBXN2A |
| ENSG00000167671 | UBXN6 |
| ENSG00000104691 | UBXN8 |
| ENSG00000118939 | UCHL3 |
| ENSG00000116750 | UCHL5 |
| ENSG00000109424 | UCP1 |
| ENSG00000176125 | UFSP1 |
| ENSG00000109103 | UNC119 |
| ENSG00000175970 | UNC119B |
| ENSG00000130477 | UNC13A |
| ENSG00000115446 | UNC50 |
| ENSG00000113763 | UNC5A |
| ENSG00000124602 | UNC5CL |
| ENSG00000156687 | UNC5D |
| ENSG00000110057 | UNC93B1 |
| ENSG00000059145 | UNKL |
| ENSG00000005007 | UPF1 |
| ENSG00000105668 | UPK1A |
| ENSG00000100373 | UPK3A |
| ENSG00000130307 | USHBP1 |
| ENSG00000184979 | USP18 |
| ENSG00000273820 | USP27X |

| | |
|---|---|
| ENSG00000187555 | USP7 |
| ENSG00000011260 | UTP18 |
| ENSG00000103043 | VAC14 |
| ENSG00000162738 | VANGL2 |
| ENSG00000160293 | VAV2 |
| ENSG00000148704 | VAX1 |
| ENSG00000035403 | VCL |
| ENSG00000165637 | VDAC2 |
| ENSG00000111424 | VDR |
| ENSG00000150630 | VEGFC |
| ENSG00000197415 | VEPH1 |
| ENSG00000128564 | VGF |
| ENSG00000170162 | VGLL2 |
| ENSG00000136059 | VILL |
| ENSG00000106018 | VIPR2 |
| ENSG00000196715 | VKORC1L1 |
| ENSG00000128218 | VPREB3 |
| ENSG00000215305 | VPS16 |
| ENSG00000139719 | VPS33A |
| ENSG00000184056 | VPS33B |
| ENSG00000155975 | VPS37A |
| ENSG00000166887 | VPS39 |
| ENSG00000223501 | VPS52 |
| ENSG00000019102 | VSIG2 |
| ENSG00000134258 | VTCN1 |
| ENSG00000100568 | VTI1B |
| ENSG00000109072 | VTN |
| ENSG00000095787 | WAC |
| ENSG00000239779 | WBP1 |
| ENSG00000120688 | WBP4 |
| ENSG00000163625 | WDFY3 |

| | |
|---|---|
| ENSG00000198554 | WDHD1 |
| ENSG00000071127 | WDR1 |
| ENSG00000120008 | WDR11 |
| ENSG00000138442 | WDR12 |
| ENSG00000101940 | WDR13 |
| ENSG00000127580 | WDR24 |
| ENSG00000162923 | WDR26 |
| ENSG00000160193 | WDR4 |
| ENSG00000103091 | WDR59 |
| ENSG00000140395 | WDR61 |
| ENSG00000133316 | WDR74 |
| ENSG00000105875 | WDR91 |
| ENSG00000166483 | WEE1 |
| ENSG00000103175 | WFDC1 |
| ENSG00000156076 | WIF1 |
| ENSG00000070540 | WIPI1 |
| ENSG00000169884 | WNT10B |
| ENSG00000085741 | WNT11 |
| ENSG00000134245 | WNT2B |
| ENSG00000108379 | WNT3 |
| ENSG00000114251 | WNT5A |
| ENSG00000115596 | WNT6 |
| ENSG00000154764 | WNT7A |
| ENSG00000188064 | WNT7B |
| ENSG00000176871 | WSB2 |
| ENSG00000142279 | WTIP |
| ENSG00000113645 | WWC1 |
| ENSG00000151718 | WWC2 |
| ENSG00000198373 | WWP2 |
| ENSG00000076924 | XAB2 |
| ENSG00000047597 | XK |

| | |
|---|---|
| ENSG00000206579 | XKR4 |
| ENSG00000221947 | XKR9 |
| ENSG00000122121 | XPNPEP2 |
| ENSG00000082898 | XPO1 |
| ENSG00000143324 | XPR1 |
| ENSG00000127337 | YEATS4 |
| ENSG00000174851 | YIF1A |
| ENSG00000167645 | YIF1B |
| ENSG00000058799 | YIPF1 |
| ENSG00000130733 | YIPF2 |
| ENSG00000119820 | YIPF4 |
| ENSG00000100027 | YPEL1 |
| ENSG00000090238 | YPEL3 |
| ENSG00000196449 | YRDC |
| ENSG00000170027 | YWHAG |
| ENSG00000164924 | YWHAZ |
| ENSG00000182223 | ZAR1 |
| ENSG00000124256 | ZBP1 |
| ENSG00000109906 | ZBTB16 |
| ENSG00000236104 | ZBTB22 |
| ENSG00000011590 | ZBTB32 |
| ENSG00000166860 | ZBTB39 |
| ENSG00000204859 | ZBTB48 |
| ENSG00000168795 | ZBTB5 |
| ENSG00000178951 | ZBTB7A |
| ENSG00000160685 | ZBTB7B |
| ENSG00000213588 | ZBTB9 |
| ENSG00000135482 | ZC3H10 |
| ENSG00000163874 | ZC3H12A |
| ENSG00000130749 | ZC3H4 |
| ENSG00000144161 | ZC3H8 |

| | |
|---|---|
| ENSG00000078487 | ZCWPW1 |
| ENSG00000175048 | ZDHHC14 |
| ENSG00000104219 | ZDHHC2 |
| ENSG00000099904 | ZDHHC8 |
| ENSG00000188706 | ZDHHC9 |
| ENSG00000158552 | ZFAND2B |
| ENSG00000156639 | ZFAND3 |
| ENSG00000142065 | ZFP14 |
| ENSG00000152518 | ZFP36L2 |
| ENSG00000136866 | ZFP37 |
| ENSG00000187815 | ZFP69 |
| ENSG00000181007 | ZFP82 |
| ENSG00000179588 | ZFPM1 |
| ENSG00000056097 | ZFR |
| ENSG00000166140 | ZFYVE19 |
| ENSG00000178764 | ZHX2 |
| ENSG00000172667 | ZMAT3 |
| ENSG00000122515 | ZMIZ2 |
| ENSG00000004838 | ZMYND10 |
| ENSG00000066185 | ZMYND12 |
| ENSG00000101040 | ZMYND8 |
| ENSG00000175787 | ZNF169 |
| ENSG00000275111 | ZNF2 |
| ENSG00000165512 | ZNF22 |
| ENSG00000159917 | ZNF235 |
| ENSG00000172466 | ZNF24 |
| ENSG00000254004 | ZNF260 |
| ENSG00000006194 | ZNF263 |
| ENSG00000063587 | ZNF275 |
| ENSG00000158805 | ZNF276 |
| ENSG00000166526 | ZNF3 |

| | |
|---|---|
| ENSG00000166188 | ZNF319 |
| ENSG00000181894 | ZNF329 |
| ENSG00000198816 | ZNF358 |
| ENSG00000178175 | ZNF366 |
| ENSG00000161642 | ZNF385A |
| ENSG00000186918 | ZNF395 |
| ENSG00000197050 | ZNF420 |
| ENSG00000102935 | ZNF423 |
| ENSG00000112200 | ZNF451 |
| ENSG00000196263 | ZNF471 |
| ENSG00000265763 | ZNF488 |
| ENSG00000165655 | ZNF503 |
| ENSG00000101493 | ZNF516 |
| ENSG00000198795 | ZNF521 |
| ENSG00000167625 | ZNF526 |
| ENSG00000187187 | ZNF546 |
| ENSG00000186017 | ZNF566 |
| ENSG00000105732 | ZNF574 |
| ENSG00000176472 | ZNF575 |
| ENSG00000198440 | ZNF583 |
| ENSG00000166716 | ZNF592 |
| ENSG00000272602 | ZNF595 |
| ENSG00000167962 | ZNF598 |
| ENSG00000197483 | ZNF628 |
| ENSG00000167528 | ZNF641 |
| ENSG00000122482 | ZNF644 |
| ENSG00000143373 | ZNF687 |
| ENSG00000169957 | ZNF768 |
| ENSG00000197782 | ZNF780A |
| ENSG00000278129 | ZNF8 |
| ENSG00000106400 | ZNHIT1 |

| | |
|---|---|
| ENSG00000121903 | ZSCAN20 |
| ENSG00000182318 | ZSCAN22 |
| ENSG00000162378 | ZYG11B |

**Table S5. SRY target genes differentially expressed in HGG samples presenting higher versus lower SRY expression.**

| Up-regulated SRY target genes | | | | Down-regulated SRY target genes | | | |
|---|---|---|---|---|---|---|---|
| **gene.id** | **gene.symbol** | **log2FC** | **FDR** | **gene.id** | **gene.symbol** | **log2FC** | **FDR** |
| ENSG00000173432.12 | SAA1 | 4.32425 | 3.63E-21 | ENSG00000089116.4 | LHX5 | -2.41216 | 3.24E-11 |
| ENSG00000133115.12 | STOML3 | 3.63732 | 8.01E-21 | ENSG00000111249.14 | CUX2 | -2.23459 | 8.20E-19 |
| ENSG00000164093.17 | PITX2 | 3.44177 | 3.25E-16 | ENSG00000173157.17 | ADAMTS20 | -2.19144 | 5.77E-13 |
| ENSG00000142173.16 | COL6A2 | 2.56552 | 4.72E-25 | ENSG00000129152.4 | MYOD1 | -2.04494 | 8.53E-06 |
| ENSG00000162344.4 | FGF19 | 2.47689 | 1.01E-05 | ENSG00000261678.3 | SCRT1 | -1.87773 | 9.29E-15 |
| ENSG00000168447.11 | SCNN1B | 2.38752 | 4.69E-15 | ENSG00000198910.14 | L1CAM | -1.82938 | 2.19E-13 |
| ENSG00000164400.6 | CSF2 | 2.33977 | 3.39E-05 | ENSG00000139874.6 | SSTR1 | -1.77049 | 1.94E-10 |
| ENSG00000055957.11 | ITIH1 | 2.28125 | 1.70E-17 | ENSG00000130829.18 | DUSP9 | -1.72524 | 1.11E-13 |
| ENSG00000186191.8 | BPIFB4 | 2.20707 | 1.45E-09 | ENSG00000144278.15 | GALNT13 | -1.71633 | 1.02E-14 |
| ENSG00000159217.10 | IGF2BP1 | 2.17794 | 5.88E-10 | ENSG00000125845.7 | BMP2 | -1.68166 | 1.56E-19 |
| ENSG00000172482.5 | AGXT | 2.15403 | 8.99E-09 | ENSG00000166206.15 | GABRB3 | -1.64565 | 2.62E-14 |
| ENSG00000125735.11 | TNFSF14 | 2.07941 | 6.87E-18 | ENSG00000196092.14 | PAX5 | -1.62783 | 1.54E-08 |
| ENSG00000180613.11 | GSX2 | 2.05040 | 1.31E-11 | ENSG00000187486.5 | KCNJ11 | -1.59834 | 1.34E-13 |
| ENSG00000130513.6 | GDF15 | 2.03867 | 8.51E-15 | ENSG00000188064.10 | WNT7B | -1.58947 | 1.65E-11 |
| ENSG00000111424.12 | VDR | 2.02660 | 1.11E-21 | ENSG00000145451.13 | GLRA3 | -1.58300 | 1.29E-07 |
| ENSG00000170454.6 | KRT75 | 2.00140 | 4.39E-08 | ENSG00000106018.14 | VIPR2 | -1.50719 | 5.76E-09 |
| ENSG00000108821.14 | COL1A1 | 1.99015 | 1.24E-12 | ENSG00000092054.13 | MYH7 | -1.50042 | 1.15E-13 |
| ENSG00000132000.13 | PODNL1 | 1.98743 | 1.30E-18 | ENSG00000143858.12 | SYT2 | -1.48863 | 6.23E-10 |
| ENSG00000131435.13 | PDLIM4 | 1.96120 | 2.74E-15 | ENSG00000100604.13 | CHGA | -1.48787 | 5.36E-09 |
| ENSG00000130635.16 | COL5A1 | 1.92142 | 6.42E-14 | ENSG00000166006.14 | KCNC2 | -1.47307 | 2.35E-07 |
| ENSG00000072694.21 | FCGR2B | 1.91926 | 1.01E-13 | ENSG00000196475.6 | GK2 | -1.47259 | 5.69E-05 |
| ENSG00000182742.6 | HOXB4 | 1.91117 | 7.90E-08 | ENSG00000148386.9 | LCN9 | -1.46618 | 1.89E-05 |
| ENSG00000007314.12 | SCN4A | 1.89578 | 6.19E-16 | ENSG00000154118.13 | JPH3 | -1.44596 | 9.78E-11 |
| ENSG00000069482.7 | GAL | 1.89385 | 3.09E-12 | ENSG00000152672.8 | CLEC4F | -1.44198 | 1.92E-10 |
| ENSG00000114115.10 | RBP1 | 1.88222 | 1.63E-16 | ENSG00000113763.12 | UNC5A | -1.43506 | 8.28E-15 |
| ENSG00000184350.11 | MRGPRE | 1.88165 | 1.75E-11 | ENSG00000151025.11 | GPR158 | -1.41448 | 4.32E-18 |
| ENSG00000179023.8 | KLHDC7A | 1.87936 | 4.62E-08 | ENSG00000167654.18 | ATCAY | -1.39233 | 6.02E-13 |
| ENSG00000128710.6 | HOXD10 | 1.86935 | 1.11E-06 | ENSG00000165370.3 | GPR101 | -1.38557 | 6.89E-06 |

| | | | | | | | |
|---|---|---|---|---|---|---|---|
| ENSG00000073734.10 | ABCB11 | 1.84746 | 2.53E-12 | ENSG00000105251.11 | SHD | -1.37973 | 3.45E-12 |
| ENSG00000144891.18 | AGTR1 | 1.83369 | 4.17E-15 | ENSG00000124194.17 | GDAP1L1 | -1.35361 | 3.33E-15 |
| ENSG00000128342.5 | LIF | 1.82165 | 2.86E-11 | ENSG00000136750.13 | GAD2 | -1.35131 | 1.50E-06 |
| ENSG00000142227.11 | EMP3 | 1.80453 | 5.75E-14 | ENSG00000168621.15 | GDNF | -1.34882 | 4.08E-11 |
| ENSG00000115107.20 | STEAP3 | 1.79207 | 1.39E-21 | ENSG00000112164.6 | GLP1R | -1.33823 | 1.79E-05 |
| ENSG00000129038.16 | LOXL1 | 1.76591 | 6.44E-15 | ENSG00000092051.17 | JPH4 | -1.33259 | 9.29E-15 |
| ENSG00000080511.5 | RDH8 | 1.75890 | 9.57E-08 | ENSG00000170579.17 | DLGAP1 | -1.32822 | 7.05E-17 |
| ENSG00000106004.5 | HOXA5 | 1.74024 | 5.95E-06 | ENSG00000005513.10 | SOX8 | -1.31692 | 4.82E-15 |
| ENSG00000109182.12 | CWH43 | 1.73374 | 2.09E-05 | ENSG00000152954.12 | NRSN1 | -1.28815 | 3.37E-12 |
| ENSG00000167941.3 | SOST | 1.73370 | 1.43E-06 | ENSG00000183570.16 | PCBP3 | -1.28730 | 3.16E-14 |
| ENSG00000170162.14 | VGLL2 | 1.72571 | 1.88E-05 | ENSG00000103460.17 | TOX3 | -1.28615 | 1.87E-13 |
| ENSG00000124731.13 | TREM1 | 1.71619 | 2.29E-09 | ENSG00000187122.17 | SLIT1 | -1.27871 | 1.91E-12 |
| ENSG00000159713.11 | TPPP3 | 1.69436 | 2.42E-16 | ENSG00000141668.10 | CBLN2 | -1.26319 | 1.70E-06 |
| ENSG00000253293.5 | HOXA10 | 1.68587 | 1.85E-06 | ENSG00000172346.15 | CSDC2 | -1.26014 | 1.29E-08 |
| ENSG00000016402.13 | IL20RA | 1.67235 | 1.98E-12 | ENSG00000029534.21 | ANK1 | -1.25947 | 4.46E-13 |
| ENSG00000159184.8 | HOXB13 | 1.66512 | 1.67E-04 | ENSG00000008300.17 | CELSR3 | -1.25849 | 2.60E-15 |
| ENSG00000112319.19 | EYA4 | 1.63957 | 8.35E-09 | ENSG00000136297.15 | MMD2 | -1.25028 | 3.34E-10 |
| ENSG00000129654.8 | FOXJ1 | 1.63759 | 9.63E-14 | ENSG00000166292.12 | TMEM100 | -1.24161 | 2.57E-11 |
| ENSG00000115507.10 | OTX1 | 1.62282 | 1.58E-11 | ENSG00000115353.11 | TACR1 | -1.23867 | 3.20E-07 |
| ENSG00000140287.11 | HDC | 1.59201 | 4.53E-07 | ENSG00000165379.13 | LRFN5 | -1.23640 | 1.97E-07 |
| ENSG00000172236.18 | TPSAB1 | 1.58373 | 1.36E-05 | ENSG00000130477.15 | UNC13A | -1.23078 | 1.54E-12 |
| ENSG00000162493.16 | PDPN | 1.56786 | 1.88E-09 | ENSG00000139767.10 | SRRM4 | -1.22447 | 3.79E-07 |
| ENSG00000137709.10 | POU2F3 | 1.55229 | 5.54E-08 | ENSG00000197959.14 | DNM3 | -1.22412 | 1.14E-16 |
| ENSG00000116014.10 | KISS1R | 1.53709 | 3.95E-06 | ENSG00000166159.11 | LRTM2 | -1.21904 | 1.04E-05 |
| ENSG00000158710.15 | TAGLN2 | 1.52507 | 3.51E-18 | ENSG00000148408.13 | CACNA1B | -1.21504 | 1.07E-06 |
| ENSG00000156966.7 | B3GNT7 | 1.51611 | 2.00E-15 | ENSG00000159409.15 | CELF3 | -1.21235 | 2.24E-10 |
| ENSG00000205076.4 | LGALS7 | 1.51536 | 1.77E-03 | ENSG00000182621.18 | PLCB1 | -1.20889 | 5.99E-17 |
| ENSG00000174697.5 | LEP | 1.51482 | 9.73E-07 | ENSG00000171587.15 | DSCAM | -1.20032 | 2.24E-15 |
| ENSG00000197766.8 | CFD | 1.51421 | 2.37E-16 | ENSG00000129682.16 | FGF13 | -1.17799 | 8.63E-09 |
| ENSG00000102007.11 | PLP2 | 1.50668 | 1.25E-14 | ENSG00000137460.9 | FHDC1 | -1.17718 | 2.61E-13 |
| ENSG00000173917.10 | HOXB2 | 1.49977 | 9.07E-07 | ENSG00000153823.19 | PID1 | -1.17645 | 4.34E-19 |

| | | | | | | | |
|---|---|---|---|---|---|---|---|
| ENSG00000130876.11 | SLC7A10 | 1.49171 | 5.68E-07 | ENSG00000180720.7 | CHRM4 | -1.16018 | 4.70E-11 |
| ENSG00000232258.6 | TMEM114 | 1.48269 | 6.72E-06 | ENSG00000206075.14 | SERPINB5 | -1.15691 | 1.04E-02 |
| ENSG00000159674.12 | SPON2 | 1.47182 | 3.14E-10 | ENSG00000145808.10 | ADAMTS19 | -1.15050 | 5.05E-07 |
| ENSG00000060982.15 | BCAT1 | 1.46940 | 1.12E-13 | ENSG00000144119.4 | C1QL2 | -1.13210 | 1.68E-06 |
| ENSG00000121310.17 | ECHDC2 | 1.45553 | 1.80E-19 | ENSG00000128683.14 | GAD1 | -1.12953 | 3.99E-10 |
| ENSG00000015413.9 | DPEP1 | 1.45216 | 2.76E-08 | ENSG00000175874.10 | CREG2 | -1.12513 | 5.72E-05 |
| ENSG00000106006.6 | HOXA6 | 1.43097 | 7.40E-04 | ENSG00000102385.12 | DRP2 | -1.12027 | 4.64E-17 |
| ENSG00000106366.9 | SERPINE1 | 1.43009 | 1.21E-07 | ENSG00000100276.10 | RASL10A | -1.11996 | 1.11E-08 |
| ENSG00000178934.5 | LGALS7B | 1.42748 | 2.19E-04 | ENSG00000186868.16 | MAPT | -1.11174 | 7.29E-20 |
| ENSG00000087510.7 | TFAP2C | 1.42504 | 2.62E-09 | ENSG00000176749.9 | CDK5R1 | -1.11041 | 3.00E-15 |
| ENSG00000118231.5 | CRYGD | 1.42375 | 6.55E-05 | ENSG00000081052.13 | COL4A4 | -1.09753 | 2.16E-08 |
| ENSG00000004838.14 | ZMYND10 | 1.41997 | 8.58E-11 | ENSG00000175497.17 | DPP10 | -1.09691 | 1.75E-07 |
| ENSG00000213892.12 | CEACAM16 | 1.41975 | 1.63E-04 | ENSG00000172461.11 | FUT9 | -1.09639 | 6.88E-10 |
| ENSG00000156222.12 | SLC28A1 | 1.41321 | 4.86E-13 | ENSG00000135973.3 | GPR45 | -1.09599 | 3.36E-11 |
| ENSG00000174807.4 | CD248 | 1.40639 | 6.88E-11 | ENSG00000110693.18 | SOX6 | -1.09514 | 3.64E-17 |
| ENSG00000106333.13 | PCOLCE | 1.38243 | 3.78E-14 | ENSG00000183166.11 | CALN1 | -1.09382 | 5.46E-05 |
| ENSG00000120659.15 | TNFSF11 | 1.37774 | 9.63E-05 | ENSG00000139973.16 | SYT16 | -1.08769 | 1.71E-07 |
| ENSG00000107159.13 | CA9 | 1.37285 | 1.01E-05 | ENSG00000175161.14 | CADM2 | -1.08446 | 1.86E-12 |
| ENSG00000106436.7 | MYL10 | 1.36730 | 9.09E-05 | ENSG00000108309.14 | RUNDC3A | -1.07059 | 8.60E-12 |
| ENSG00000073792.16 | IGF2BP2 | 1.36349 | 2.40E-07 | ENSG00000050628.20 | PTGER3 | -1.06732 | 3.08E-05 |
| ENSG00000196136.18 | SERPINA3 | 1.35950 | 1.62E-10 | ENSG00000185737.13 | NRG3 | -1.06132 | 2.28E-08 |
| ENSG00000185338.6 | SOCS1 | 1.35381 | 2.96E-13 | ENSG00000182568.17 | SATB1 | -1.05576 | 2.41E-19 |
| ENSG00000188641.14 | DPYD | 1.35280 | 3.47E-15 | ENSG00000166342.19 | NETO1 | -1.05530 | 1.75E-06 |
| ENSG00000130592.17 | LSP1 | 1.35246 | 1.93E-13 | ENSG00000168993.15 | CPLX1 | -1.05264 | 1.38E-08 |
| ENSG00000241644.2 | INMT | 1.35056 | 2.23E-09 | ENSG00000137478.15 | FCHSD2 | -1.05112 | 1.63E-27 |
| ENSG00000120057.5 | SFRP5 | 1.34427 | 5.20E-06 | ENSG00000166501.14 | PRKCB | -1.05111 | 3.77E-08 |
| ENSG00000172638.13 | EFEMP2 | 1.34411 | 4.06E-15 | ENSG00000134569.10 | LRP4 | -1.04811 | 6.97E-13 |
| ENSG00000170178.7 | HOXD12 | 1.33686 | 6.82E-04 | ENSG00000110987.9 | BCL7A | -1.04465 | 4.64E-18 |
| ENSG00000182472.9 | CAPN12 | 1.33654 | 1.26E-12 | ENSG00000162374.18 | ELAVL4 | -1.04042 | 1.03E-08 |
| ENSG00000178814.17 | OPLAH | 1.33558 | 1.35E-19 | ENSG00000168772.11 | CXXC4 | -1.03631 | 1.90E-13 |
| ENSG00000106153.13 | CHCHD2 | 1.33497 | 7.84E-18 | ENSG00000187720.14 | THSD4 | -1.03542 | 6.33E-08 |

| | | | | | | | |
|---|---|---|---|---|---|---|---|
| ENSG00000099953.10 | MMP11 | 1.32860 | 2.65E-11 | ENSG00000109906.14 | ZBTB16 | -1.03374 | 1.67E-09 |
| ENSG00000188396.4 | TCTEX1D4 | 1.32619 | 2.68E-10 | ENSG00000130540.14 | SULT4A1 | -1.02962 | 1.33E-04 |
| ENSG00000137078.9 | SIT1 | 1.32510 | 4.77E-10 | ENSG00000164398.15 | ACSL6 | -1.02896 | 6.58E-10 |
| ENSG00000172426.16 | RSPH9 | 1.31637 | 4.40E-14 | ENSG00000080644.16 | CHRNA3 | -1.02718 | 4.21E-05 |
| ENSG00000183798.5 | EMILIN3 | 1.31608 | 8.92E-06 | ENSG00000173848.19 | NET1 | -1.02693 | 7.76E-12 |
| ENSG00000146242.9 | TPBG | 1.30602 | 3.59E-10 | ENSG00000111262.6 | KCNA1 | -1.02436 | 1.72E-05 |
| ENSG00000121743.4 | GJA3 | 1.30357 | 4.59E-06 | ENSG00000079841.18 | RIMS1 | -1.02351 | 8.56E-07 |
| ENSG00000180818.5 | HOXC10 | 1.30211 | 4.97E-03 | ENSG00000198626.17 | RYR2 | -1.02076 | 2.10E-04 |
| ENSG00000205403.13 | CFI | 1.30003 | 3.03E-11 | ENSG00000101076.17 | HNF4A | -1.01638 | 2.06E-03 |
| ENSG00000147065.17 | MSN | 1.29308 | 3.46E-21 | ENSG00000135625.8 | EGR4 | -1.00945 | 6.20E-04 |
| ENSG00000111058.8 | ACSS3 | 1.28270 | 4.40E-13 | ENSG00000187922.14 | LCN10 | -1.00617 | 8.07E-04 |
| ENSG00000157227.13 | MMP14 | 1.28067 | 2.52E-14 | ENSG00000174576.10 | NPAS4 | -1.00525 | 2.76E-04 |
| ENSG00000139610.2 | CELA1 | 1.26443 | 2.70E-08 | ENSG00000124140.14 | SLC12A5 | -1.00500 | 5.67E-05 |
| ENSG00000135926.15 | TMBIM1 | 1.26408 | 3.47E-24 | ENSG00000176769.9 | TCERG1L | -1.00118 | 1.68E-04 |
| ENSG00000197587.11 | DMBX1 | 1.26125 | 3.71E-04 | ENSG00000019505.8 | SYT13 | -1.00103 | 1.37E-04 |
| ENSG00000104951.16 | IL4I1 | 1.25809 | 7.53E-11 | | | | |
| ENSG00000182223.7 | ZAR1 | 1.25345 | 2.32E-05 | | | | |
| ENSG00000197405.8 | C5AR1 | 1.25148 | 2.31E-11 | | | | |
| ENSG00000006327.14 | TNFRSF12A | 1.25058 | 1.33E-08 | | | | |
| ENSG00000162998.5 | FRZB | 1.23839 | 7.09E-13 | | | | |
| ENSG00000106927.12 | AMBP | 1.23269 | 1.39E-06 | | | | |
| ENSG00000108849.8 | PPY | 1.22996 | 2.16E-04 | | | | |
| ENSG00000042493.16 | CAPG | 1.22898 | 5.23E-15 | | | | |
| ENSG00000141526.17 | SLC16A3 | 1.22745 | 4.42E-13 | | | | |
| ENSG00000104783.14 | KCNN4 | 1.22744 | 7.89E-10 | | | | |
| ENSG00000120094.9 | HOXB1 | 1.22367 | 2.09E-02 | | | | |
| ENSG00000100739.11 | BDKRB1 | 1.22303 | 1.99E-07 | | | | |
| ENSG00000134871.19 | COL4A2 | 1.22024 | 2.30E-09 | | | | |
| ENSG00000128709.13 | HOXD9 | 1.21939 | 3.34E-04 | | | | |
| ENSG00000186510.12 | CLCNKA | 1.21878 | 1.81E-08 | | | | |
| ENSG00000117122.14 | MFAP2 | 1.21686 | 2.56E-06 | | | | |

| | | | |
|---|---|---|---|
| ENSG00000124216.4 | SNAI1 | 1.21679 | 1.59E-09 |
| ENSG00000110448.11 | CD5 | 1.21484 | 5.38E-13 |
| ENSG00000135407.10 | AVIL | 1.21331 | 1.46E-13 |
| ENSG00000139549.4 | DHH | 1.21197 | 1.35E-10 |
| ENSG00000187678.10 | SPRY4 | 1.21167 | 6.40E-09 |
| ENSG00000012211.13 | PRICKLE3 | 1.20582 | 1.36E-17 |
| ENSG00000121236.21 | TRIM6 | 1.20370 | 1.37E-14 |
| ENSG00000163827.13 | LRRC2 | 1.20190 | 1.56E-08 |
| ENSG00000099985.4 | OSM | 1.19380 | 5.47E-08 |
| ENSG00000105664.11 | COMP | 1.18715 | 1.98E-07 |
| ENSG00000258818.4 | RNASE4 | 1.18701 | 2.71E-09 |
| ENSG00000175505.11 | CLCF1 | 1.18022 | 2.20E-07 |
| ENSG00000112539.15 | C6orf118 | 1.17864 | 5.38E-10 |
| ENSG00000108342.13 | CSF3 | 1.17857 | 7.16E-04 |
| ENSG00000064300.9 | NGFR | 1.17742 | 1.95E-10 |
| ENSG00000085831.15 | TTC39A | 1.17670 | 2.15E-12 |
| ENSG00000106483.12 | SFRP4 | 1.17532 | 9.30E-10 |
| ENSG00000134013.16 | LOXL2 | 1.17232 | 4.43E-10 |
| ENSG00000258083.2 | OR9A4 | 1.17098 | 5.21E-03 |
| ENSG00000091490.11 | SEL1L3 | 1.15462 | 2.56E-09 |
| ENSG00000196420.7 | S100A5 | 1.15127 | 1.44E-06 |
| ENSG00000140839.11 | CLEC18B | 1.14719 | 1.29E-08 |
| ENSG00000132432.14 | SEC61G | 1.14498 | 1.07E-04 |
| ENSG00000115884.11 | SDC1 | 1.14412 | 3.01E-08 |
| ENSG00000188624.3 | IGFL3 | 1.14402 | 1.74E-07 |
| ENSG00000197273.4 | GUCA2A | 1.14247 | 5.75E-03 |
| ENSG00000221986.7 | MYBPHL | 1.13556 | 1.94E-12 |
| ENSG00000170006.12 | TMEM154 | 1.13546 | 2.29E-13 |
| ENSG00000100504.17 | PYGL | 1.13454 | 6.75E-15 |
| ENSG00000157873.17 | TNFRSF14 | 1.13169 | 1.52E-19 |
| ENSG00000156535.15 | CD109 | 1.12761 | 3.07E-14 |

| | | | |
|---|---|---|---|
| ENSG00000162458.13 | FBLIM1 | 1.12421 | 2.65E-08 |
| ENSG00000104321.11 | TRPA1 | 1.12003 | 1.71E-04 |
| ENSG00000115414.21 | FN1 | 1.11826 | 1.10E-10 |
| ENSG00000139515.6 | PDX1 | 1.11782 | 2.19E-02 |
| ENSG00000173641.18 | HSPB7 | 1.11684 | 4.88E-08 |
| ENSG00000099994.11 | SUSD2 | 1.11579 | 1.21E-12 |
| ENSG00000178401.16 | DNAJC22 | 1.10872 | 1.18E-13 |
| ENSG00000162520.15 | SYNC | 1.10248 | 1.53E-14 |
| ENSG00000189171.14 | S100A13 | 1.10247 | 4.53E-10 |
| ENSG00000087077.14 | TRIP6 | 1.10005 | 2.42E-18 |
| ENSG00000170608.3 | FOXA3 | 1.09930 | 2.51E-05 |
| ENSG00000233608.4 | TWIST2 | 1.09823 | 5.76E-08 |
| ENSG00000101608.13 | MYL12A | 1.09463 | 2.31E-16 |
| ENSG00000117602.13 | RCAN3 | 1.09231 | 2.11E-14 |
| ENSG00000125384.7 | PTGER2 | 1.09023 | 2.93E-07 |
| ENSG00000162777.17 | DENND2D | 1.08909 | 7.19E-12 |
| ENSG00000167914.12 | GSDMA | 1.08846 | 2.07E-05 |
| ENSG00000167315.18 | ACAA2 | 1.08728 | 2.35E-18 |
| ENSG00000066185.13 | ZMYND12 | 1.08620 | 9.35E-11 |
| ENSG00000174567.8 | GOLT1A | 1.08258 | 7.64E-07 |
| ENSG00000105971.15 | CAV2 | 1.08243 | 3.28E-11 |
| ENSG00000165140.11 | FBP1 | 1.08116 | 5.61E-11 |
| ENSG00000124159.15 | MATN4 | 1.07672 | 1.23E-05 |
| ENSG00000125618.17 | PAX8 | 1.07406 | 2.73E-07 |
| ENSG00000122861.16 | PLAU | 1.06127 | 3.07E-06 |
| ENSG00000133135.14 | RNF128 | 1.06098 | 8.86E-05 |
| ENSG00000161638.11 | ITGA5 | 1.05883 | 1.06E-10 |
| ENSG00000110492.15 | MDK | 1.05810 | 1.20E-08 |
| ENSG00000134333.14 | LDHA | 1.04929 | 3.47E-11 |
| ENSG00000141574.8 | SECTM1 | 1.04812 | 4.58E-08 |
| ENSG00000180043.11 | FAM71E2 | 1.04686 | 2.89E-02 |

| | | | |
|---|---|---|---|
| ENSG00000144191.12 | CNGA3 | 1.04261 | 5.28E-06 |
| ENSG00000134363.12 | FST | 1.04129 | 2.87E-07 |
| ENSG00000197956.10 | S100A6 | 1.03950 | 7.89E-12 |
| ENSG00000146013.11 | GFRA3 | 1.03882 | 4.47E-06 |
| ENSG00000173662.21 | TAS1R1 | 1.03307 | 6.36E-12 |
| ENSG00000117215.15 | PLA2G2D | 1.03031 | 3.13E-06 |
| ENSG00000142552.8 | RCN3 | 1.02969 | 6.08E-11 |
| ENSG00000111321.11 | LTBR | 1.02713 | 2.22E-17 |
| ENSG00000197561.7 | ELANE | 1.01854 | 2.56E-06 |
| ENSG00000177697.19 | CD151 | 1.01750 | 2.13E-16 |
| ENSG00000171631.15 | P2RY6 | 1.01605 | 3.35E-09 |
| ENSG00000087086.15 | FTL | 1.01536 | 2.25E-13 |
| ENSG00000158716.9 | DUSP23 | 1.01434 | 6.65E-14 |
| ENSG00000205863.11 | C1QTNF9B | 1.01417 | 3.10E-06 |
| ENSG00000090339.9 | ICAM1 | 1.01412 | 1.29E-07 |
| ENSG00000134222.16 | PSRC1 | 1.01139 | 5.61E-11 |
| ENSG00000164136.17 | IL15 | 1.01079 | 3.36E-08 |
| ENSG00000111335.13 | OAS2 | 1.00855 | 3.36E-07 |
| ENSG00000171872.5 | KLF17 | 1.00556 | 2.67E-06 |
| ENSG00000132386.11 | SERPINF1 | 1.00329 | 5.02E-09 |
| ENSG00000124256.15 | ZBP1 | 1.00165 | 6.91E-06 |
| ENSG00000172183.15 | ISG20 | 1.00094 | 6.60E-10 |

| Table S6. Summary of disease associtaion of genes downregulated in HGG and present in | | | | | |
|---|---|---|---|---|---|
| **Ensembl gene ID** | **Gene name** | **Disease Associtation** | **Cancer Association** | **With a putative or validated tumor suppressor function** | **References** |
| ENSG00000125845.7 | BMP2 | https://www.disgenet.org/browser/1/1/0/650/ | Y | Y | https://www.ncbi.nlm.nih.gov/pmc/articles/PMC6825014/ |
| ENSG00000008300.17 | CELSR3 | https://www.disgenet.org/browser/1/1/0/1951/ | - | | |
| ENSG00000168993.15 | CPLX1 | https://www.disgenet.org/browser/1/1/0/10815/ | - | | |
| ENSG00000171587.15 | DSCAM | https://www.disgenet.org/browser/1/1/0/1826/ | - | | |
| ENSG00000129682.16 | FGF13 | https://www.disgenet.org/browser/1/1/0/2258/ | Y | Y | https://pubmed.ncbi.nlm.nih.gov/18930044/ |
| ENSG00000166206.15 | GABRB3 | https://www.disgenet.org/browser/1/1/0/2562/ | - | | |
| ENSG00000128683.14 | GAD1 | https://www.disgenet.org/browser/1/1/0/2571/ | - | | |
| ENSG00000136750.13 | GAD2 | https://www.disgenet.org/browser/1/1/0/2572/ | Y | | |
| ENSG00000168621.15 | GDNF | https://www.disgenet.org/browser/1/1/0/2668/ | Y | | |
| ENSG00000101076.17 | HNF4A | https://www.disgenet.org/browser/1/1/0/3172/ | Y | | |
| ENSG00000092051.17 | JPH4 | https://www.disgenet.org/browser/1/1/0/84502/ | Y | Y | https://pubmed.ncbi.nlm.nih.gov/19065659/ |
| ENSG00000187486.5 | KCNJ11 | https://www.disgenet.org/browser/1/1/0/3767/ | Y | | |
| ENSG00000092054.13 | MYH7 | https://www.disgenet.org/browser/1/1/0/4625/ | - | | |
| ENSG00000129152.4 | MYOD1 | https://www.disgenet.org/browser/1/1/0/4654/ | Y | Y | https://pubmed.ncbi.nlm.nih.gov/14767572/ |
| ENSG00000079841.18 | RIMS1 | https://www.disgenet.org/browser/1/1/0/22999/ | Y | | |
| ENSG00000198626.17 | RYR2 | https://www.disgenet.org/browser/1/1/0/6262/ | Y | Y | http://www.ncbi.nlm.nih.gov/pubmed/31135957 |
| ENSG00000261678.3 | SCRT1 | https://www.disgenet.org/browser/1/1/0/83482/ | Y | | |
| ENSG00000124140.14 | SLC12A5 | https://www.disgenet.org/browser/1/1/0/57468/ | Y | Y | https://pubmed.ncbi.nlm.nih.gov/21911617/ |
| ENSG00000187122.17 | SLIT1 | https://www.disgenet.org/browser/1/1/0/6585/ | Y | Y | https://www.ncbi.nlm.nih.gov/pmc/articles/PMC2409788/ |
| ENSG00000019505.8 | SYT13 | https://www.disgenet.org/browser/1/1/0/57586/ | Y | Y | https://pubmed.ncbi.nlm.nih.gov/20840848/ |
| ENSG00000139973.16 | SYT16 | https://www.disgenet.org/browser/1/1/0/83851/ | Y | Y | https://www.sciencedirect.com/science/article/abs/pii/S1567576919326220 |
| ENSG00000143858.12 | SYT2 | https://www.disgenet.org/browser/1/1/0/127833/ | - | | |
| ENSG00000130477.15 | UNC13A | https://www.disgenet.org/browser/1/1/0/23025/ | - | | |
| ENSG00000113763.12 | UNC5A | https://www.disgenet.org/browser/1/1/1/90249/ | Y | Y | https://pubmed.ncbi.nlm.nih.gov/12655055/ |
| ENSG00000188064.10 | WNT7B | https://www.disgenet.org/browser/1/1/0/7477/ | Y | Y | https://www.ncbi.nlm.nih.gov/pmc/articles/PMC5768065/ |
| ENSG00000109906.14 | ZBTB16 | https://www.disgenet.org/browser/1/1/0/7704/ | Y | | |

**Table S7. SKCM patients from the TCGA cohort.**

| Patient id | Gender | Age | Tumor type | Pathological stage (T) | Pathological stage (N) | Pathological stage (M) | Pathological stage | PFI status | PFI time | SRY expression |
|---|---|---|---|---|---|---|---|---|---|---|
| TCGA-BF-A1PX | Male | 56 | Primary | T4 | N2 | M0 | III | 0 | 282 | 0.47270 |
| TCGA-BF-A1Q0 | Male | 80 | Primary | T4 | N0 | M0 | II | 0 | 831 | 0.00000 |
| TCGA-BF-A3DM | Male | 63 | Primary | T2 | N0 | M0 | II | 0 | 601 | 0.00000 |
| TCGA-BF-A5EO | Male | 65 | Primary | T4 | N0 | M0 | II | 0 | 703 | 0.00000 |
| TCGA-BF-A5EQ | Male | 63 | Primary | T4 | N0 | M0 | II | 0 | 323 | 0.50450 |
| TCGA-BF-A5ER | Male | 63 | Primary | T4 | N0 | M0 | II | 0 | 327 | 0.00000 |
| TCGA-BF-A9VF | Male | 77 | Primary | T4 | N0 | M0 | II | 0 | 440 | 0.00000 |
| TCGA-BF-AAOX | Male | 83 | Primary | T4 | N0 | M0 | II | 0 | 444 | 0.00000 |
| TCGA-BF-AAP1 | Male | 86 | Primary | T4 | N0 | M0 | II | 0 | 409 | 0.00000 |
| TCGA-BF-AAP2 | Male | 62 | Primary | T3 | N0 | M0 | II | 0 | 405 | 0.13530 |
| TCGA-BF-AAP4 | Male | 61 | Primary | T4 | N0 | M0 | II | 0 | 335 | 0.00000 |
| TCGA-BF-AAP6 | Male | 55 | Primary | T4 | N2 | M0 | III | 0 | 325 | 0.07560 |
| TCGA-BF-AAP8 | Male | 58 | Primary | T4 | N0 | M0 | II | 0 | 447 | 0.00000 |
| TCGA-D3-A5GT | Male | 43 | Primary | T2 | N3 | M0 | III | 1 | 291 | 0.00000 |
| TCGA-D9-A3Z4 | Male | 54 | Primary | T4 | N3 | M0 | III | 1 | 192 | 0.07900 |
| TCGA-D9-A4Z2 | Male | 50 | Primary | T4 | N3 | M0 | III | 1 | 80 | 2.73480 |
| TCGA-D9-A4Z5 | Male | 68 | Primary | T4 | N0 | M0 | II | 0 | 218 | 0.00000 |
| TCGA-DA-A960 | Male | 73 | Primary | T3 | N0 | M0 | II | 0 | 804 | 0.05620 |
| TCGA-EB-A1NK | Male | 48 | Primary | T4 | N0 | M0 | II | 0 | 1039 | 0.22650 |
| TCGA-EB-A24C | Male | 56 | Primary | T4 | NX | M0 | [Not available] | 1 | 465 | 0.43620 |
| TCGA-EB-A24D | Male | 72 | Primary | T4 | N2 | M0 | III | 1 | 500 | 0.17130 |
| TCGA-EB-A299 | Male | 63 | Primary | T2 | N0 | M0 | II | 0 | 378 | 1.75940 |
| TCGA-EB-A3HV | Male | 37 | Primary | T4 | N0 | M0 | II | 0 | 39 | 0.59960 |
| TCGA-EB-A3XB | Male | 63 | Primary | T4 | NX | M0 | II | 0 | 796 | 0.06530 |
| TCGA-EB-A3XC | Male | 74 | Primary | T4 | N0 | M0 | II | 1 | 74 | 0.55490 |
| TCGA-EB-A3XF | Male | 57 | Primary | T4 | N0 | M0 | II | 0 | 278 | 0.00000 |
| TCGA-EB-A41A | Male | 90 | Primary | T4 | N0 | M0 | II | 0 | 262 | 0.04490 |
| TCGA-EB-A42Z | Male | 49 | Primary | T4 | N1 | M0 | III | 0 | 441 | 0.06440 |
| TCGA-EB-A430 | Male | 83 | Primary | T4 | N0 | M0 | II | 0 | 0 | 0.00000 |
| TCGA-EB-A431 | Male | 34 | Primary | T4 | N0 | M0 | II | 0 | 568 | 0.00000 |
| TCGA-EB-A44N | Male | 59 | Primary | T4 | N0 | M0 | II | 1 | 115 | 3.22260 |
| TCGA-EB-A44O | Male | 69 | Primary | T4 | N0 | M0 | II | 1 | 60 | 0.00000 |
| TCGA-EB-A4IS | Male | 77 | Primary | T3 | NX | M0 | II | 0 | 774 | 0.20160 |
| TCGA-EB-A4P0 | Male | 82 | Primary | T4 | N0 | M0 | II | 0 | 326 | 0.00000 |
| TCGA-EB-A51B | Male | 53 | Primary | T4 | NX | M0 | II | 0 | 931 | 0.00000 |

| | | | | | | | | | | |
|---|---|---|---|---|---|---|---|---|---|---|
| TCGA-EB-A553 | Male | 62 | Primary | T4 | N0 | M0 | II | 0 | 226 | 0.00000 |
| TCGA-EB-A57M | Male | 56 | Primary | T4 | N1 | M0 | III | 0 | 472 | 0.64910 |
| TCGA-EB-A5SE | Male | 73 | Primary | T3 | NX | M0 | II | 1 | 367 | 0.16800 |
| TCGA-EB-A5VU | Male | 56 | Primary | T4 | N1 | M0 | III | 1 | 217 | 0.82450 |
| TCGA-EB-A6QY | Male | 71 | Primary | T4 | N0 | M0 | II | 1 | 36 | 1.41670 |
| TCGA-EB-A85I | Male | 66 | Primary | T4 | N0 | M0 | II | 0 | 362 | 0.07000 |
| TCGA-EB-A97M | Male | 66 | Primary | T4 | N0 | M0 | II | 0 | 414 | 0.00000 |
| TCGA-ER-A194 | Male | 77 | Primary | [Not available] | N0 | M0 | [Not available] | 1 | 1354 | 0.04930 |
| TCGA-ER-A19T | Male | 51 | Primary | T4 | N3 | M1 | IV | 1 | 118 | 0.09380 |
| TCGA-ER-A2NB | Male | 57 | Primary | T4 | N2 | M0 | III | 1 | 461 | 0.07830 |
| TCGA-ER-A2NF | Male | 53 | Primary | T3 | N3 | M0 | III | 1 | 230 | 0.14440 |
| TCGA-ER-A42H | Male | 76 | Primary | [Not available] | [Not available] | [Not available] | [Not available] | 1 | 259 | 0.00000 |
| TCGA-FR-A3R1 | Male | 69 | Primary | T4 | N0 | M0 | II | 1 | 678 | 0.00000 |
| TCGA-FR-A726 | Male | 90 | Primary | T4 | N0 | M0 | II | 1 | 273 | 0.06010 |
| TCGA-FS-A1ZN | Male | 43 | Primary | T4 | N1 | M0 | III | 1 | 361 | 0.27730 |
| TCGA-FW-A5DX | Male | 71 | Primary | T4 | N3 | [Not available] | III | 0 | 640 | 0.00000 |
| TCGA-GF-A2C7 | Male | 48 | Primary | T4 | N0 | M0 | II | 0 | 21 | 0.17850 |
| TCGA-GF-A769 | Male | 39 | Primary | T4 | NX | M0 | II | 1 | 1070 | 0.00000 |
| TCGA-GN-A263 | Male | 24 | Primary | T4 | N3 | M1 | IV | 1 | 39 | 0.19440 |
| TCGA-GN-A26C | Male | 77 | Primary | T4 | N2 | M0 | III | 1 | 593 | 0.00000 |
| TCGA-GN-A8LN | Male | 68 | Primary | T4 | NX | M0 | II | 1 | 186 | 0.09410 |
| TCGA-IH-A3EA | Male | 61 | Primary | T4 | N0 | M0 | II | 1 | 314 | 0.00000 |
| TCGA-WE-A8K4 | Male | 85 | Primary | T4 | NX | M0 | II | 0 | 614 | 0.00000 |
| TCGA-XV-A9W2 | Male | 81 | Primary | T1 | N0 | M0 | I | 0 | 417 | 0.46330 |
| TCGA-XV-A9W5 | Male | 51 | Primary | T2 | N0 | M0 | I/II | 0 | 392 | 0.00000 |
| TCGA-YG-AA3N | Male | 67 | Primary | T4 | N0 | M0 | II | 0 | 306 | 0.34870 |

**Table S8. SRY target genes differentially expressed in SKCM samples presenting higher versus lower SRY expression.**

| Up-regulated SRY target genes | | | | Down-regulated SRY target genes | | | |
|---|---|---|---|---|---|---|---|
| gene.id | gene.symbol | log2FC | FDR | gene.id | gene.symbol | log2FC | FDR |
| ENSG00000084110.11 | HAL | 6.15889 | 8.19E-27 | ENSG00000184524.6 | CEND1 | -3.57983 | 5.18E-07 |
| ENSG00000126233.2 | SLURP1 | 6.06521 | 1.85E-09 | ENSG00000128564.7 | VGF | -3.28182 | 5.29E-05 |
| ENSG00000159337.7 | PLA2G4D | 6.04358 | 1.45E-16 | ENSG00000160282.14 | FTCD | -3.09140 | 2.01E-05 |
| ENSG00000162040.7 | HS3ST6 | 5.37205 | 7.20E-12 | ENSG00000130226.17 | DPP6 | -2.68652 | 4.26E-03 |
| ENSG00000139648.7 | KRT71 | 5.04137 | 1.71E-02 | ENSG00000152092.16 | ASTN1 | -2.43589 | 4.25E-02 |
| ENSG00000167768.4 | KRT1 | 4.92854 | 3.27E-05 | ENSG00000156222.12 | SLC28A1 | -2.32539 | 4.78E-03 |
| ENSG00000170426.2 | SDR9C7 | 4.88782 | 3.90E-09 | ENSG00000141744.4 | PNMT | -2.20153 | 1.40E-02 |
| ENSG00000169035.12 | KLK7 | 4.82332 | 1.50E-06 | ENSG00000073670.14 | ADAM11 | -2.19384 | 4.35E-04 |
| ENSG00000205076.4 | LGALS7 | 4.79335 | 8.35E-06 | ENSG00000173662.21 | TAS1R1 | -2.17513 | 3.44E-03 |
| ENSG00000094796.5 | KRT31 | 4.71539 | 6.18E-08 | ENSG00000123999.5 | INHA | -2.08443 | 1.37E-04 |
| ENSG00000167741.11 | GGT6 | 4.63555 | 8.73E-11 | ENSG00000116176.6 | TPSG1 | -1.97301 | 1.33E-02 |
| ENSG00000180316.12 | PNPLA1 | 4.60789 | 5.48E-20 | ENSG00000101188.5 | NTSR1 | -1.88947 | 1.02E-02 |
| ENSG00000168447.11 | SCNN1B | 4.55201 | 1.11E-17 | ENSG00000179111.9 | HES7 | -1.84638 | 5.63E-03 |
| ENSG00000137709.10 | POU2F3 | 4.51016 | 3.26E-12 | ENSG00000123901.9 | GPR83 | -1.84492 | 1.12E-02 |
| ENSG00000188624.3 | IGFL3 | 4.37395 | 7.59E-08 | ENSG00000066923.17 | STAG3 | -1.82944 | 3.67E-04 |
| ENSG00000167914.12 | GSDMA | 4.35864 | 6.05E-16 | ENSG00000168993.15 | CPLX1 | -1.80072 | 4.20E-03 |
| ENSG00000137699.17 | TRIM29 | 4.31821 | 1.39E-07 | ENSG00000129673.10 | AANAT | -1.77969 | 5.41E-04 |
| ENSG00000109182.12 | CWH43 | 4.29055 | 9.79E-06 | ENSG00000105048.17 | TNNT1 | -1.69728 | 4.43E-02 |
| ENSG00000016402.13 | IL20RA | 4.28818 | 2.44E-17 | ENSG00000064300.9 | NGFR | -1.67668 | 4.16E-02 |
| ENSG00000109101.8 | FOXN1 | 4.06128 | 2.08E-05 | ENSG00000008300.17 | CELSR3 | -1.66349 | 1.41E-04 |
| ENSG00000111319.13 | SCNN1A | 4.05949 | 2.25E-17 | ENSG00000154678.18 | PDE1C | -1.65970 | 3.74E-02 |
| ENSG00000132698.15 | RAB25 | 4.02920 | 9.04E-08 | ENSG00000135406.14 | PRPH | -1.65122 | 8.79E-03 |
| ENSG00000129455.15 | KLK8 | 4.01667 | 1.01E-04 | ENSG00000198576.4 | ARC | -1.63498 | 1.94E-02 |
| ENSG00000167757.14 | KLK11 | 4.00795 | 2.19E-05 | ENSG00000125878.6 | TCF15 | -1.62713 | 2.24E-02 |
| ENSG00000100170.10 | SLC5A1 | 3.99512 | 4.25E-10 | ENSG00000197587.11 | DMBX1 | -1.62292 | 3.55E-02 |
| ENSG00000155066.16 | PROM2 | 3.79794 | 5.13E-10 | ENSG00000156113.24 | KCNMA1 | -1.58899 | 1.74E-02 |
| ENSG00000006555.11 | TTC22 | 3.76135 | 2.80E-11 | ENSG00000172183.15 | ISG20 | -1.58309 | 3.94E-03 |
| ENSG00000186832.9 | KRT16 | 3.70787 | 1.20E-03 | ENSG00000178773.15 | CPNE7 | -1.56043 | 3.55E-02 |

| | | | | | | | |
|---|---|---|---|---|---|---|---|
| ENSG00000167880.8 | EVPL | 3.70000 | 9.50E-08 | ENSG00000186897.5 | C1QL4 | -1.55970 | 1.63E-02 |
| ENSG00000124466.9 | LYPD3 | 3.69799 | 7.07E-10 | ENSG00000171596.7 | NMUR1 | -1.53427 | 3.07E-02 |
| ENSG00000074211.14 | PPP2R2C | 3.62818 | 5.92E-08 | ENSG00000105376.5 | ICAM5 | -1.44195 | 1.87E-02 |
| ENSG00000178934.5 | LGALS7B | 3.60432 | 2.23E-04 | ENSG00000178531.6 | CTXN1 | -1.41247 | 9.20E-03 |
| ENSG00000206075.14 | SERPINB5 | 3.59754 | 4.35E-04 | ENSG00000242252.2 | BGLAP | -1.39631 | 1.32E-02 |
| ENSG00000140254.12 | DUOXA1 | 3.58792 | 3.11E-09 | ENSG00000105464.3 | GRIN2D | -1.38344 | 2.15E-02 |
| ENSG00000174502.19 | SLC26A9 | 3.55548 | 1.29E-10 | ENSG00000103269.14 | RHBDL1 | -1.34370 | 8.31E-04 |
| ENSG00000179178.11 | TMEM125 | 3.47047 | 1.68E-10 | ENSG00000181409.14 | AATK | -1.33135 | 1.53E-02 |
| ENSG00000166736.11 | HTR3A | 3.36909 | 1.20E-04 | ENSG00000167874.7 | TMEM88 | -1.33124 | 5.17E-04 |
| ENSG00000169583.13 | CLIC3 | 3.36545 | 1.22E-10 | ENSG00000144485.11 | HES6 | -1.29025 | 1.51E-02 |
| ENSG00000261150.3 | EPPK1 | 3.29730 | 1.79E-06 | ENSG00000104826.15 | LHB | -1.28598 | 3.32E-02 |
| ENSG00000183347.15 | GBP6 | 3.24154 | 8.06E-06 | ENSG00000130598.16 | TNNI2 | -1.22260 | 4.85E-02 |
| ENSG00000175793.12 | SFN | 3.20586 | 5.99E-04 | ENSG00000139540.12 | SLC39A5 | -1.18848 | 2.68E-02 |
| ENSG00000129451.12 | KLK10 | 3.17750 | 1.32E-04 | ENSG00000106366.9 | SERPINE1 | -1.15347 | 3.49E-02 |
| ENSG00000131746.13 | TNS4 | 3.10463 | 1.53E-05 | ENSG00000179085.8 | DPM3 | -1.08359 | 1.74E-03 |
| ENSG00000189334.9 | S100A14 | 3.02989 | 6.07E-04 | ENSG00000102878.17 | HSF4 | -1.07267 | 1.55E-02 |
| ENSG00000170454.6 | KRT75 | 3.01825 | 8.92E-03 | ENSG00000170835.16 | CEL | -1.07165 | 2.14E-02 |
| ENSG00000147689.16 | FAM83A | 3.01526 | 1.87E-03 | ENSG00000169750.9 | RAC3 | -1.03986 | 4.81E-02 |
| ENSG00000189280.3 | GJB5 | 3.00106 | 1.92E-04 | ENSG00000173991.6 | TCAP | -1.01185 | 2.01E-02 |
| ENSG00000189143.9 | CLDN4 | 2.99221 | 6.40E-10 | | | | |
| ENSG00000184363.10 | PKP3 | 2.96779 | 1.83E-04 | | | | |
| ENSG00000189433.7 | GJB4 | 2.95099 | 3.10E-03 | | | | |
| ENSG00000090512.12 | FETUB | 2.88309 | 1.39E-03 | | | | |
| ENSG00000128422.17 | KRT17 | 2.82619 | 3.37E-03 | | | | |
| ENSG00000170054.16 | SERPINA9 | 2.79908 | 5.61E-03 | | | | |
| ENSG00000149043.16 | SYT8 | 2.77053 | 3.47E-06 | | | | |
| ENSG00000150556.17 | LYPD6B | 2.75587 | 3.94E-06 | | | | |
| ENSG00000188064.10 | WNT7B | 2.66025 | 4.33E-05 | | | | |
| ENSG00000142623.11 | PADI1 | 2.62448 | 1.38E-03 | | | | |
| ENSG00000173338.13 | KCNK7 | 2.61516 | 1.89E-10 | | | | |
| ENSG00000104892.17 | KLC3 | 2.60382 | 3.98E-05 | | | | |

| | | | |
|---|---|---|---|
| ENSG00000171346.16 | KRT15 | 2.51228 | 9.55E-03 |
| ENSG00000156711.17 | MAPK13 | 2.38424 | 2.87E-13 |
| ENSG00000129194.8 | SOX15 | 2.30133 | 7.81E-07 |
| ENSG00000161249.21 | DMKN | 2.30006 | 3.02E-04 |
| ENSG00000170006.12 | TMEM154 | 2.29678 | 1.28E-05 |
| ENSG00000156738.18 | MS4A1 | 2.29364 | 4.04E-03 |
| ENSG00000172554.12 | SNTG2 | 2.26575 | 3.87E-05 |
| ENSG00000133661.17 | SFTPD | 2.24306 | 6.78E-05 |
| ENSG00000174564.13 | IL20RB | 2.22867 | 5.33E-04 |
| ENSG00000043039.7 | BARX2 | 2.22645 | 2.24E-03 |
| ENSG00000134873.10 | CLDN10 | 2.15084 | 3.94E-04 |
| ENSG00000179023.8 | KLHDC7A | 2.14520 | 2.82E-02 |
| ENSG00000146648.19 | EGFR | 2.06709 | 1.19E-04 |
| ENSG00000050628.20 | PTGER3 | 2.05600 | 4.15E-09 |
| ENSG00000139292.13 | LGR5 | 2.05212 | 4.18E-04 |
| ENSG00000196092.14 | PAX5 | 2.02230 | 8.55E-04 |
| ENSG00000104881.16 | PPP1R13L | 2.01098 | 6.35E-08 |
| ENSG00000156463.18 | SH3RF2 | 1.92874 | 4.08E-02 |
| ENSG00000115353.11 | TACR1 | 1.92602 | 2.87E-04 |
| ENSG00000167968.13 | DNASE1L2 | 1.91117 | 4.86E-04 |
| ENSG00000140287.11 | HDC | 1.89800 | 2.39E-03 |
| ENSG00000064692.20 | SNCAIP | 1.88551 | 3.39E-05 |
| ENSG00000168081.9 | PNOC | 1.88437 | 1.67E-02 |
| ENSG00000105664.11 | COMP | 1.85259 | 2.19E-02 |
| ENSG00000111405.9 | ENDOU | 1.84180 | 4.27E-02 |
| ENSG00000104435.14 | STMN2 | 1.83338 | 1.22E-03 |
| ENSG00000149654.11 | CDH22 | 1.82985 | 1.07E-02 |
| ENSG00000171056.8 | SOX7 | 1.82826 | 2.37E-05 |
| ENSG00000163435.16 | ELF3 | 1.82815 | 2.12E-03 |
| ENSG00000099337.5 | KCNK6 | 1.80542 | 2.79E-08 |
| ENSG00000223572.10 | CKMT1A | 1.80472 | 2.12E-02 |

| | | | |
|---|---|---|---|
| ENSG00000115884.11 | SDC1 | 1.76865 | 8.69E-05 |
| ENSG00000163472.19 | TMEM79 | 1.75594 | 1.15E-10 |
| ENSG00000163492.15 | CCDC141 | 1.73508 | 4.60E-02 |
| ENSG00000165124.18 | SVEP1 | 1.71672 | 3.13E-04 |
| ENSG00000085741.13 | WNT11 | 1.69664 | 1.82E-02 |
| ENSG00000114251.15 | WNT5A | 1.69360 | 1.34E-04 |
| ENSG00000177272.9 | KCNA3 | 1.66685 | 6.46E-03 |
| ENSG00000180875.5 | GREM2 | 1.66536 | 5.91E-03 |
| ENSG00000161270.20 | NPHS1 | 1.65821 | 4.60E-02 |
| ENSG00000167183.3 | PRR15L | 1.63474 | 1.25E-02 |
| ENSG00000078596.11 | ITM2A | 1.63358 | 9.27E-04 |
| ENSG00000103034.14 | NDRG4 | 1.60657 | 1.04E-02 |
| ENSG00000245848.3 | CEBPA | 1.59933 | 6.35E-05 |
| ENSG00000197406.7 | DIO3 | 1.59185 | 2.87E-02 |
| ENSG00000078549.15 | ADCYAP1R1 | 1.56337 | 2.04E-03 |
| ENSG00000186567.13 | CEACAM19 | 1.54252 | 1.44E-03 |
| ENSG00000169067.4 | ACTBL2 | 1.53500 | 2.42E-02 |
| ENSG00000172382.10 | PRSS27 | 1.50183 | 8.18E-04 |
| ENSG00000177191.2 | B3GNT8 | 1.49902 | 1.85E-03 |
| ENSG00000127152.18 | BCL11B | 1.47496 | 1.64E-02 |
| ENSG00000140832.10 | MARVELD3 | 1.44904 | 2.66E-02 |
| ENSG00000161958.11 | FGF11 | 1.39343 | 2.90E-02 |
| ENSG00000174348.14 | PODN | 1.37395 | 3.61E-03 |
| ENSG00000213903.9 | LTB4R | 1.34721 | 8.08E-04 |
| ENSG00000162383.13 | SLC1A7 | 1.34515 | 1.78E-02 |
| ENSG00000105289.15 | TJP3 | 1.28394 | 4.11E-02 |
| ENSG00000121858.11 | TNFSF10 | 1.21810 | 1.74E-02 |
| ENSG00000122025.15 | FLT3 | 1.21088 | 3.22E-02 |
| ENSG00000176641.11 | RNF152 | 1.20433 | 1.34E-03 |
| ENSG00000105641.4 | SLC5A5 | 1.20085 | 2.78E-02 |
| ENSG00000136943.11 | CTSV | 1.18177 | 2.16E-02 |

| | | | |
|---|---|---|---|
| ENSG00000174640.15 | SLCO2A1 | 1.13993 | 5.63E-03 |
| ENSG00000153823.19 | PID1 | 1.10481 | 7.91E-04 |
| ENSG00000172935.9 | MRGPRF | 1.08795 | 3.23E-02 |
| ENSG00000165646.14 | SLC18A2 | 1.07061 | 3.71E-03 |
| ENSG00000178175.12 | ZNF366 | 1.04833 | 9.67E-03 |
| ENSG00000151388.11 | ADAMTS12 | 1.02862 | 3.12E-02 |
| ENSG00000162493.16 | PDPN | 1.02715 | 4.25E-02 |
| ENSG00000163874.11 | ZC3H12A | 1.02069 | 1.74E-02 |
| ENSG00000108821.14 | COL1A1 | 1.00248 | 2.60E-02 |
